# Deep learning linking mechanistic models to single-cell transcriptomics data reveals transcriptional bursting in response to DNA damage

**DOI:** 10.1101/2024.07.10.602845

**Authors:** Zhiwei Huang, Songhao Luo, Zihao Wang, Zhenquan Zhang, Benyuan Jiang, Qing Nie, Jiajun Zhang

## Abstract

Cells must adopt flexible regulatory strategies to make decisions regarding their fate, including differentiation, apoptosis, or survival in the face of various external stimuli. One key cellular strategy that enables these functions is stochastic gene expression programs. However, understanding how transcriptional bursting, and consequently, cell fate, responds to DNA damage on a genome-wide scale poses a challenge. In this study, we propose an interpretable and scalable inference framework, DeepTX, that leverages deep learning methods to connect mechanistic models and scRNA-seq data, thereby revealing genome-wide transcriptional burst kinetics. This framework enables rapid and accurate solutions to transcription models and the inference of transcriptional burst kinetics from scRNA-seq data. Applying this framework to several scRNA-seq datasets of DNA-damaging drug treatments, we observed that fluctuations in transcriptional bursting induced by different drugs was associated with distinct fate decisions: IdU treatment was associated with differentiation in mouse embryonic stem cells by increasing the burst size of gene expression, while low- and high-dose 5FU treatments in human colon cancer cells were associated with changes in burst frequency that corresponded to apoptosis- and survival-related fate, respectively. Together, these results show that DeepTX enables genome-wide inference of transcriptional bursting from single-cell transcriptomics data and can generate hypotheses about how bursting dynamics relate to cell fate decisions.

## Introduction

Cells must employ flexible regulatory strategies to determine their fate, including cell differentiation, survival, or apoptosis, in response to diverse external stimuli (***López-Maury L et al., 2008; Stadhouders R et al., 2019***). One key cellular strategy that enables both beneficial and detrimental functions is the stochastic nature of gene expression programs, arising from intricate biochemical processes and termed “noise” (***Raj A & van Oudenaarden A, 2008; Balázsi G et al., 2011***; ***Losick R & Desplan C, 2008***). Noise can arise at many steps —from stochastic transcription factor binding and chromatin remodeling, to variability in RNA transcription and protein translation. Among these diverse sources, transcriptional noise sits at the very start of the central dogma, converting upstream molecular fluctuations into the mRNA outputs that drive all downstream regulatory and functional processes. Transcriptional noise originates from a discontinuous pattern, wherein mRNA production occurs through alternating active and inactive gene states ***(Eling N et al., 2019; Tunnacliffe E & Chubb JR, 2020***). This phenomenon, known as “transcriptional bursting”, has been observed in numerous experiments across both prokaryotic and eukaryotic cells (***Golding I et al., 2005; Chubb JR et al., 2006; Larson DR et al., 2011***). The burst kinetics, characterized by burst frequency and burst size, describe the stochastic behavior of molecular interactions at the gene expression process, offering valuable insights into how cells encode and leverage variability. However, it remains an unresolved question how cells make fate decisions in response to external signals through genome-wide regulation of transcriptional burst mechanisms (***Lammers NC et al., 2020; Rodriguez J & Larson DR, 2020)***.

DNA damage is a crucial and prevalent factor contributing to cellular responses, inducing varying degrees of cytotoxicity and thereby influencing diverse cell fate decisions ***(Krenning L et al., 2019; Su TT, 2006; Hafner A et al., 2019***). For instance, low cytotoxicity limits cell proliferation (***van den Berg J et al., 2019; Arora M et al., 2017; Barr AR et al., 2017; Deng Z et al., 2023***), whereas heightened cytotoxicity prompts cell differentiation and senescence (Müllers E et al., 2014; Feringa FM et al., 2018; Toledo LI et al., 2008; ***Zhao Y et al., 2023***). Furthermore, extreme scenarios of high cytotoxicity can cause cells to engage in the apoptotic process ***(Yousefzadeh M et al., 2021; Carneiro BA & El-Deiry WS, 2020; Roos WP & Kaina B, 2013; Zheng P et al., 2018***). Experiments have shown that DNA damage can disrupt the gene transcription process (***Lans H et al., 2019***). Specifically, DNA damage slows down the movement of RNA Pol II along the DNA strand (***Muñoz MJ et al., 2009***). When more severe DNA damage is encountered, RNA Pol II exhibits a sliding pause behavior for DNA repair before resuming transcription (***Figure 1A***) (***Gregersen LH & Svejstrup JQ, 2018; Geijer ME & Marteijn JA, 2018; Giono LE et al., 2016***). These changes in RNA Pol II movement behavior would further reflect the cellular regulation of gene expression, particularly burst kinetics ***(Friedrich D et al., 2019; Calia GP et al., 2023***). A recent study has shown that cellular responses to DNA damage involve the regulation of burst frequency (BF) on specific genes, rather than a genome-wide conclusion (***Friedrich D et al., 2019***). They have demonstrated that cells respond to DNA damage by increasing gene expression noise, but they do not delve into the topic of transcriptional bursts (***Calia GP et al., 2023; Desai RV et al., 2021***). Therefore, the question of how DNA damage causes cells to regulate transcriptional burst kinetics on a genome-wide scale remains unresolved.

**Figure 1.**
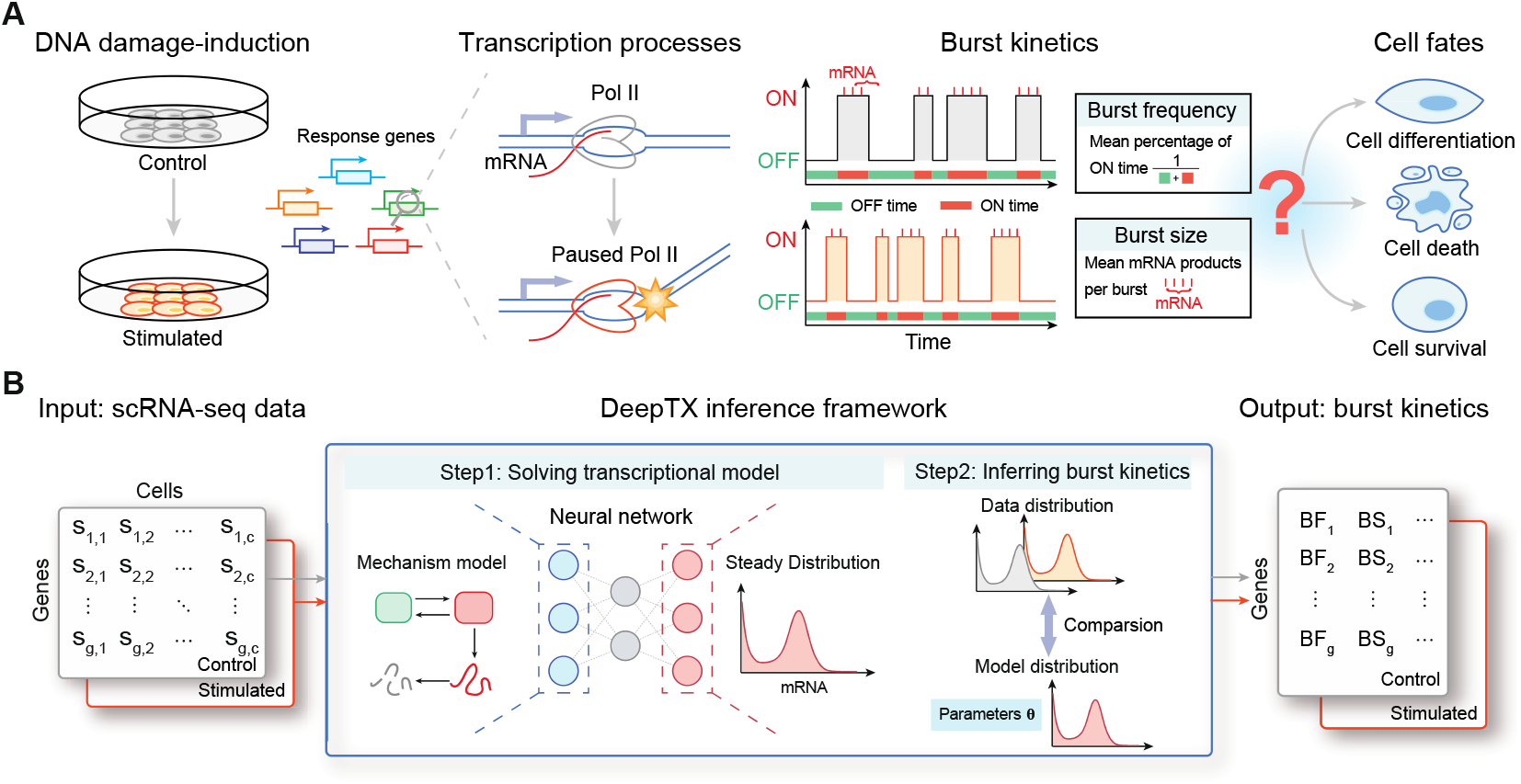
Overview of DeepTX framework. **(A)** Cells stimulated by DNA damaging drugs cause damage to the DNA double strands, which will slow down or even stop the movement of RNA Pol II on the DNA double strands. Changes in the state of RNA Pol II movement will lead to changes in the kinetics of gene expression bursts (including burst frequency and burst size), which will affect cell fate decisions such as apoptosis, differentiation, and survival. **(B)** The input to the DeepTX framework is scRNA-seq data, and the output is the burst kinetics corresponding to each gene. The core of the DeepTX inference framework is a hierarchical model, which is a mixture of the stationary distribution of the mechanism model solved by the neural network (Step 1) and the binomial distribution followed by the sequencing process. This hierarchical model is used to infer the out dynamics parameters corresponding to the data distribution (Step 2).

To understand the dynamics of transcriptional bursts, ideally, one needs to observe the fluctuations of gene expression over a continuous time interval, such as smFISH, scNT-seq, and HT-smFISH (***Femino AM et al., 1998; Qiu Q et al., 2020; Safieddine A et al., 2023)***. However, most existing techniques can only observe a limited number of genes, making their application to genome-wide studies challenging. Fortunately, single-cell RNA sequencing (scRNA-seq) technology provides excellent opportunities to explore this question, emerging as the leading method for genome-wide mRNA measurements and revealing gene expression noise within individual cells ***(Tanay A & Regev A, 2017; Picelli S et al., 2013; Zheng GX et al., 2017***). Many studies have employed scRNA-seq to elucidate the diverse sources of gene expression noise and the fundamental principles of transcriptional dynamics on a genome-wide scale ***(Eling N et al., 2019; Faure AJ et al., 2017; Morgan MD & Marioni JC, 2018; Ochiai H et al ., 2020)***. However, scRNA-seq data provide only static snapshots of cellular states. To infer the underlying transcriptional burst kinetics, one can apply mathematical models of stochastic gene expression, typically under the assumption that the observed distributions of scRNA-seq data reflect the steady-state outcome of the underlying dynamic processes ***(Larsson AJM et al., 2019; Luo S et al., 2023; Luo S et al., 2023***).

For mathematical modeling, the gene expression model for DNA damage ought to encompass more sophisticated generalized properties compared to the traditional telegraph model, which is a common model describing gene expression burst kinetics by genes switching stochastically between active (ON) and inactive (OFF) states with single-step process (exponentially distributed waiting times) (***Stumpf PS et al., 2017; Voss TC & Hager GL, 2014***). However, the presence of DNA damage necessitates modeling the transcriptional process as a multi-step process, rather than a single-step process, to capture the additional complexity introduced by the damage (***Singh A et al., 2013; Cavallaro M et al., 2021***). Many efforts in this field are addressing the challenges of describing this multi-step process of gene expression in an interpretable and tractable model. One common modeling approach is to introduce multiple intermediate states between inactivate and activate states (***Zhang J et al., 2012; Zhang J & Zhou T, 2014; Zhou T & Zhang J, 2012***). Although this multi-state model can fit the experimental data better, it is hard to rationalize the state numbers and parameters (***Desai RV et al., 2021; Rodriguez J et al., 2019; Zoller B et al., 2015***). Another alternative modeling approach is direct non-Markovian modeling for non-exponential waiting times in gene state, which maps the multiple parameters from multi-state models to a small number of interpretable parameters that are easily observed experimentally (***Daigle BJ, Jr. et al., 2015; Kumar N et al., 2015; Schwabe A et al., 2012; Stinchcombe AR et al., 2012; Zhang J & Zhou T, 2019; Zhang JJ & Zhou TS, 2019***). However, the ensuing difficulty is solving the analytic solution of the steady distribution from a non-Markovian model, which may require some numerical simulation methods that are computationally resource-intensive and time-consuming, such as Monte Carlo methods (***Sisson SA et al., 2007***).

Statistical inference, particularly in recovering burst kinetic parameters from genome-wide scRNA-seq data, necessitates efficient and scalable inference algorithms (***Gómez-Schiavon M et al., 2017***). Therefore, the stochastic dynamical system must be quickly solved as parameters are continuously updated throughout the inference process. Deep learning exhibits a broad array of applications as a contemporary method for addressing complex systems (***Jiang Q et al., 2021; Wang S et al., 2019; Michoski C et al., 2020***). For example, some studies employed neural networks to establish the mapping relationship between a wide range of parameters in the gene expression model and the corresponding pre-simulated steady distribution (***Figure 2B***) (***Jiang Q et al., 2021; Wang S et al., 2019; Davis CN et al., 2020; Tang Y et al., 2023***). This approach, based on neural networks, has been utilized for parameter inference in both deterministic and stochastic models (***Jiang Q et al., 2021; Gaskin T et al., 2023; Sukys A et al., 2022; Tang W et al., 2023)***.

**Figure 2.**
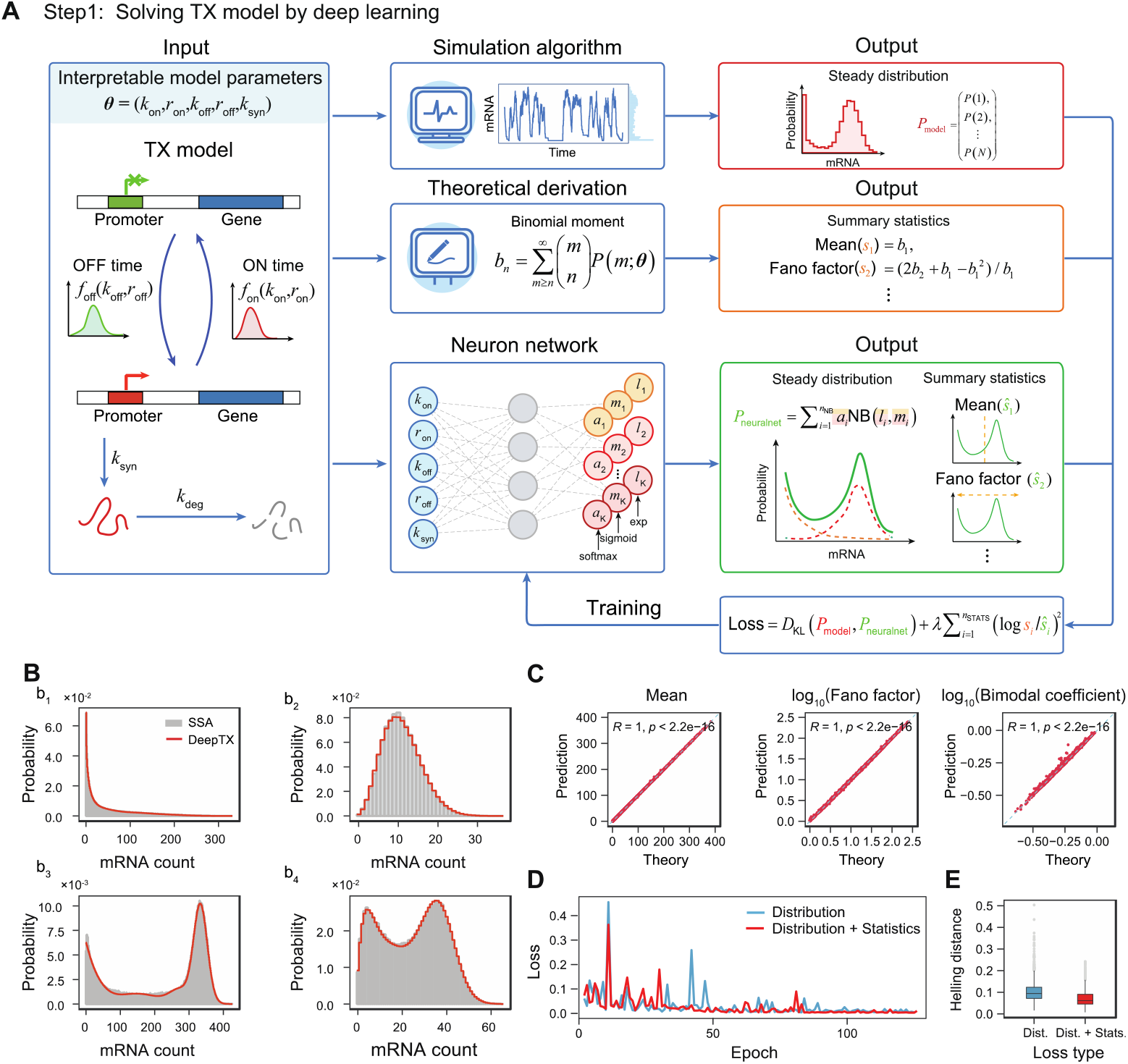
Solving TXmodel using deep learning. **(A)** With the TXmodel and corresponding model parameters ***θ***, we obtain a steady distribution using the SSA simulation and the moment statistics employing binomial moment theory. These results are compared with the distribution and moment statistics of the neural network’s output to calculate the neural network’s loss value. The parameters of the neural network are optimized using gradient descent until convergence is achieved, indicating attainment of optimal parameters. **(B)** Comparison between SSA results and DeepTX prediction results for four sets of test parameters. The gray bars and the red stepped line correspond to the distributions obtained from SSA simulations and DeepTX predictions, respectively. **(C)** Verification results of the moment statistics predicted by DeepTX and the true moments on a test set containing 1000 elements. **(D)** The loss curve during DeepTX training. The blue curve represents that the loss function is composed of KL divergence of distributions, and the red curve represents that the loss function is composed of KL divergence of distributions and statistics. **(E)** Box plots of the Hellinger distance between the true distribution and the predicted distribution by DeepTX of different loss types on the test set.

In this study, we introduce DeepTX, a deep learning inference framework that integrates mechanistic models and deep learning methods to elucidate the genome-wide regulation of transcriptional bursts in DNA damage response using scRNA-seq data. The DeepTX framework comprises two modules: DeepTXsolver and DeepTXinferrer. Specifically, DeepTX initially employs the DeepTXsolver module to efficiently solve complex gene expression dynamics models using neural network architectures. It subsequently utilizes the DeepTXinferrer module to accurately infer potential transcriptional burst kinetic parameters using Bayesian methods. DeepTX demonstrates good performance on synthetic datasets. Furthermore, to investigate genome-wide regulation of transcriptional bursts in response to DNA damage, we applied the DeepTX framework to three DNA damage-related scRNA-seq datasets representing cell differentiation, apoptosis, and cell survival. As a result, we observed that the fate decision of mouse embryonic stem cells (mESCs) to undergo cell differentiation in response to DNA damage caused by 5’-iodo-2’-deoxyuridine (IdU) compounds was associated with enhanced burst size (BS enhancer) of genes linked to delayed mitosis phase transition, which may reflect regulatory programs related to reprogramming and differentiation. In colorectal cancer cells, low-dose 5-fluorouracil (5FU) treatment was associated with increased burst frequency (BF enhancer) of genes connected to intracellular oxidative stress, coinciding with apoptosis-related processes. By contrast, high-dose 5FU treatment was associated with BF enhancer genes enriched in telomerase-related pathways, which may mitigate oxidative stress and correlate with cellular resistance. In conclusion, DeepTX is a computational framework with utility and extensibility to infer transcriptional dynamics from scRNA-seq data.

## Results

### Overview of DeepTX framework

To understand how cell influences underlying transcriptional mechanisms in response to DNA damage, as described in the introduction section (***Figure 1A***), we propose an effective computational inference framework, referred to as DeepTX (***Figure 1B***). DeepTX takes as input a set of scRNA-seq datasets and then outputs genome-wide burst kinetics parameters with the benefit from deep learning methods (***Figure 1B***). The inference process of the DeepTX framework is composed of two crucial modules, DeepTXsolver and DeepTXinferrer. The first module, DeepTXsolver, is to solve for the stationary distribution of a gene expression dynamic model (***Figure 1B middle***). A neural network is constructed in DeepTXsolver to approximate the mapping from model parameters to the corresponding stationary distribution. As in many biological contexts, there is a need to model dynamics that describe more realistic gene expression processes that are difficult to solve. For instance, gene expression under DNA damage is better represented by multi-step transcriptional models, which more faithfully reflect the underlying regulatory complexity. Moreover, sequencing noise should be integrated into the modeling framework (see Methods). The second module, DeepTXinferrer, infers the burst kinetics parameter (***Figure 1B middle***). Using a trained neural network from DeepTXsolver, we can easily and quickly obtain the stationary distribution (called model distribution) of any parameter within a reasonable parameter space. DeepTXinferrer compares the model distribution to the stationary distribution of each gene in the scRNA-seq data (called data distribution) as the parameters are iterated until convergence and utilize the Bayesian inference to output the posterior distribution of burst kinetic parameter (***Figure 1B right)***. We will describe these two modules in detail in the following two sections.

### DeepTX solves the transcriptional model using deep learning

To enable the recovery of kinetic information from static scRNA-seq data, we first performed rational mechanistic modeling of gene expression processes with DNA damage. The gene transcription burst process is often characterized using the traditional two-state model (***Singh A et al., 2013; Cavallaro M et al., 2021***), which assumes that genes switch randomly between OFF and ON states with exponentially distributed waiting times for each state. However, the gene expression process is inherently a multi-step process, which particularly cannot be neglected under conditions of DNA damage. DNA damage can result in slowing or even stopping the RNA pol II movement (***Lans H et al., 2019***) and cause many macromolecules to be recruited for damage repair. This process will affect the spatially localized behavior of the promoter (***Lans H et al., 2019***), causing the dwell time of promoter inactivation and activation that cannot be approximated by a simple two state. We therefore used a generalized model we developed previously (***Luo S et al., 2023***), called TXmodel here, which extended the state waiting time to an arbitrary distribution, i.e., the gene expression system is a non-Markovian system (***Figure 2A and see Methods 1.2***). More specifically, the waiting times for the transitions between the OFF and ON states are represented by two random variables, assumed to follow the arbitrary distributions *f*_off_ (*k*_off_, *r*_off_ ) and *f*_on_ (*k*_on_, *r*_on_ ), respectively. Additionally, we assumed that the rates of mRNA synthesis and degradation are constants, i.e., the waiting times for mRNA transcription and degradation follow exponential distributions with rate parameters *k*_syn_ and *k*_deg_, respectively (***Figure 2A left***). Nevertheless, this non-Markovian gene expression system is difficult to solve stationary distribution analytically, while numerical solution methods are time-consuming.

For that reason, we utilized a deep learning approach, which has been demonstrated to be effective in solving stochastic systems (***Gupta A et al., 2021***), to construct a mapping from the parameter of the mechanistic TXmodel to its corresponding stationary distribution. More specifically, we aimed to train a fully connected neural network referred as DeepTXsolver, whose inputs are the parameters of the TXmodel ***θ*** and whose output is a stationary distribution parameterized by a mixed negative binomial distribution *P*_neuralnet_, which has good performance in approximating the solution of the chemical master equation (***Perez-Carrasco R et al., 2020; Öcal K et al., 2022***) (***Figure 2A, Supplementary Figure S1 and see Methods 1.3***). To generate effective training sets for DeepTXsolver, we first generated a large number of parameter sets within a reasonable range of parameter space by using Sobol sampling (see Methods 1.3) (***Sobol IM, 1967***). Subsequently, for each parameter set, we employed a modified stochastic simulation algorithm (SSA) for non-Markovian TXmodels to generate numerous samples (***Figure 2A and see Methods 1.3***). The resulting stationary distributions *P*_simulation_ served as training labels. Additionally, we demonstrated that the arbitrary order binomial moments of the TXmodel can be analytically solved iteratively within the context of queueing theory (***Zhang Z et al., 2021; Zhang J et al., 2024***). This enables the straightforward computation of significant summary statistics, including mean and Fano factor (***Figure 2A, see Methods 1.2, and Supplementary Text 1.2***). Thus, our loss function comprises two primary components: (i) the Kullback-Leibler (KL) divergence between the predicted distribution generated by the neural network *P*_neuralnet_ and the labeled distribution generated by the SSA *P*_simulation_, which is a widely used metric for quantifying the difference between two probability distributions; (ii) the logarithmic error of the predicted summary statistics computed by the neural network ***ŝ*** and the labeled summary statistics ***s*** computed by theoretical derivation (***Figure 2A and see Methods 1.3***). Furthermore, we compared various optimizers and hyperparameters of model architectures to determine the best predicting performance (***see Methods 1.3 and Supplementary Figure S2***).

Consequently, the loss function of DeepTXsolver converges effectively during the training process. We utilized DeepTXsolver to predict the stationary distribution using parameters from the test set. We observed that it effectively fits all four representative distributions from the TXmodel, encompassing unimodal distribution at zero point, unimodal distribution at non-zero point, bimodal distribution with one peak at zero point, and bimodal distribution with both peaks at non-zero point (***Figure 2B***). These distributions have been extensively linked to cell fate decisions in numerous experiments (***Gupta PB et al., 2011; Cohen AA et al., 2008; Bessarabova M et al., 2010***). Moreover, we demonstrated that DeepTXsolver accurately predicted crucial distribution properties, including mean, Fano factor, and bimodal coefficient, with a high correlation between theoretical and predicted summary statistics (***r > 0.99, p-value< 2.2*10***^***-16***^ ***of t-test Figure 2C***). It is noteworthy that, to assess the impact of the presence or absence of summary statistics on DeepTXsolver, we performed separate experiments using different loss functions. We demonstrated that incorporating summary statistics into the model led to quicker convergence of DeepTXsolver on the training set (***Figure 2D***) and yielded more robust and accurate predictions on the test set (***Figure 2E***).

Overall, DeepTXsolver effectively establishes mappings from parameters to stationary distributions. This approach circumvents the high computational resource requirements of classical simulation algorithms and ensures the algorithm’s efficiency and scalability in subsequent statistical inference, particularly in genome-wide inference.

### DeepTX infers genome-wide transcriptional kinetics from scRNA-seq data

Having obtained the trained neural network model, we can rapidly compute the stationary distributions of any TXmodel parameters. These distributions represent the outcomes of the underlying true gene expression process, referred to as the “true distribution” *P*_model_ (***Figure 3A***). However, during the inference process, the gene expression distributions measured by the scRNA-seq data we utilized, termed the “observed distribution of the data” *P*_data_, were subject to noise, including errors stemming from the measurement technique (***Sarkar A & Stephens M, 2021***) (***Figure 3A***). Thus, we introduced a mechanistic hierarchical model to bridge the gap between the true and observed distributions. This hierarchical model considers that the observed data results from the interplay between the gene expression process and the measurement process. It combines the two distributions using a convolutional form 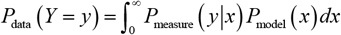 (see Methods 1.4). *P*_model_ ( *x*) represents the mixed negative binomial distribution approximated, whereas for *P*_measure_ ( *y* | *x*), we opted for the binomial distribution. This choice is informed by the fact that it can approximate hypergeometric distributions representing sequencing sampling without replacement when the sample size is sufficiently large. Consequently, the final distribution can maintain the form of a mixed negative binomial distribution (***Supplementary Note 1.4***), denoted as the observed distribution of the model *P*_data_ (***Figure 3A***).

**Figure 3.**
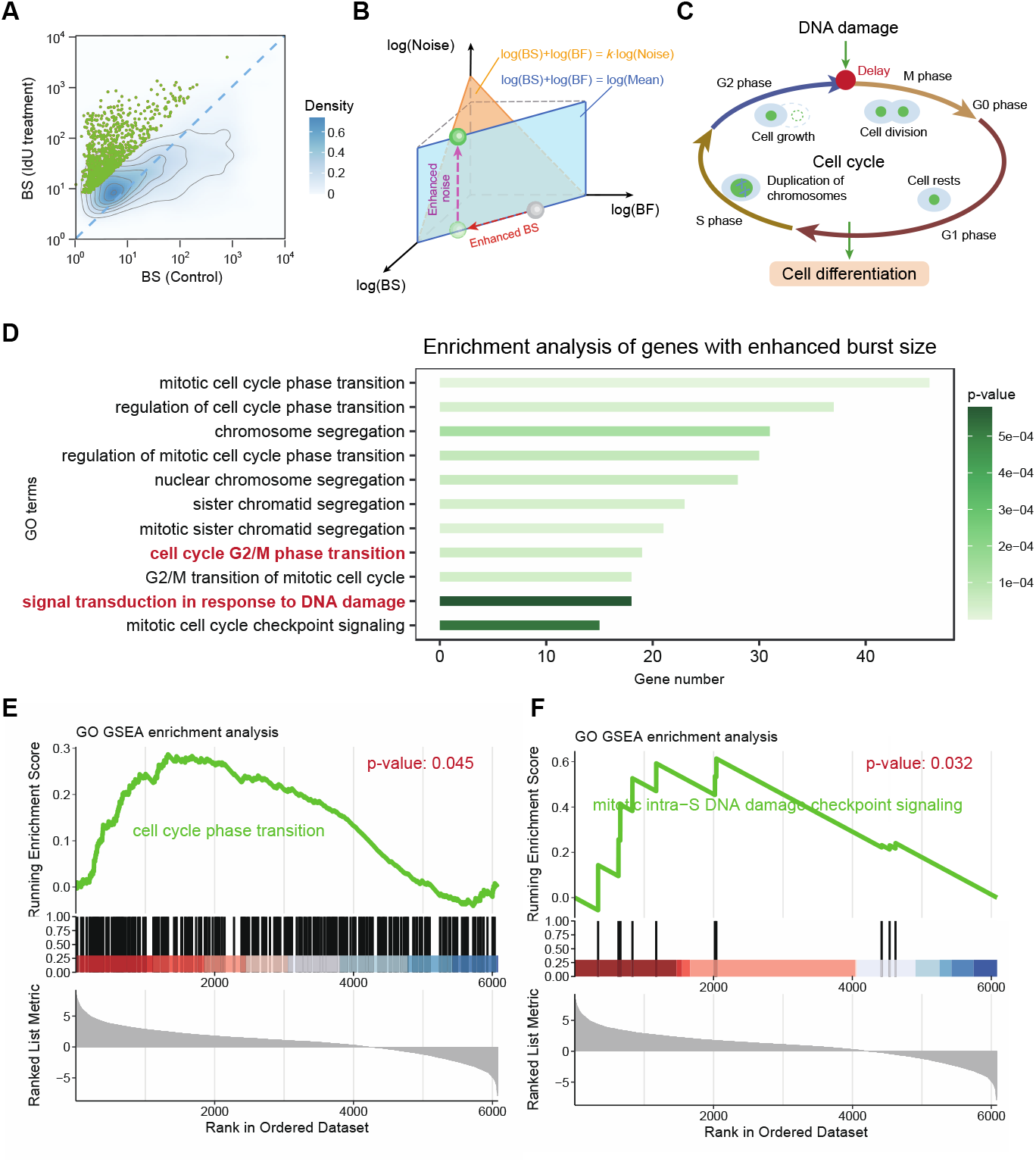
Inferring TXmodel from scRNA-seq data. **(A)** The dynamic parameters are sampled from a given prior distribution, and as input to the neural network, the solution of the corresponding dynamic model can be obtained by using this parameter. The solution of the dynamic model is mixed with the binomial distribution to obtain the model observed distribution. Loss values were obtained using the observed distribution of the data obtained from the scRNA-seq data compared to the observed distribution of the model. The loss values are optimized until convergence using gradient descent to obtain the parameters of the mechanism model and the posterior distribution of its parameters. **(B)** The scatters represent the real BS and BF values of the SSA synthetic set parameters, and the depth of the color represents the error between the inferred BS (BF) and the true BS (BF). **(C, D)** The blue solid line represents the edge density of the burst kinetics of the model, and the red dashed line represents the true value. **(E)** The marginal density for the five parameters of the model, where the red dashed line represents the true value. The relative error (RE) quantifies the discrepancy between the true parameter and the peak of the inferred posterior distribution

Leveraging the hierarchical model, we can construct a scalable and interpretable module called DeepTXinferrer, facilitating genome-wide inference of transcriptional burst kinetics from scRNA-seq data. First, we input the prior distribution of mechanistic model parameters and a set of scRNA-seq data, and we can get the observed distribution of the model. *P*_obsmodel_ . (derived successively from the trained neural network and the hierarchical model) and the observed distribution of the data *P*_obsdata_, respectively (***Figure 3A***). Second, we used the Hellinger distance *L* (***θ*** ) = *D* (*P*_obsmodel_, *P*_obsdata_ ) to measure the difference between *P*_obsmodel_ and *P*_obsdata_ (see Methods 1.4). It’s worth noting that the loss function remains solvable via gradients, despite the inclusion of complex computations, such as neural networks and hierarchical models. Specifically, to avoid the gradient-based optimization process from getting trapped in local optima, we employ a black-box optimization approach to select the initial point for parameter inference (***Das S & Suganthan PN, 2010***). Consequently, we utilize a gradient-based optimizer to update the iterative parameters until convergence. Lastly, we employ a method based on loss potentials, which computes the posterior distribution of the parameters during the optimization of the loss function (***Gaskin T et al., 2023***) (***Figure 3A, see Methods 1.4***). Subsequently, this process yields results of the transcriptional burst kinetics derived from the estimated parameters of the mechanistic model.

We assessed the effectiveness of DeepTXinferrer by conducting validation on synthetic data (see Methods 1.4). Synthetic samples were generated for each parameter set, and our inference algorithm was then applied to derive the output transcriptional burst kinetics (***see Methods 1.4***). As a result, the estimation of transcriptional burst kinetics demonstrated high accuracy and strong correlation across most parameter regions (***r > 0.99, p-value< 2.2*10^-16^ of t-test Supplementary Figure S3b, c***), particularly within the region of fully expressed mRNA abundance ***(Figure 3B)***. Moreover, the estimated posterior distribution of each parameter suggests that the peak value closely approximates the true value (***Figure 3C-E***). Additionally, we verified the robustness of our algorithm concerning the number of samples, observing that higher sample numbers corresponded to increased accuracy in estimation (***Supplementary Figure 3d, e***). Compared to previous work (***Larsson AJM et al., 2019; Luo S et al., 2023; Tang W et al., 2023; Gu J et al., 2025***), DeepTX demonstrates superior performance in both efficiency and accuracy (Supplementary Figure S4). In summary, our inference algorithm exhibited strong performance on synthetic data, affirming its reliability for application to real-world data in inferring transcriptional bursts.

In the subsequent sections, we will employ the DeepTXinferrer to explore how cells determine diverse fates in response to DNA damage stimuli through genome-wide regulation of transcriptional bursting. We analyzed three sets of scRNA-seq data representing distinct cell fates: cell differentiation, apoptosis, and survival. The first set of data is a mutually controlled scRNA-seq dataset of mESC cell differentiation induced by IdU drug treatment. This data set contains 12481 genes of transcriptomes from 812 cells and 13780 genes of transcriptomes from 744 cells, respectively. The second set of data is a mutually controlled scRNA-seq dataset of human colon cancer cells treated with low-dose 5FU drug that causes apoptosis. This data set contains 8534 genes of transcriptomes from 1673 cells and 7077 genes of transcriptomes from 632 cells, respectively. The third set of data is a mutually controlled scRNA-seq dataset of human colon cancer cells in which high-dose 5FU drug treatment resulted in cell-resistant survival. This data set contains 8534 genes of transcriptomes from 1673 cells and 6661 genes of transcriptomes from 619 cells, respectively.

### DeepTX reveals burst size enhancement induced by IdU treatment associated with cell differentiation

The administration of the IdU drug, a thymine analogue, influences the gene expression process within cells, subsequently impacting cell differentiation (***Desai RV et al., 2021; Li C et al., 2024***). The perturbation in the gene expression process primarily stems from DNA damage, as IdU is randomly integrated into the DNA chain, thereby inducing DNA damage (***Li C et al., 2024***). Although DNA damage caused by IdU treatment has little impact on genome-wide mean gene expression, it increases variance across the genome, thereby affecting cell differentiation (***Desai RV et al., 2021***). This is different from the understanding that increases in variance are usually caused by fluctuations in the mean gene expression (***Newman JR et al., 2006***). Meanwhile, the mean can be deconvolved as the product of BS and BF. This raises the question of the relationship between bursting dynamics and variance on a genome-wide scale, and how changes in bursting dynamics affect cell differentiation.

We used the DeepTX framework to infer a set of preprocessed scRNA-seq data (see Methods 2.1) to obtain underlying burst kinetics. The difference between the statistics and distribution obtained by inference and the statistics and distribution of sequencing data is minimal, ensuring the correctness of the inference results (***Supplementary Figure S5a-l***). Subsequently, we observed considerable variations in BS and BF across different genes (***Figure 4A, Supplementary Figure S6d-f***). Notably, genes with significantly increased BS exhibit corresponding decreases in BF, suggesting relative stability in the average gene expression level (***Supplementary Figure S6d-f***). Additionally, we observed a significant increase in transcriptional BS for most genes in the treatment group (***Figure 4A***). This indicates that the rise in variance primarily results from the enhanced BS induced by DNA damage (***Figure 4B***). Notably, this conclusion is consistent with the results of theoretical analysis (***Supplementary text 2***). To further explore the biological characteristics of BS enhancement, we identified genes with upregulated BS using differential analysis method. Further, we perform GO enrichment analysis (***Consortium GO, 2015***) on the identified genes. We observed that the genes with enhanced BS revealed significant enrichment in terms related to mitotic cell cycle checkpoint signaling, alongside pathways involved in DNA damage response and cell cycle transitions (***Smits VA & Medema RH, 2001***) (***Figure 4D***). Specifically, the mitotic checkpoint serves as a crucial safeguard to ensure accurate chromosome segregation and maintain genomic stability under DNA damage conditions. Activation of the mitotic checkpoint can influence cell fate decisions and differentiation potential. Sustained activation of the spindle assembly checkpoint (SAC) has been reported to induce mitotic slippage and polyploidization, which in turn may enhance the differentiation potential of embryonic stem cells. Subsequently, we performed Gene Set Enrichment Analysis (GSEA) (***Subramanian A et al., 2005***) on the identified differentially expressed genes associated with BS and also observed enrichment in pathways related to cell cycle transitions and mitotic intra-S DNA damage checkpoint signaling (***Figure 4E, F***). Similarly, we subjected genes significantly enhanced by BF in the treatment group (***Supplementary Figure S6f***) to GO analysis (***Supplementary Figure S6g***). While it is tempting to hypothesize that enhanced BS may contribute to DNA damage-related checkpoint activation and thereby influence cell cycle progression and differentiation, our current results only indicate an association between burst size enhancement and pathways involved in DNA damage response and checkpoint signaling. This is consistent with previous experimental observations showing that IdU treatment induces DNA damage and alters cell fate decisions through effects on the cell cycle (***Rosina M et al., 2019; Riccio F et al., 2022; Liu Z et al., 2017***) (***Figure 4C***). Additionally, research indicates that enhanced cell differentiation results from delayed cell cycle transitions (***Rosina M et al., 2019***).

**Figure 4.**
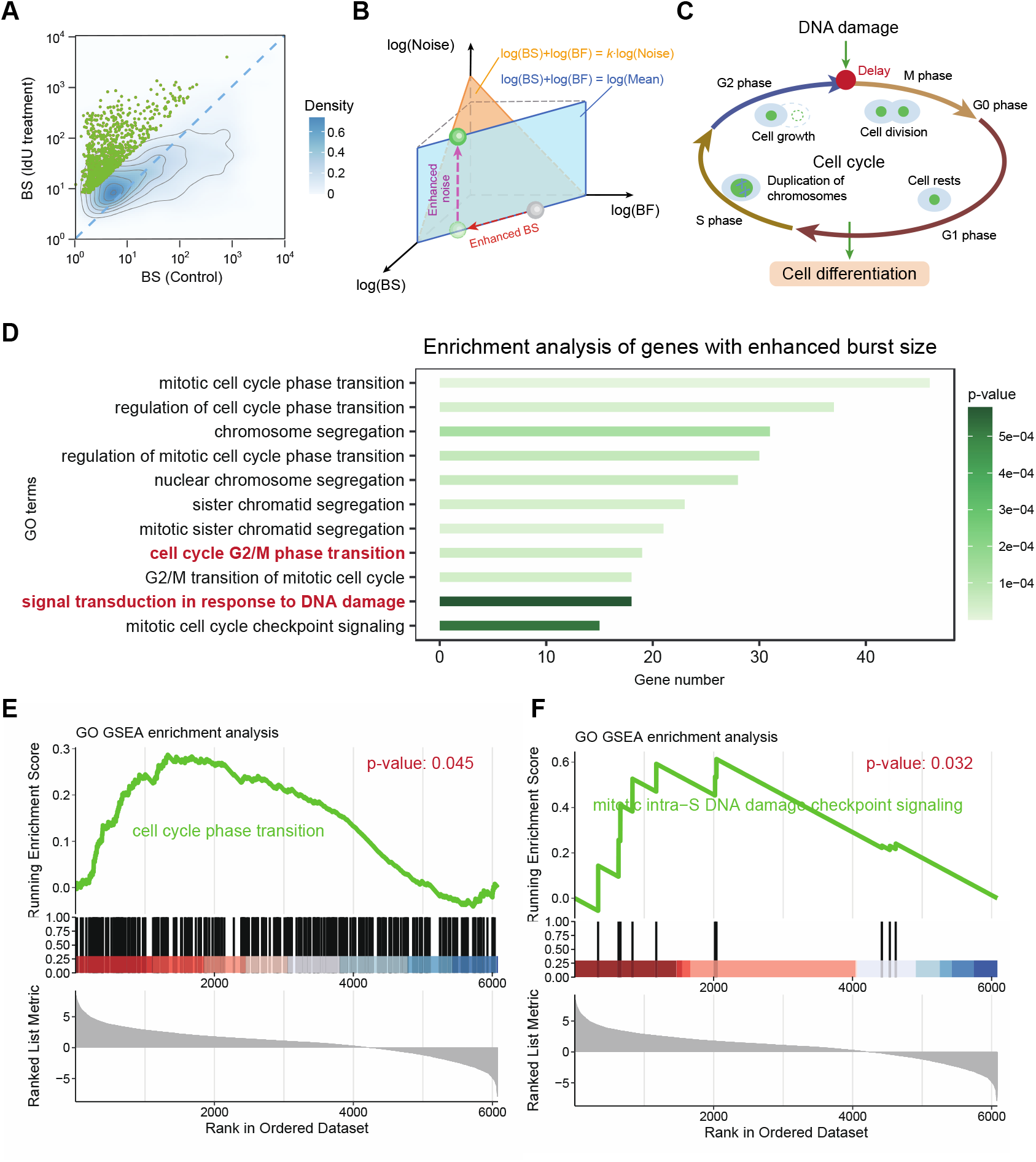
Burst size enhancer delays cell cycle. **(A)** Density map of BS of genes treated with IdU drugs compared to BS of genes not treated with drugs. Green dots represent up-regulated BS differential genes. **(B)** Schematic representation of the simultaneous enhancement of both BS and noise of a gene accompanied by the mean value of the gene remaining unchanged after IdU drug treatment. In the log-log-log 3D space, the mean expression level lies on a blue plane defined by log(BS) + log(BF) = log(Mean), indicating that it is determined by the product of burst size and burst frequency. The orange plane captures a scaling relationship between expression noise and burst kinetics, described by log(BS) + log(BF) = *κ* log(Noise), where *κ* is a constant reflecting their covariation. Notably, the trajectory of the green sphere shows that, under a fixed mean expression level (i.e., confined to the blue plane), increased gene expression noise primarily results from an increase in burst size. **(C)** The schematic diagram illustrates the hypothesis that DNA damage induced by IdU treatment may affect gene transcription, potentially influencing cell cycle transitions during mitosis and regulating cellular differentiation. **(D)** Gene GO enrichment analysis is performed on the green dots of BS in (A) to obtain enrichment pathway diagram. The bold red pathways highlight the key components involved in the cellular differentiation mechanism. **(E, F)** Pathways obtained by GSEA enrichment analysis of BS of genes.

### DeepTX reveals burst frequency enhancement under low-dose 5FU treatment and its association with apoptosis

The chemotherapeutic compound, 5-fluorouracil (5FU), exerts its cytotoxic effects on colorectal cancer cells by inflicting damage to their DNA (***Kuipers EJ et al., 2015; Bunz F et al., 1999; Chang GS et al., 2014***). This damage triggers apoptosis, a process of programmed cell death, thus inhibiting the growth and proliferation of malignant cells (***Kuipers EJ et al., 2015; Bunz F et al., 1999; Chang GS et al., 2014***). However, the impact of 5FU-induced DNA damage on the gene expression process and its potential relationship to cell apoptosis remains incompletely understood. In this section, we employ the DeepTX to investigate the effect of 5FU-induced DNA damage on the gene expression process in colorectal cancer cells with scRNA-seq data (***Park SR et al., 2020***). First, we inferred the burst kinetics from the scRNA-seq data of cells treated with low doses (10 μm) of 5FU and controls. After preprocessing the data (***see Methods 2.2***), we inferred the scRNA-seq data to obtain a distribution that fits the scRNA-seq data (***Supplementary Figure S7a-l***), as well as the underlying burst kinetics (***Figure 5A, B***).Further, we found that the mean difference between the two groups of scRNA-seq data was not significant (*p*-value = 0.5 of *t*-test), but the variance and BS were significantly different (***p-value<0.05 of t-test Figure 5A and Supplementary Figure S8a-c)***. Then, we performed a differential analysis of the BS and BF of gene expression between the control cells and the cells treated with 10 μM 5FU, and then performed GO gene enrichment analysis on differential genes (***Figure 5C and Supplementary Figure S8d-f***). We found that BS down-regulated differential genes were mainly enriched in apoptosis pathways (***Hongmei Z, 2012***), electron exchange pathways, and oxidation-related pathways (***Figure 5E and Supplementary Figure S8g***). Specifically, the enrichment analysis indicated that although the burst size of genes in 5FU-treated cells was generally larger than that in the control group, the genes primarily influencing cell apoptosis exhibited a down-regulated BS.

**Figure 5.**
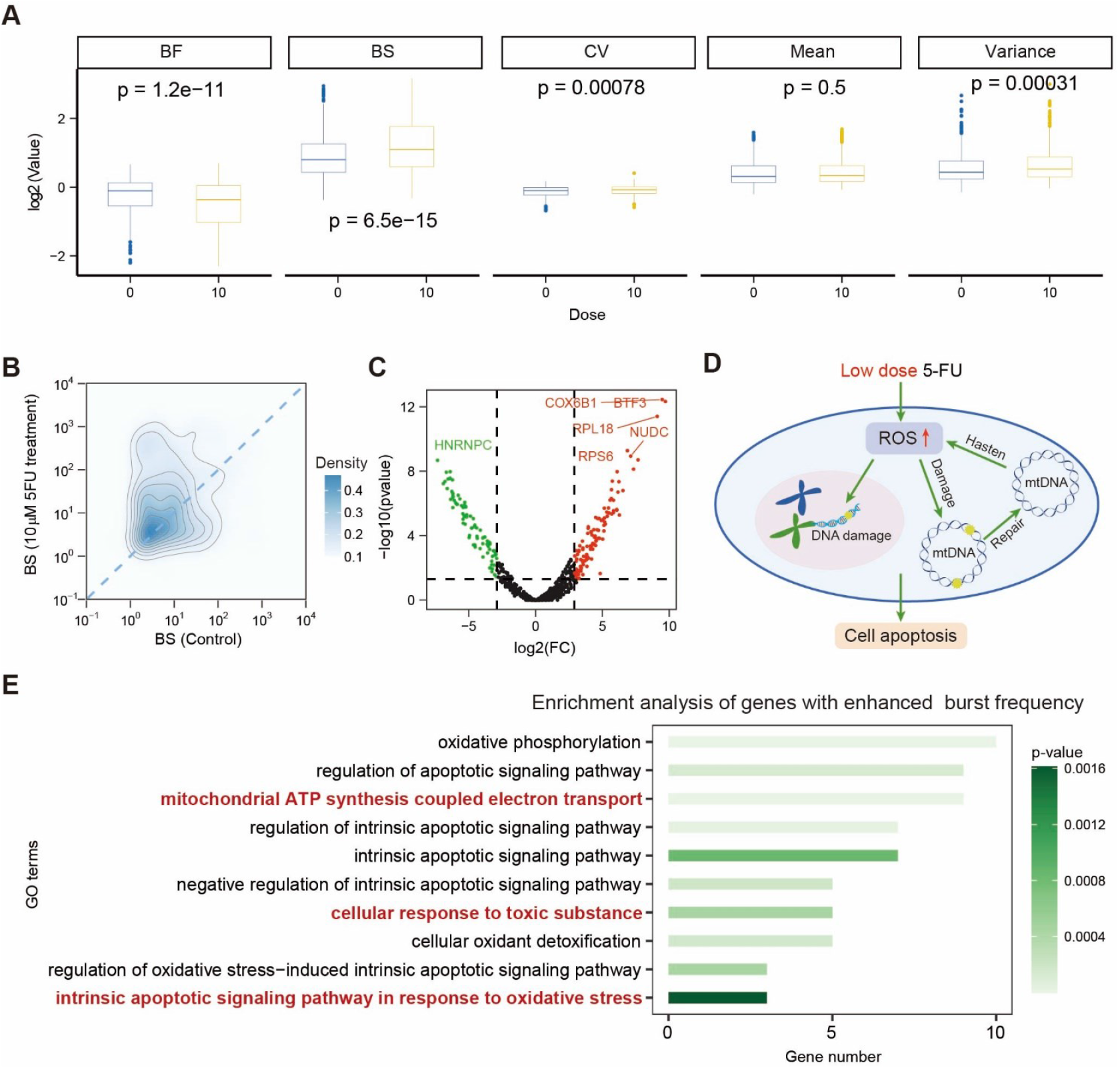
Low dose 5FU treatment induces cell apoptosis by enhancing burst frequency. **(A)** Comparison of gene bursting kinetics and statistics between control and 10-dose 5FU treated cells. **(B)** Density map of inferred burst size. **(C)** Volcano map of the fold changes (FC) of inferred burst size between control and 10-dose 5FU treated cells. **(D)** The diagram hypothesizing the potential association between low-dose 5FU treatment and the induction of apoptosis. **(E)** The enrichment analysis results were derived from genes that were downregulated in BS and upregulated in BF in the experimental group. The bold red pathways highlight the key components involved in the cellular apoptotic mechanism.

These enrichment results are in line with previously reported biological processes (***Figure 5D***). First, studies in recent years have found that many anticancer drugs trigger apoptosis and necrosis of cancer cells by activating the production of reactive oxygen species (ROS) (***Sun Y & Rigas B, 2008***). And experiments have shown that 5FU can induce ROS production in colorectal cancer cells (oxidation-related pathways) (***Laha D et al., 2015; Bwatanglang IB et al., 2016; Chenna S et al., 2022***). Meanwhile, mitochondria are the main source of ROS production, and defects in the mitochondrial electron transport chain will increase ROS production (electron exchange pathways) ***(Adam-Vizi V, 2005)***. As a result, excessive production of ROS can cause mitochondrial DNA damage and nuclear DNA damage ***(Figure 5 D***). In particular, when mitochondrial DNA is damaged, mitochondrial damage repair will further increase the pressure of ROS (***Figure 5D right***) (***Shokolenko IN et al., 2014***). Overall, our results support an association between 5FU-induced burst kinetics changes and pathways implicated in oxidative stress and apoptosis (***Figure 5D***) (***Handali S et al., 2018; Hwang PM et al., 2001***).

### DeepTX reveals burst frequency enhancement under high-dose 5FU treatment and its association with survival pathways

Low-dose 5FU treatment can induce cell apoptosis, while research shows high-dose 5FU treatment can induce cell resistance and evade apoptosis to survive (***Kuranaga N et al., 2001***). Hence, this part mainly studies the impact of differences in burst kinetics of gene expression between high-dose (50 μm) 5FU-treated and control colorectal cancer cells on cell fate decisions.

After data preprocessing (see Methods 2.2), we similarly observed the phenomenon that the average expression value of genes remained stable, while there is an enhanced increase in the variance genome-wide (***Supplementary Figure S10a-c***). Then, after preprocessing the data (see Methods 2.2), we inferred the scRNA-seq data to obtain the distribution that fits the scRNA-seq data (***Supplementary Figure S9a-l***), as well as the underlying burst kinetics (***Figure 6A***). Further, we performed a difference analysis on the BS of the two sets of inferred data, and selected genes with large differences in BS and BF for GO gene enrichment (***Figure 6A, B and Supplementary Figure S10d-f***). We found that the BS down-regulates differential genes were mainly enriched in the Cajal body related pathway, the electron exchange pathway, and telomerase-related pathways (***Nabetani A & Ishikawa F, 2011***), but the apoptosis-related pathways were not significant (***Figure 6D***). In particular, according to experimental studies, telomerase-related pathways can activate telomerase and play an important role in the elongation of telomeres (***Cristofari G et al., 2007***). These enrichment results suggest that high-dose 5FU treatment may be associated with processes such as telomerase activation and mitochondrial function maintenance, both of which have been implicated in cell survival and apoptosis evasion in previous experimental studies. For example, prior work indicates that hTERT translocation can activate telomerase pathways to support telomere maintenance and reduce oxidative stress, which is thought to contribute to apoptosis resistance (***Kuranaga N et al., 2001***). Overall, the observed transcriptional bursting changes are consistent with these reported survival-associated mechanisms.

**Figure 6.**
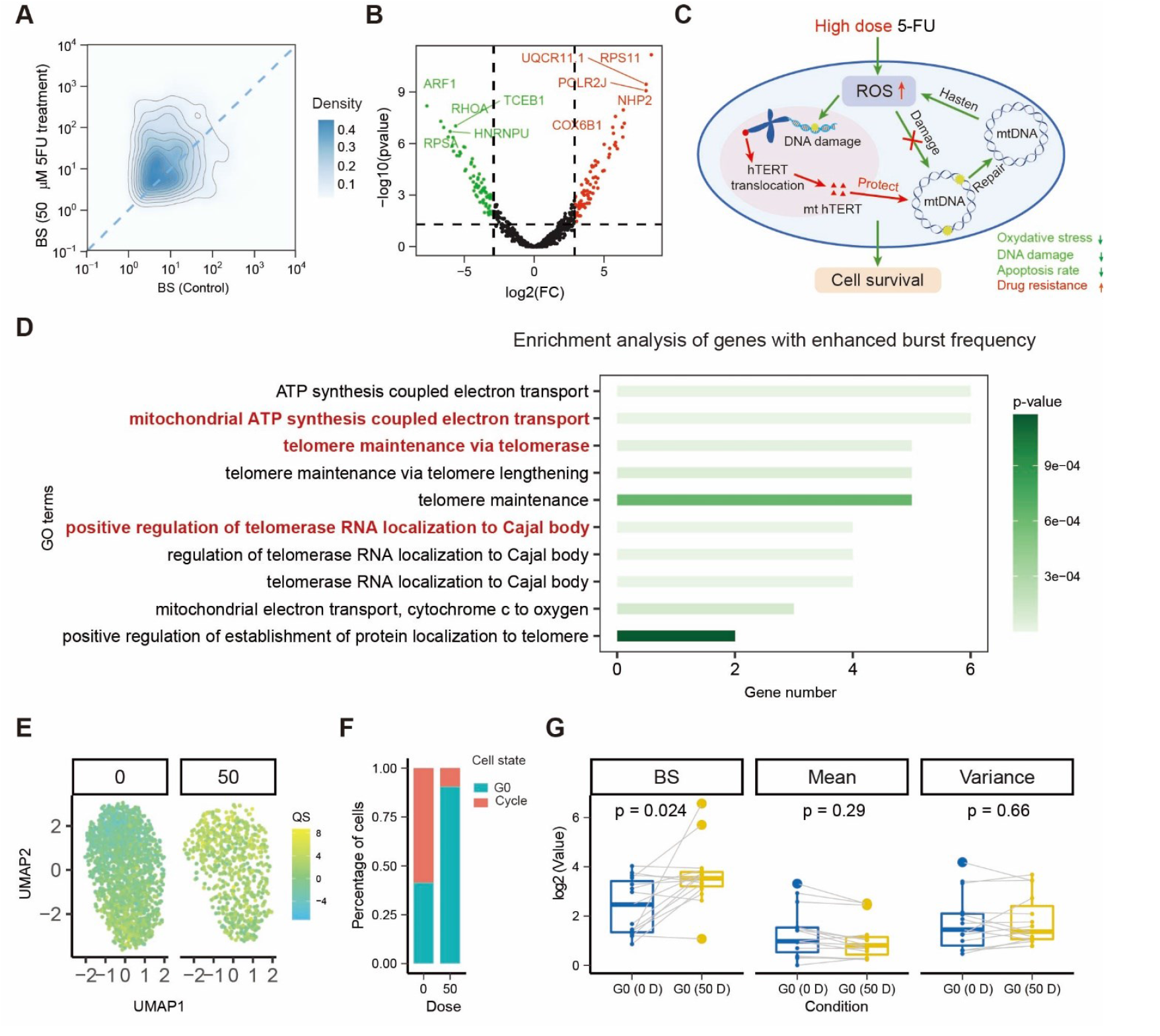
High dose 5FU treatment may promotes cell drug resistance by altering burst frequency. **(A)** Density map of inferred burst size. **(B)** Volcano map of the fold changes (FC) of inferred burst size between control and 50-dose 5FU treated cells. **(C)** Schematic diagram hypothesizing the mechanism by which 5FU drug treatment may induce cells to acquire antioxidant properties, potentially enabling continued survival. **(D)** The enrichment analysis results were derived from genes that were downregulated in BS and upregulated in BF in the experimental group. **(E)** UMAP of G0 arrest quality score (QS) of control and 50-dose 5FU treated cells. **(F)** Proportion of G0 arrested (drug-resistant) and cycling (non-resistant) cells. **(G)** Comparison of gene bursting kinetics and statistics of G0 arrested genes between control and 50-dose 5FU treated cells. The light grey lines indicate the trend of changes in the corresponding bursting kinetics values, mean and variance for each gene.

Gene enrichment results are consistent with existing ways of cells evading apoptosis (***Figure 6C***). That is, endogenous ROS production exceeds the cellular antioxidant defense capacity, leading to chemical damage to mitochondrial DNA (mtDNA). Meanwhile, high doses of 5FU causes Human Telomerase Reverse Transcriptase (hTERT) translocation, which on the one hand triggers telomerase activity and maintenance pathways, which enables cells to evade apoptosis (***Kuranaga N et al., 2001***). On the other hand, this translocation will enrich hTERT in mitochondrial DNA, thereby reducing oxidative stress and mitochondrial DNA damage to protect mitochondria and prevent cells from apoptosis (***Lipinska N et al., 2017***).

Resistance to therapy has also been associated with G0 arrest, a non-proliferative state characteristic of persister cells (***Wiecek AJ et al., 2023***). To verify the presence of resistant cells in cells treated with high doses of 5FU, we selected genes related to G0 arrest and performed GSVA (***Hänzelmann S et al., 2013***) to obtain the G0 arrest quality score (***Wiecek AJ et al., 2023***) of each cell and perform UMAP dimensionality reduction visualization (***Figure 6E***). Through arrest quality score discrimination, we found that the proportion of resistant cells was large among cells treated with high doses of 5FU (***Figure 6F***). This result was consistent with the results of our enrichment analysis. Specifically, by comparing the BS and mean values of these genes, we found that most of the BS of G0 arrest related genes were up-regulated, but the mean changes were small (***Figure 6G***).

## Discussion

Gene expression is inherently stochastic, and transcriptional bursting contributes to inter-individual variability in gene expression patterns (***Dar RD et al., 2012; Zenklusen D et al., 2008; Phillips R et al., 2012***), although it represents only one of multiple sources of cellular heterogeneity. Experimental observations confirm this bursting mechanism as a cellular response to environmental changes, including DNA damage (***Friedrich D et al., 2019; Desai RV et al., 2021***), but its precise role in DNA damage response and subsequent cell fate decisions remains an emerging hypothesis. In this study, we introduced a mechanism-based deep learning framework (DeepTX) for performing genome-wide inference on transcriptional burst kinetics of a comprehensive gene expression model using scRNA-seq data. DeepTX provides an efficient computational approach to characterize burst kinetics across genes, enabling the generation of testable hypotheses about how transcriptional bursting may be associated with DNA damage response and potential changes in cellular state.

Utilizing DeepTX, we analyzed three sets of scRNA-seq data comprising control and stimulus pairs driving different cell fate decisions: cell differentiation, apoptosis, and survival. This analysis unveiled significant insights into the underlying mechanisms of transcriptional burst kinetics. Firstly, the IdU drug treatment–induced enhancement of BS in genes appears to be associated with a delayed transition in the cell mitosis phase, which may in turn be related to changes in cell reprogramming and differentiation. Interestingly, while prior studies have suggested that noise enhancement contributes to cell fate decisions, our observations indicate that BS enhancement may reflect a deeper regulatory feature correlated with cell differentiation. Secondly, low-dose 5FU treatment is associated with increased oxidative stress and an accompanying elevation in BF, which may relate to enhanced apoptotic activity. Consistent with previous findings linking apoptosis to ROS stress, our study suggests that ROS-induced DNA damage is correlated with changes in gene expression burst kinetics, which may contribute to the apoptotic outcomes observed. Thirdly, high-dose 5FU treatment induces telomerase extension, mitigating oxidative stress and fostering drug resistance by upregulating BF. Despite prior evidence of drug-resistant cells, our enrichment analysis underscores telomerase extension’s role in resisting oxidative stress, underscoring the impact of BF enhancement.

We highlight several advantages of our DeepTX framework, which differs significantly from previous work that utilizes scRNA-seq data to study diverse noise sources from several aspects (***Eling N et al., 2019; Faure AJ et al., 2017; Morgan MD & Marioni JC, 2018; Ochiai H et al., 2020)***. Firstly, DeepTX employs a mechanistic hierarchical model that is both interpretable and extensible. To enhance the model’s realism in interpreting scRNA-seq data, we developed a novel approach integrating the sequencing process and the underlying gene expression process. Specifically, we employed the TXmodel, a complex non-Markovian stochastic dynamic model that extends the waiting times for gene OFF and ON states into a non-exponential distribution, reflecting the multi-step nature of gene regulation. As a result, TXmodel not only captures gene expression dynamics under DNA damage but also depicts other multi-step regulatory processes, such as chromatin opening ***(Lomvardas S & Thanos D, 2002)***, preinitiation complex formation ***(Fuda NJ et al., 2009)***, transcription factors binding ***(Nordick B et al., 2022)***. Secondly, despite the mathematical complexity of the mechanistic hierarchical model, DeepTX renders it tractable. This is achieved by (i) directly mapping model parameters to corresponding stationary distributions via a neural network trained on extensive simulated single-cell RNA data and theoretical analysis summary statistics for TXmodel; (ii) approximating the stationary distribution output of TXmodel, coupled with the sequencing process, by a mixed negative binomial distribution, maintaining the final distribution format. Thirdly, DeepTX’s inference process is scalable, allowing genome-wide bursting kinetics inference from static scRNA-seq data. Despite the computational complexity of parameter inference involving hierarchical models and trained neural networks, DeepTX retains auto-differentiation for efficient and accurate gradient computation.

The DeepTX research paradigm contributes to a growing line of area aiming to link mechanistic models of gene regulation with scRNA-seq data. Maizels provided a comprehensive review of computational strategies for incorporating dynamic mechanisms into single-cell transcriptomics ***(Maizels RJ, 2024)***. In this context, RNA velocity is one of the most important examples as it infers short-term transcriptional trends based on splicing kinetics and deterministic ODEs model. However, such approaches are limited by their deterministic assumptions and cannot fully capture the stochastic nature of gene regulation. DeepTX can be viewed as an extension of this framework to stochastic modelling, explicitly addressing transcriptional bursting kinetics under DNA damage. Similarly, DeepCycle, developed by Sukys and Grima ***(Sukys A & Grima R, 2025***), investigates transcriptional burst kinetics during the cell cycle, employing a stochastic age-dependent model and a neural network to infer burst parameters while correcting for measurement noise. By contrast, MIGNON integrates genomic variation data and static transcriptomic measurements into a mechanistic pathway model (HiPathia) to infer pathway-level activity changes, rather than gene-level stochastic transcriptional dynamics ***(Garrido-Rodriguez M et al., 2021***). In this sense, DeepTX and MIGNON are complementary, with DeepTX resolving burst kinetics at the single-gene level and MIGNON emphasizing pathway responses to genomic perturbations, which could inspire future extensions of DeepTX that incorporate sequence-level information.

The DeepTX research paradigm is poised for extension to encompass both dynamic models and biological data. On the one hand, our focus is currently confined to the transcriptional processes influenced by DNA damage, limiting the explanatory scope of the TXmodel to larger-scale biological phenomena such as alternative splicing (***Wan Y et al., 2021; Gorin G & Pachter L, 2022***), protein translation (***Luo S et al., 2023; Golan-Lavi R et al., 2017***), epigenetic modification (***Nicolas D et al., 2018***), and chromatin movement (***Liu T et al., 2016; Wang Z et al., 2024; Wang Z et al., 2023***). Moreover, further development is needed to incorporate additional regulatory factors, including signaling pathways (***Garrido-Rodriguez M et al., 2021***), cell cycle progression, cell size (***Sukys A & Grima R, 2025***), and leaky transcription (***Kepler TB & Elston TC, 2001***). These broader biological processes necessitate more intricate dynamic models, often challenging to solve and thus challenging to apply to realistic biological data.

On the other hand, inference from static scRNA-seq relies on the steady-state assumption, which has some limitations. First, in some scenarios, the cell system may exhibit highly transient transcriptional programs that do not satisfy stationarity, leading to biased or misleading parameter estimates. For example, immediately following a synchronized developmental stimulus such as serum shock induced activation of immediate-early genes, transcription levels increase significantly (***Hope B et al., 1992***). Second, because DeepTX infers the mean burst frequency and size across the population, it cannot recover the underlying time-resolved dynamics or distinguish heterogeneous kinetic subpopulations. Third, lack of time-resolved measurements may affect the accuracy of inferences about dynamic parameters, especially the unidentifiability of parameters inferred from steady-state distributions, i.e., multiple parameters leading to the same steady-state distribution. The unidentifiability of individual parameters is a common and critical problem in systems biology studies. Hence, in DeepTX, we employed a Bayesian approach based on loss potential to infer the posterior distributions of the parameters. Although DeepTX also encounters the issue of unidentifiability for individual parameters (Supplementary Figure S11), the multimodal nature of the posterior distribution suggests that multiple distinct parameter sets can produce similarly good fits to the observed data, highlighting the inherent non-identifiability of the model. Nevertheless, in the multimodal posterior distribution, at least one of the posterior peaks aligns closely with the ground truth, thereby demonstrating the validity of the inferred result. In future work, integrating time-series single-cell measurements with other emerging data types could help overcome these limitations. Examples include increasingly comprehensive transcriptome data with both temporal and spatial resolution (***Wen L & Tang F, 2022***), as well as state-of-the-art multi-omics data (***Maizels RJ, 2024;Miao Z et al., 2021***). Moreover, another limitation of the current implementation is that DeepTX is only trained on simulated datasets. While providing controlled benchmarks for mechanistic validation, it may not fully recapitulate the heterogeneity and stochastic complexity of real biological systems. In the future work, publicly available annotated scRNA-seq datasets could be used to complement this simulation-based training strategy and enhance generalizability.

In conclusion, DeepTX provides a step toward inferring complex transcriptional kinetics from scRNA-seq data through mechanism-based deep learning inference. It sheds light on the augmented burst kinetics in DNA damage response across various cell fate decisions, thereby offering novel insights.

## Methods

### 1 The DeepTX framework

#### 1.1 Framework overview

The DeepTX Framework aims to infer the underlying bursting dynamics from static single-cell RNA sequencing (scRNA-seq) data to explore the impact of changes in bursting dynamics on cell fate decisions. The DeepTX framework includes two modules, DeepTXsolver and DeepTXinferrer. DeepTXsolver can efficiently and accurately solve transcription models using neural network architecture, and DeepTXinferrer can infer scRNA-seq data to obtain its underlying transcription burst kinetics using Bayesian methods.

The input of the DeepTXsolver is the parameters of the mechanism model, and the output is the corresponding stationary distribution solution of the model. The input to the DeepTXinferrer is the discrete probability distribution of gene expression obtained from scRNA-seq data, and the output of the model is the parameters of the mechanism model.

The construction of the DeepTX framework consists of the following parts: (1) Model the gene expression process. (2) Train a neural network that solves the mechanism model using the data set generated by stochastic simulation algorithm (SSA) simulation. (3) Hierarchical model modeling to infer underlying burst kinetic parameters from scRNA-seq data.

When building the DeepTX Framework, we made the following assumptions: (1) The gene expression of cells was in a stationary distribution during sequencing. This assumption has been widely adopted in scRNA-seq studies, as it effectively captures mRNA expression distributions and enables the inference of underlying dynamic parameters. (***Larsson AJM et al., 2019; Luo S et al., 2023; Ramsköld D et al., 2024; Gupta A et al., 2022***) (2) Gene expression counts of the same cell type follow the same distribution. Given that cells of the same type typically share similar transcriptional programs, their gene expression distributions are approximately identical, supporting the validity of this commonly used assumption. (3) The solution of the model can be approximated by a mixed negative binomial distribution. Since the stationary solution of chemical master equation can be expressed as a Poisson mixture, which can be more efficiently approximated using negative binomial mixtures (***Gorin G et al., 2024***). (4) The state space sampled from a sufficiently long single simulation is statistically equivalent to that obtained from multiple simulations at steady state in gene expression models (***Kuntz J et al., 2021***).

#### 1.2 Modelling stochastic gene expression processes

To recover dynamic transcriptional burst kinetic parameters from static scRNA-seq data, we require the stochastic modeling approach to profile the gene expression process. The traditional telegraph model is often used to model gene transcriptional bursting, which assumes that switching between genes follows an exponential distribution (i.e., switching is a single-step process) (***Peccoud J & Ycart B, 1995; Kim JK & Marioni JC, 2013***). However, this assumption is an oversimplification of the gene expression process in the context of DNA damage. Due to the fact that DNA damage slows down and even stops the elongation of RNA pol II across the double-strand DNA (***Lans H et al., 2019***), the gene expression process should be regarded as a multi-step process. This property causes the waiting time for the gene switching between active and inactive states to follow a non-exponential distribution (***Daigle BJ, Jr. et al., 2015; Zhang J & Zhou T, 2019; Zhang JJ & Zhou TS, 2019***). Therefore, we use TXmodel (***Luo S et al., 2023***) to better characterize the gene transcription process in the context of DNA damage ***(Figure 2***), whose chemical reaction network (CRN) can be written as follows:

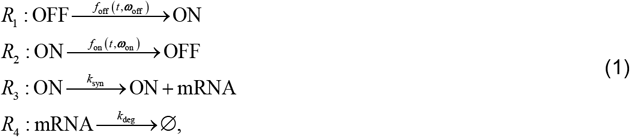

where *R*_1_ and *R*_2_ describe the gene-state switching randomly between the ON state and OFF state, and *f*_off_ (*t*,***ω***_off_ ) and *f*_on_ (*t*,***ω***_on_ ) are the distribution of dwell time in ON and OFF state, respectively. Specifically, dwell time distribution refers to the probability distribution of the time in which a gene remains in a particular transcriptional state (ON or OFF) before transitioning to the other state. While the gene is in the active state, the synthesis of mRNA molecules is governed by a Poisson process with a constant synthesis rate *k*_syn_ ( *R*_3_ ). Meanwhile, each mRNA molecule can be stochastically degraded ( *R*_4_ ) with a constant degradation rate *k*_deg_ . Overall, the TXmodel can be determined by a set of parameters, denoted as ***θ*** = (***ω***_off_,***ω***_on_, *k*_syn_, *k*_deg_ ) .

The TXmodel as an extension of the traditional telegraph model can capture more realistic transcriptional burst kinetics, with BF and BS being two important parameters. The definition of BF is

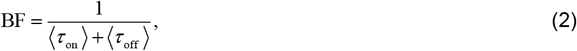

where 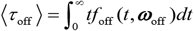 and 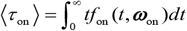 are the mean dwell time in OFF and ON state, respectively. BF represents the average time it takes for the promoter to complete one full stochastic cycle between its active and inactive states. This definition is slightly different from the traditional definition 1/ (*k*_deg_ *τ_off_ ), which can be regarded as a simplified version of our definition (Eq. 2), under the assumptions that τ _on_ is negligible and *k*_deg_ = 1 (i.e., rate parameters are normalized to be dimensionless). Although it is reasonable to neglect activation time τ _on_, as it is typically much shorter than inactive time under some conditions, we chose a more complete way to define the burst frequency so that it is applicable to more general situations. Additionally, Eq. 2 has been widely used in recent literature (***Ramsköld D et al., 2024; Zoller B et al., 2018; Hoppe C et al, 2020)***. The definition of BS is

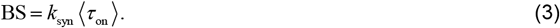

BS refers to the average number of mRNA transcripts produced during a single transcriptional activation event of a gene. Notably, the mean transcription level of mRNA ⟨*y*⟩ can be expressed as 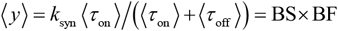.

Without loss of generality, the gamma distribution can be used to characterize non-exponential distributions, i.e., 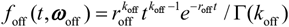 such that ***ω***_off_ =(*r* _off_, *k*_off_) and 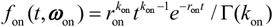 such that ***ω***_on_ =(*r*_on_, *k*_on_ ), where Γ(·) is Gamma function, *r*_off_ and *r*_on_ are the rate-parameters of state switching, *k*_off_ and *k*_on_ are the possible number of reaction steps between OFF and ON state. The rationale for employing the gamma distribution lies in the complex, multi-step nature of promoter state transitions. Biologically, both the “ON” and “OFF” states of a promoter encompass multiple underlying sub-states, with transitions governed by a series of molecular events, each characterized by exponentially distributed waiting times. When aggregated, these sequential exponential processes yield non-exponential overall waiting time distributions. The gamma distribution, being a natural analytical extension of such a convolution of multiple exponential distributions, provides a mathematically and biologically appropriate approximation for such processes. Here, in the TX model, the state-switching waiting time follows a non-Markovian distribution (i.e., it is not exponentially distributed), indicating that the system exhibits non-Markovian dynamics.

It is a challenge to solve the stationary distribution of this gene expression system directly from a theoretical approach. But we proved that the binomial moments of this system can be obtained in an iterative form, by rationalizing this gene expression system as a queueing theory model. Thus, we can theoretically compute some important summary statistics ***s*** (***θ*** ) given a parameter ***θ***, such as mean, variance, kurtosis, and bimodal coefficient (***see details in Supplementary Text 1.2***).

#### 1.3 Training neural network to solve gene expression model

The stationary distribution of CRN (1) is difficult to be solved analytically. In general, stochastic simulation algorithms are used to approximate the solution of chemical master equations (***Boguñá M et al., 2014; Masuda N & Rocha LE, 2018***). However, the inference process involves updating the parameters, and each update requires a simulation to obtain the stationary distribution, which makes statistical inference time-consuming. We note that deep learning methods may have potential applications in solving this challenge because it is widely used for approximating the solution of partial differential equations, chemical master equations, and non-Markovian chemical master equations (***Jiang Q et al., 2021; Wang S et al., 2019; Davis CN et al., 2020; Gupta A et al., 2021; Bar-Sinai Y et al., 2019)***.

Therefore, in this part, we used a neural network to approximately solve CRN (1). This enables CRN (1) to be solved efficiently and accurately. Details will be introduced in the following part.

##### Generating synthetic training dataset

To construct the mapping from the parameters of TXmodel to the stationary distribution, we first need to generate the neural network’s training set, i.e., a large-scale range of parameter sets as features and corresponding probability distributions as labels. For the features, we used the Sobol algorithm (***Sobol IM, 1967***) to conduct large-scale sampling to generate the neural network data set, which can cover the entire parameter space more evenly than random sampling.

For the labels, we perform the SSA of the TXmodel to obtain the stationary distribution given a set of preset parameters. For clarity, the synthetic training dataset is denoted by {(***θ***_*i*_, *P*_simulation,*i*_ )}, where ***θ***_*i*_ presents the model parameters of TXmodel, and *P*_simulation,*I*_ represents the stationary distribution solution obtained by SSA (***Figure 2A***). Specifically, the value range of each parameter component of ***θ***_*i*_ is *k*_off_ ∈(1,15), *r*_off_ ∈(0.1,10), *k*_on_ ∈(1,15), *r*_on_ ∈(0.01,10), *k*_syn_ ∈(0.1, 400) . The experimentally measured values of the mean duration of the OFF state, burst duration, transcriptional burst sizes, and the number of intermediate steps in transcriptional state transitions fall within the specified parameter ranges. (***Tunnacliffe E & Chubb JR, 2020; Gupta A et al., 2022***).

##### Deep learning model architecture

We build the basic deep learning model architecture in DeepTX via fully connected neural networks, which has been effectively used to solve approximate solutions from biochemical reaction networks (***Sukys A et al., 2022; Gupta A et al., 2021***). The deep learning model architecture consists of an input layer, hidden layers, and an output layer.

Firstly, the input layer contains 5 neurons corresponding to the parameters of the mechanism model ***θ***_*i*_ = (*k*_off,*i*_, *r*_*o*ff,*i*_, *k*_on,*i*_, *r*_on,*i*_, *k*_syn,*i*_ ) . Secondly, the input layer is followed by *n*_hidden_ hidden layers with each layer containing 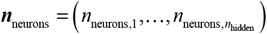 neurons. For the neurons in each hidden layer, we implement a nonlinear mapping by the commonly used ReLU activation function (***Supplementary Figure S1***):

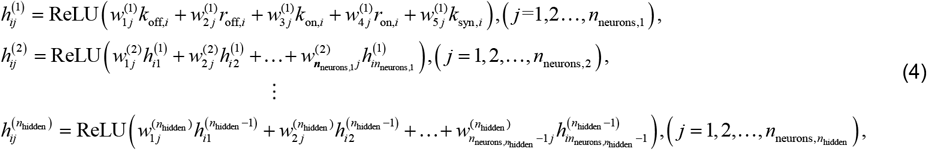

where 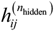 represents the value of the *j* -th neuron of the *n*_hidden_ -th hidden layer of the *i* -th group of parameters, 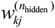 represents the neural network weight connecting the *k* -th neuron of the *n*_hidden_ -th hidden layer and the *j* -th neuron of the (*n*_hidden_ −1) -th hidden layer.

Thirdly, In the output layer, we aim to allow neurons to approximate synthetic scRNA-seq data generated by TXmodel. Negative binomial distributions are a class of distributions that are widely used in modeling scRNA-seq data and have been shown to account for both the “drop-out” phenomenon and the fact that the Fano factor is greater than 1 in scRNA-seq data (***Risso D et al., 2018; Love MI et al., 2014***). Also, the negative binomial distribution has been employed as an approximate stationary solution to the TXmodel for many gene expression systems (***Sukys A et al., 2022; Öcal K et al., 2022***). We assign weights and parameters (*a*_*ij*_, *m*_*ij*_,*l*_*ij*_ ) to the output layer neurons (***Supplementary Figure S1***):

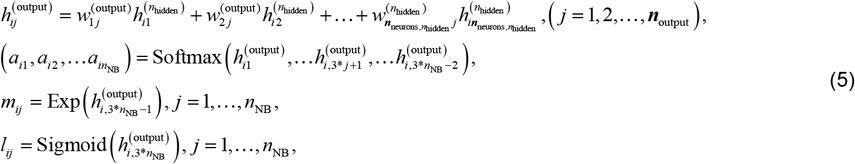

where 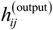 represents the value of the *j* -th neuron of the output layer of the *i* -th set of parameters, *a*_*ij*_ is the weight coefficient of the negative binomial distribution, satisfying 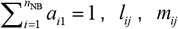 represents the *i* -th parameter of the *j* -th negative binomial distribution of the mixed negative binomial distribution, *n*_NB_ is the number of negative binomial distribution. And the mixed negative binomial distributions can be expressed as (***Supplementary Figure S1***):

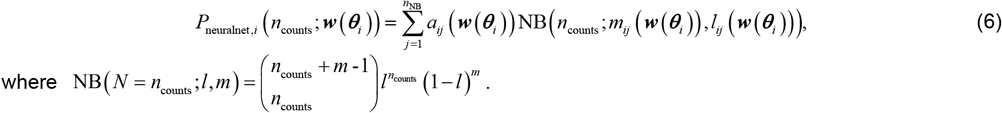

where 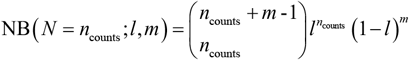.

##### Loss function of DeepTXsolver

Our loss function includes two parts: one part is the KL divergence between the mixed negative binomial distribution generated by neural network *P*_neuralnet,*i*_ and label distribution *P*_simulation,*i*_ generated by SSA, and the another part is the log difference of the moment statistics between theoretical solution of the TXmodel *s*_*ij*_ and the output of the neural network *ŝ*_*ij*_ :

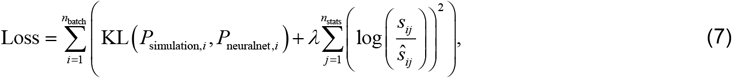

where 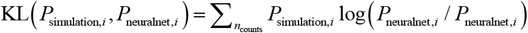, and *λ* is the hyperparameter, represents the weight coefficient of the statistical loss, and *λ* = 0.1, *n*_batch_ represents the number of samples, where each sample is characterized by (***θ***_*i*_, *P*_simulation,*i*_ ) in each batch, and *n*_stats_ is the number of summary statistics. For the latter part of loss function, we can still obtain the moment statistic by theoretical derivation, although the analytic distribution is difficult to solve. Adding moment statistics to the loss function can make the neural network more robust (***Figure 2E***). Specifically, we use four statistics here: *s*_*i*1_ indicates the mean, *s*_*i*2_ denotes the Fano factor, *s*_*i*3_ denotes kurtosis, *s*_*i*4_ represents the bimodal coefficient. Specifically, the Fano factor, which normalizes variance by the mean, provides a robust measure of transcriptional noise across genes or conditions with varying expression levels. It’s worth mentioning that the statistics here is scalable and we can add more information inside the loss function.

##### Training details of DeepTXsolver

Effective model training necessitates the selection of an appropriate optimizer to minimize the loss function. In our case, we employed the Adam optimizer (***Kingma DP & Ba J, 2014***), a widely adopted choice in deep learning due to its robust performance. And each time, a small batch of samples is selected to update the neural network parameters ***w*** (***Goodfellow I et al., 2016***), and the iteration of all samples is called an epoch. Furthermore, we utilized the Glorot Uniform method to establish an optimal initial point for our neural network parameter (***Glorot X & Bengio Y, 2010***).

Batch size and learning rate are key optimization hyperparameters affecting model convergence. A smaller batch size enhances model generalizability but lengthens training time, hence a batch size of 64 is chosen for balance (***Keskar NS et al., 2016***). In gradient descent process, the learning rate significantly influences the step size, where large rates may induce instability or divergence, and conversely, small rates may precipitate premature convergence to local optima. To counteract this, a decaying learning rate is used, allowing larger early updates for faster convergence and preventing instability in later stages. Further, training of the model stops if it has been trained for over 200 epochs or if the learning rate has been reduced five times. These criteria are adaptive and efficient as further training does not significantly enhance the model’s performance.

##### Hyperparameter tunning of DeepTXsolver

To obtain a neuron network model with accurate prediction and generalization, we compared the model architectures in terms of the number of neurons per layer, the number of hidden layers, and the number of neurons in the output layer, and compared the performance of the model with different dataset sizes. And we applied the Hellinger distance between the true distribution and the predicted distribution by the neuron network as a judgment criterion.

First, we compared the performance of architectures with a single hidden layer with (16, 32, 64, 128, 256, 512) neurons, and found that the performance of the model barely increased after increasing to 128 neurons (***Supplementary Figure S2a***). Secondly, after comparing the performance of architectures with different numbers of hidden layers, we found that a single hidden layer has the best performance (***Supplementary Figure S2b***). Thirdly, comparing the performance of 3*i* (*i* = 1, 2,… 10) neurons in the output layer, we found that after 12 neurons (4 negative binomial distribution mixtures), the performance of the model was not improved (***Supplementary Figure S2c***). Particularly, four negative binomial distribution mixtures are able to fit the bimodal nature of the distribution as well as ensure good training efficiency (***Öcal K et al., 2022***).

Finally, to obtain a dataset that contributes to convergence as well as model generalization while ensuring training efficiency, we compared the model performance for dataset sizes of (500, 1000, 5000,10000,15000) and found that increasing the dataset size further does not significantly improve the model performance after to 15000 samples (***Supplementary Figure S2d***). Therefore, we identified a single hidden layer of 128 neurons and an output layer of 12 neurons as the appropriate model architecture, and 15,000 samples as the appropriate training set.

#### 1.4 Inferring burst kinetics from scRNA-seq data with DeepTXinferrer

This section describes how to recover dynamic burst kinetics from static scRNA-seq data. In principle, once we have constructed the mapping of the parameters to the corresponding stationary distributions through the pre-trained neural network in section 1.3, we can quickly access the corresponding likelihood functions when updating the parameters during the statistical inference process. However, true scRNA-seq data is the result of a combination of gene expression processes and sequencing processes, and the current pre-trained neural network can only yield a description of the gene expression process modeled by TXmodel. Therefore, we require to extent it into a hierarchical model to depict the more realistic generation mechanism behind the scRNA-seq data (***Sarkar A & Stephens M, 2021***).

##### Modeling observed scRNA-seq data with hierarchical model

The hierarchical model aims to couple two different processes, specifically, the true gene expression mechanism process and the sequencing process that immediately follows. In terms of a formula, the probability distribution of observations can be represented as a convolution:

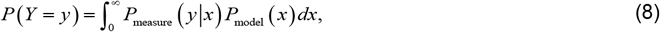

Where *P*_measure_ is the measure model and *P*_model_ is the expression model.

First, we model the sequencing process. Each cell can be regarded as a pool of mRNA, and 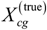 represents the true expression amount of the *g* -th gene in the *c* -th cell. 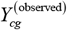 refers to the observed value obtained by scRNA-seq from the given true expression value 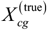, satisfying the following conditional probability distribution

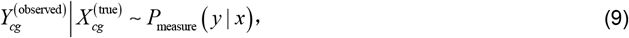

Where *P*_measure_ ( *y* | *x*) characterizes all noise in the sequencing process, and is generally assumed to be Binomial distribution or Poisson distribution, which has been proven experimentally and theoretically (***Sarkar A & Stephens M, 2021; Wang J et al., 2018***). By incorporating an additional sampling probability, denoted as *α*_*cg*_, into the sequencing process, we are able to characterize capture efficiency. We make the assumption that intercellular molecules are independent of one another, and that only proportional products are captured and sequenced, following a Binomial distribution

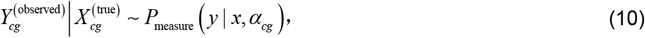

Where *α*_*cg*_ is the sampling probability. Although the experimental capture efficiency ranges from 0.06 to 0.32 **(*Zheng GX et al., 2017; Macosko EZ et al., 2015*)**, we fixed the parameter *α*_*cg*_ = 0.5 to minimize complexity and unidentifiability issues. Further, we demonstrated that inference across different efficiencies (0.5, 0.3, 0.2) consistently yielded strong correlations between inferred burst size and burst frequency (Supplementary Figure S12).

Second, we model the true gene expression process, assuming that it satisfies the following probability distribution

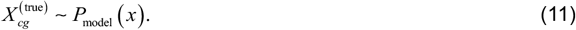

The selection of the true gene expression probability distribution is very critical. It needs to meet the following conditions: (i) It can well characterize the transcription burst kinetics. (ii) The inference derived from this distribution necessitates both efficiency and precision.

In section 1.2, we have constructed a model of the gene expression process with DNA damage, enabling the model’s stationary distribution to encapsulate the bursty dynamics inherent in gene expression. The solution to the model is approximated by a mixed negative binomial distribution, which is obtained from a trained neural network. Importantly, this mixed negative binomial distribution enables efficient inference. Hence,

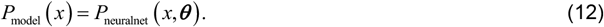

Further, the following hierarchical model can be obtained

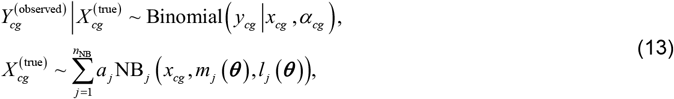

Where 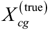 represents the random variable of the true expression value, 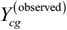 represents the random variable of the observed expression value, Binomial represents the binomial distribution, NB represents negative binomial distribution and *α*_*cg*_ represents the sampling probability. Further, we can obtain

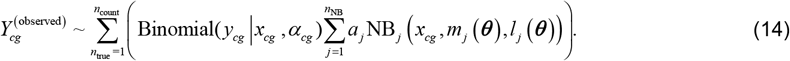

It is proved that 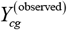 also follows a mixed negative binomial distribution (***Supplementary text 1.3***), denoted as *P*_obsmodel_ .

##### Estimation and optimization process of DeepTXinferrer

We can use the DeepTX framework to infer scRNA-seq data to obtain the corresponding gene expression burst kinetics through optimization methods. Given the expression counts of gene *g* in all cells, its corresponding probability distribution *P*_obsdata_ can be obtained. We use the Hellinger distance to compare the error between the true probability distribution *P*_obsdata_ of genes and the model probability distribution *P*_modeldata_ . The following optimization problem can be obtained

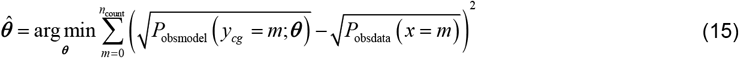

where *n*_count_ is the maximum value of mRNA expression, *P*_obsdata_ is the density probability for each gene *g* . Specifically, *P*_obsdata_ is a function that is differentiable with respect to parameter ***θ***_*g*_ . We can use the gradient descent method to obtain the optimal parameters of the TXmodel and the corresponding distribution of the TXmodel.

##### Posterior distribution

The optimization method in the previous section can obtain the parameters of the mechanism model that satisfies the distribution of scRNA-seq data. In this section, we give a method to solve the confidence interval of each optimized parameter (***Gaskin T et al., 2023***).

Each iteration of the optimization process can obtain the estimated value 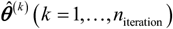 and loss value 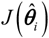 corresponding to the estimated value. In particular, according to Equation (15), the loss is calculated as follows.

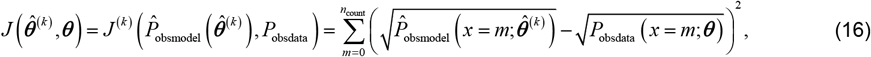

The following posterior probability can be obtained

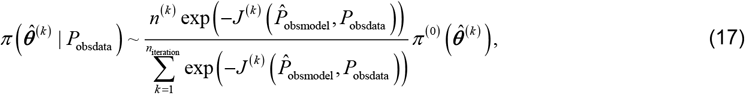

where *π* ^(0)^ is the prior, we take it as uniform density, *n*^(*k*)^ represents the number of values 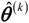 takes during the iteration process, and 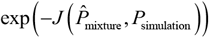 represents the loss potential of each iteration. Let the posterior probability be 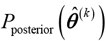, and the marginal distribution corresponding to each component of parameter 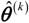 can be obtained

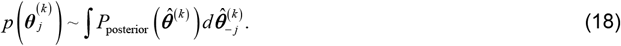

The subscript − *j* means that we integrate the components of ***θ*** ^(*k* )^ except 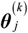 . In particular, to allow the iterative parameters to traverse the entire parameter space, multiple optimizations can be performed with different initial values to improve the performance of the posterior distribution.

##### Validation on synthetic data

We conducted an evaluation of the inference performance of the DeepTX framework on synthetic scRNA-seq data, focusing on three key aspects. (i) Model accuracy: This refers to the minimal discrepancy between the distribution corresponding to the inferred parameters and the input distribution. (ii) Parameter identifiability: This denotes the proximity of the values of the inferred parameters to the true parameters, suggesting that the model can accurately identify the true parameters. (iii) Result robustness: This implies that the outcomes of each inference do not exhibit significant variations.

We created a synthetic dataset 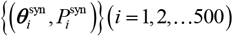 using Sobol sampling and SSA under preset parameters. The synthesized distribution 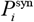 is used as input to the DeepTX framework, and upon optimization, we obtained estimated parameters and their corresponding distribution 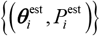. We showed that the Hellinger distance between the estimated and synthetic dataset’s distribution was minimal, indicating accurate estimations (***Supplementary Figure S3a***). Additionally, the burst kinetics of both the synthetic data and the estimated data can be computed utilizing their respective parameters, and 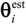 . We showed that a strong correlation was observed between the synthetic burst kinetics ***γ*** ^syn^ = (BF^syn^, BS^syn^ ) and estimated burst kinetics ***γ*** ^est^ = (BF^est^, BS^est^ ) (***Supplementary Figure S3b-c***), suggesting that burst kinetics are identifiable.

Subsequently, we conducted a robustness verification of the framework’s ability to infer burst kinetics. The error was computed as follows:

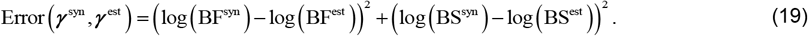

Given a set of parameters 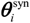, we drew samples of sizes (100, 500, 1000, 1500, 2000) from the probability distribution 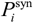 corresponding to 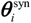 . For each sample size, we performed 50 repetitions, and then DeepTX framework is used to infer the distribution obtained from each sampling, setting different initial values each time. We found that the inferred results are robust, and the robustness increases with the increase in the number of samples (***Supplementary Figure S3d-e***).

### 2 Dataset

#### 2.1 IdU and DMSO dataset

To investigate the effects of 5’-iodo-2’-deoxyuridine (IdU) on genome-wide gene expression, Desai et al. conducted scRNA-seq data on mouse embryonic stem cells, both with and without IdU treatment. Consequently, they obtained scRNA-seq data for 12481 genes of transcriptomes from 812 cells and 13780 genes of transcriptomes from 744 cells, respectively. The authors found a phenomenon that IdU drug treatment increased transcriptional noise genome-wide but did not affect transcriptional mean, further affecting cell fate decisions.

We used the DeepTX framework to infer this set of scRNA-seq data to obtain potential transcription burst kinetics, and further study the impact of changes in transcription burst kinetics on cell fate decisions. To eliminate the impact of technical noise on data inference, we preprocess the data as follows. Firstly, We normalized the data *y*_*cg*_ using Seurat, where *y*_*cg*_ represents the expression level of the gene *g* in the cell *c* . The normalization process is that we multiplied each *y* by a scaling factor 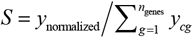, where *y*_normalized_ = 50000 . Particularly, the scaling factor *S* is related to the total expression of genes in each cell. Secondly, after normalizing the expression, we divided it into bins with a unit of 1 to obtain the bin where each expression value is located. Finally, to eliminate the impact of low-expression genes on the inference process, we filter out genes whose average gene expression is less than 1. After data processing, there are 6082 genes from 812 cells left in our IdU and 6082 genes from 744 cells left in DMSO data sets.

#### 2.2 5FU Dataset

To study the impact of 5-fluorouracil (5FU) drug-induced DNA damage on cell fate decisions, Park SR et al. performed scRNA-seq on colon cancer cells treated with 0 μm, 10 μm, and 50 μm 5FU drug (***Park SR et al., 2020***) and obtained 8534 genes from 1673 cells, 7077 genes from 632 cells, and 6661 genes from 619 cells, respectively. The authors found that Colon cancer cells treated with different doses of 5FU exhibit different transcriptional phenotypes, and these different phenotypes correspond to different cell fate decisions.

We used the DeepTX framework to infer these three sets of scRNA-seq data to obtain potential transcriptional burst kinetics, thereby studying the impact of changes in transcriptional burst kinetics caused by different doses of 5FU treatment on cell fate decisions. To eliminate the impact of technical noise on data inference, we preprocess the data in the following steps.

First, we normalize *y*_*cg*_ by multiplying a scale factor *S*, where 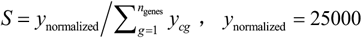 .Second, we divide the normalized data into intervals of unit 1 to round the data. Finally, to eliminate the impact of low-expression genes on the inference process, we filter out genes whose average gene expression is less than 1. After data processing, there are 730 genes from 593 cells left in control data set, 174 genes from 593 cells left in our 10 μm 5FU treatment data set and 187 genes from 564 cells left in 50 μm 5FU treatment data set.

## Additional information

### Funding

This work was supported by National Key R&D Program of China grant 2021YFA1302500 by Natural Science Foundation of P.R. China grants 12171494, 11931019, 11775314, 62373384, and 12301646 by Key-Area Research and Development Program of Guangzhou, P.R. China, grants 202007030004 and 2019B110233002 by Guangdong Basic and Applied Basic Research Foundation grants 2022A1515011540 and 2023A1515011982 by Guangdong Province Key Laboratory of Computational Science at the Sun Yat-sen University grant 2020B1212060032 by the Fundamental Research Funds for the Central Universities, Sun Yat sen University grants 23qnpy48 and 23pnqy49 and by China Postdoctoral Science Foundation grant 2023M734061.

### Author contributions

J.Z., Q.N., and B.J. conceived the project. J.Z., Z.H., and S.L. designed the algorithm. Z.H. and S.L. implemented the code, performed the simulations, analyzed the data, and interpreted the results. Z.W. and Z.Z. analyzed the data and interpreted the results. Q.N. and B.J. supported the data interpretation. Z.H., S.L., and J.Z. wrote the paper. All authors revised and approved the paper. J.Z. supervised the project.

## Additional files

### Supplementary files

Supporting information includes supplementary text S1-S2 and supplementary figure S1-S12.

### Data availability

All the scRNA-seq data can be obtained from the public database. The scRNA-seq data of mouse embryonic stem cells (mESC) + DMSO and mESC + IdU data reported here is available at the Gene Expression Omnibus (GEO) database under accession number: **GSE176044**. The scRNA-seq data of RKO cell with different 5-fluorouracil treatment is available at the Gene Expression Omnibus (GEO) database under accession number: **GSE149224**.

### Code availability

The code for reproducing the presented analysis results is available at GitHub repository (https://github.com/cellfateTX/DeepTX).

## Notes

https://github.com/cellfate/DeepTX

### Competing Interest Statement

The authors have declared no competing interest.

### Summary of Updates

Benchmarked against external methods and updated supplemental files.

## References

López-Maury, L., Marguerat, S., Bähler, J. 2008. Tuning gene expression to changing environments: from rapid responses to evolutionary adaptation. Nature Reviews Genetics 9: 583–593. DOI: 10.1038/nrg2398, PMID: 18591982

Stadhouders, R., Filion, G.J., Graf, T. 2019. Transcription factors and 3D genome conformation in cell-fate decisions. Nature 569: 345–354. DOI: 10.1038/s41586-019-1182-7, PMID: 31092938

Raj, A., van Oudenaarden, A. 2008. Nature, nurture, or chance: stochastic gene expression and its consequences. Cell 135: 216–226. DOI: 10.1016/j.cell.2008.09.050, PMID: 18957198

Balázsi, G., van Oudenaarden, A., Collins, J.J. 2011. Cellular decision making and biological noise: from microbes to mammals. Cell 144: 910–925. DOI: 10.1016/j.cell.2011.01.030, PMID: 21414483

Losick, R., Desplan, C. 2008. Stochasticity and cell fate. Science 320: 65–68. DOI: 10.1126/science.1147888, PMID: 18388284

Eling, N., Morgan, M.D., Marioni, J.C. 2019. Challenges in measuring and understanding biological noise. Nature Reviews Genetics 20: 536–548. DOI: 10.1038/s41576-019-0130-6, PMID: 31114032

Tunnacliffe, E., Chubb, J.R. 2020. What is a transcriptional burst? Trends in Genetics 36: 288–297. DOI: 10.1016/j.tig.2020.01.003, PMID: 32035656

Golding, I., Paulsson, J., Zawilski, S.M., Cox, E.C. 2005. Real-time kinetics of gene activity in individual bacteria. Cell 123: 1025–1036. DOI: 10.1016/j.cell.2005.09.031, PMID: 16360033

Chubb, J.R., Trcek, T., Shenoy, S.M., Singer, R.H. 2006. Transcriptional pulsing of a developmental gene. Current Biology 16: 1018–1025. DOI: 10.1016/j.cub.2006.03.092, PMID: 16713960

Larson, D.R., Zenklusen, D., Wu, B., Chao, J.A., Singer, R.H. 2011. Real-time observation of transcription initiation and elongation on an endogenous yeast gene. Science 332: 475–478. DOI: 10.1126/science.1202142, PMID: 21512033

Lammers, N.C., Kim, Y.J., Zhao, J., Garcia, H.G. 2020. A matter of time: using dynamics and theory to uncover mechanisms of transcriptional bursting. Current Opinion in Cell Biology 67: 147–157. DOI: 10.1016/j.ceb.2020.08.001, PMID: 33242838

Rodriguez, J., Larson, D.R. 2020. Transcription in living Cells: molecular mechanisms of bursting. Annual Review of Biochemistry 89: 189–212. DOI: 10.1146/annurev-biochem-011520-105250, PMID: 32208766

Krenning, L., van den Berg, J., Medema, R.H. 2019. Life or death after a break: what determines the choice? Molecular Cell 76: 346–358. DOI: 10.1016/j.molcel.2019.08.023, PMID: 31561953

Su, T.T. 2006. Cellular responses to DNA damage: one signal, multiple choices. Annual Review of Genetics 40: 187–208. DOI: 10.1146/annurev.genet.40.110405.090428, PMID: 16805666

Hafner, A., Bulyk, M.L., Jambhekar, A., Lahav, G. 2019. The multiple mechanisms that regulate p53 activity and cell fate. Nature Reviews Molecular Cell Biology 20: 199–210. DOI: 10.1038/s41580-019-0110-x, PMID: 30824861

van den Berg, J., Joosten, S.E., Kim, Y., Manjón, A.G., Krenning, L., Koob, L., Feringa, F.M., Klompmaker, R., van den Broek, B., Jalink, K. 2019. DNA end-resection in highly accessible chromatin produces a toxic break. Preprint at arXiv https://doi.org/10.1101/691857. DOI: 10.1101/691857, PMID:

Arora, M., Moser, J., Phadke, H., Basha, A.A., Spencer, S.L. 2017. Endogenous replication stress in mother cells leads to 1uiescence of daughter cells. Cell Reports 19: 1351–1364. DOI: 10.1016/j.celrep.2017.04.055, PMID: 28514656

Barr, A.R., Cooper, S., Heldt, F.S., Butera, F., Stoy, H., Mansfeld, J., Novák, B., Bakal, C. 2017. DNA damage during S-phase mediates the proliferation-quiescence decision in the subsequent G1 via p21 expression. Nature Communications 8: 14728. DOI: 10.1038/ncomms14728, PMID: 28317845

Deng, Z., Shen, D., Yu, M., Zhou, F., Shan, D., Fang, Y., Jin, W., Qian, K., Li, S., Wang, G., Zhang, Y., Ju, L., Xiao, Y., Wang, X. 2023. Pectolinarigenin inhibits bladder urothelial carcinoma cell proliferation by regulating DNA damage/autophagy pathways. Cell Death Discovery 9: 214. DOI: 10.1038/s41420-023-01508-9, PMID: 37393350

Müllers, E., Silva Cascales, H., Jaiswal, H., Saurin, A.T., Lindqvist, A. 2014. Nuclear translocation of Cyclin B1 marks the restriction point for terminal cell cycle exit in G2 phase. Cell Cycle 13: 2733–2743. DOI: 10.4161/15384101.2015.945831, PMID: 25486360

Feringa, F.M., Raaijmakers, J.A., Hadders, M.A., Vaarting, C., Macurek, L., Heitink, L., Krenning, L., Medema, R.H. 2018. Persistent repair intermediates induce senescence. Nature Communications 9: 3923. DOI: 10.1038/s41467-018-06308-9, PMID: 30254262

Toledo, L.I., Murga, M., Gutierrez-Martinez, P., Soria, R., Fernandez-Capetillo, O. 2008. ATR signaling can drive cells into senescence in the absence of DNA breaks. Genes & Development 22: 297–302. DOI: 10.1101/gad.452308, PMID: 18245444

Zhao, Y., Simon, M., Seluanov, A., Gorbunova, V. 2023. DNA damage and repair in age-related inflammation. Nature Reviews Immunology 23: 75–89. DOI: 10.1038/s41577-022-00751-y, PMID: 35831609

Yousefzadeh, M., Henpita, C., Vyas, R., Soto-Palma, C., Robbins, P., Niedernhofer, L. 2021. DNA damage-how and why we age? Elife 10: e62852. DOI: 10.7554/eLife.62852, PMID: 33512317

Carneiro, B.A., El-Deiry, W.S. 2020. Targeting apoptosis in cancer therapy. Nature Reviews Clinical Oncology 17: 395–417. DOI: 10.1038/s41571-020-0341-y, PMID: 32203277

Roos, W.P., Kaina, B. 2013. DNA damage-induced cell death: from specific DNA lesions to the DNA damage response and apoptosis. Cancer Letters 332: 237–248. DOI: 10.1016/j.canlet.2012.01.007, PMID: 22261329

Zheng, P., Chen, Q., Tian, X., Qian, N., Chai, P., Liu, B., Hu, J., Blackstone, C., Zhu, D., Teng, J., Chen, J. 2018. DNA damage triggers tubular endoplasmic reticulum extension to promote apoptosis by facilitating ER-mitochondria signaling. Cell Research 28: 833–854. DOI: 10.1038/s41422-018-0065-z, PMID: 30030520

Lans, H., Hoeijmakers, J.H.J., Vermeulen, W., Marteijn, J.A. 2019. The DNA damage response to transcription stress. Nature Reviews Molecular Cell Biology 20: 766–784. DOI: 10.1038/s41580-019-0169-4, PMID: 31558824

Muñoz, M.J., Pérez Santangelo, M.S., Paronetto, M.P., de la Mata, M., Pelisch, F., Boireau, S., Glover-Cutter, K., Ben-Dov, C., Blaustein, M., Lozano, J.J., Bird, G., Bentley, D., Bertrand, E., Kornblihtt, A.R. 2009. DNA damage regulates alternative splicing through inhibition of RNA polymerase II elongation. Cell 137: 708–720. DOI: 10.1016/j.cell.2009.03.010, PMID: 19450518

Gregersen, L.H., Svejstrup, J.Q. 2018. The cellular response to transcription-blocking DNA damage. Trends in Biochemical Sciences 43: 327–341. DOI: 10.1016/j.tibs.2018.02.010, PMID: 29699641

Geijer, M.E., Marteijn, J.A. 2018. What happens at the lesion does not stay at the lesion: transcription-coupled nucleotide excision repair and the effects of DNA damage on transcription in cis and trans. DNA Repair 71: 56–68. DOI: 10.1016/j.dnarep.2018.08.007, PMID: 30195642

Giono, L.E., Nieto Moreno, N., Cambindo Botto, A.E., Dujardin, G., Muñoz, M.J., Kornblihtt, A.R. 2016. The RNA response to DNA damage. Journal of Molecular Biology 428: 2636–2651. DOI: 10.1016/j.jmb.2016.03.004, PMID: 26979557

Friedrich, D., Friedel, L., Finzel, A., Herrmann, A., Preibisch, S., Loewer, A. 2019. Stochastic transcription in the p53-mediated response to DNA damage is modulated by burst frequency. Molecular Systems Biology 15: e9068. DOI: 10.15252/msb.20199068, PMID: 31885199

Calia, G.P., Chen, X., Zuckerman, B., Weinberger, L. 2023. Comparative analysis between RNA-seq and single-molecule RNA FISH indicates that the pyrimidine nucleobase idoxuridine (IdU) globally amplifies transcriptional noise. Preprint at bioRxiv https://doi.org/10.1101/2023.03.14.532632. DOI: 10.1101/2023.03.14.532632, PMID: 36993609

Desai, R.V., Chen, X., Martin, B., Chaturvedi, S., Hwang, D.W., Li, W., Yu, C., Ding, S., Thomson, M., Singer, R.H., Coleman, R.A., Hansen, M.M.K., Weinberger, L.S. 2021. A DNA repair pathway can regulate transcriptional noise to promote cell fate transitions. Science 373. DOI: 10.1126/science.abc6506, PMID: 34301855

Femino, A.M., Fay, F.S., Fogarty, K., Singer, R.H. 1998. Visualization of single RNA transcripts in situ. Science 280: 585–590. DOI: 10.1126/science.280.5363.585, PMID: 9554849

Qiu, Q., Hu, P., Qiu, X., Govek, K.W., Cámara, P.G., Wu, H. 2020. Massively parallel and time-resolved RNA sequencing in single cells with scNT-seq. Nature Methods 17: 991–1001. DOI: 10.1038/s41592-020-0935-4, PMID: 32868927

Safieddine, A., Coleno, E., Lionneton, F., Traboulsi, A.M., Salloum, S., Lecellier, C.H., Gostan, T., Georget, V., Hassen-Khodja, C., Imbert, A., Mueller, F., Walter, T., Peter, M., Bertrand, E. 2023. HT-smFISH: a cost-effective and flexible workflow for high-throughput single-molecule RNA imaging. Nature Protocols 18: 157–187. DOI: 10.1038/s41596-022-00750-2, PMID: 36280749

Tanay, A., Regev, A. 2017. Scaling single-cell genomics from phenomenology to mechanism. Nature 541: 331–338. DOI: 10.1038/nature21350, PMID: 28102262

Picelli, S., Björklund Å, K., Faridani, O.R., Sagasser, S., Winberg, G., Sandberg, R. 2013. Smart-seq2 for sensitive full-length transcriptome profiling in single cells. Nature Methods 10: 1096–1098. DOI: 10.1038/nmeth.2639, PMID: 24056875

Zheng, G.X., Terry, J.M., Belgrader, P., Ryvkin, P., Bent, Z.W., Wilson, R., Ziraldo, S.B., Wheeler, T.D., McDermott, G.P., Zhu, J., Gregory, M.T., Shuga, J., Montesclaros, L., Underwood, J.G., Masquelier, D.A., Nishimura, S.Y., Schnall-Levin, M., Wyatt, P.W., Hindson, C.M., Bharadwaj, R., Wong, A., Ness, K.D., Beppu, L.W., Deeg, H.J., McFarland, C., Loeb, K.R., Valente, W.J., Ericson, N.G., Stevens, E.A., Radich, J.P., Mikkelsen, T.S., Hindson, B.J., Bielas, J.H. 2017. Massively parallel digital transcriptional profiling of single cells. Nature Communications 8: 14049. DOI: 10.1038/ncomms14049, PMID: 28091601

Faure, A.J., Schmiedel, J.M., Lehner, B. 2017. Systematic analysis of the determinants of gene expression noise in embryonic stem cells. Cell Systems 5: 471-484.e474. DOI: 10.1016/j.cels.2017.10.003, PMID: 29102610

Morgan, M.D., Marioni, J.C. 2018. CpG island composition differences are a source of gene expression noise indicative of promoter responsiveness. Genome Biology 19: 81. DOI: 10.1186/s13059-018-1461-x, PMID: 29945659

Ochiai, H., Hayashi, T., Umeda, M., Yoshimura, M., Harada, A., Shimizu, Y., Nakano, K., Saitoh, N., Liu, Z., Yamamoto, T., Okamura, T., Ohkawa, Y., Kimura, H., Nikaido, I. 2020. Genome-wide kinetic properties of transcriptional bursting in mouse embryonic stem cells. Science Advances 6: eaaz6699. DOI: 10.1126/sciadv.aaz6699, PMID: 32596448

Larsson, A.J.M., Johnsson, P., Hagemann-Jensen, M., Hartmanis, L., Faridani, O.R., Reinius, B., Segerstolpe, A., Rivera, C.M., Ren, B., Sandberg, R. 2019. Genomic encoding of transcriptional burst kinetics. Nature 565: 251–254. DOI: 10.1038/s41586-018-0836-1, PMID: 30602787

Luo, S., Wang, Z., Zhang, Z., Zhou, T., Zhang, J. 2023. Genome-wide inference reveals that feedback regulations constrain promoter-dependent transcriptional burst kinetics. Nucleic Acids Research 51: 68–83. DOI: 10.1093/nar/gkac1204, PMID: 36583343

Luo, S., Zhang, Z., Wang, Z., Yang, X., Chen, X., Zhou, T., Zhang, J. 2023. Inferring transcriptional bursting kinetics from single-cell snapshot data using a generalized telegraph model. Royal Society Open Science 10: 221057. DOI: 10.1098/rsos.221057, PMID: 37035293

Stumpf, P.S., Smith, R.C., Lenz, M., Schuppert, A., Müller, F.-J., Babtie, A., Chan, T.E., Stumpf, M.P., Please, C.P., Howison, S.D. 2017. Stem cell differentiation as a non-Markov stochastic process. Cell Systems 5: 268-282. e267. DOI: 10.1016/j.cels.2017.08.009, PMID: 28957659

Voss, T.C., Hager, G.L. 2014. Dynamic regulation of transcriptional states by chromatin and transcription factors. Nature Reviews Genetics 15: 69–81. DOI: 10.1038/nrg3623, PMID: 24342920

Singh, A., Vargas, C.A., Karmakar, R. 2013. Stochastic analysis and inference of a two-state genetic promoter model. Proceedings of the American Control Conference: 4563–4568. DOI:

Cavallaro, M., Walsh, M.D., Jones, M., Teahan, J., Tiberi, S., Finkenstadt, B., Hebenstreit, D. 2021. 3 (‘)-5 (‘) crosstalk contributes to transcriptional bursting. Genome Biology 22: 56. DOI: 10.1186/s13059-020-02227-5, PMID: 33541397

Zhang, J., Chen, L., Zhou, T. 2012. Analytical distribution and tunability of noise in a model of promoter progress. Biophysical Journal 102: 1247–1257. DOI: 10.1016/j.bpj.2012.02.001, PMID: 22455907

Zhang, J., Zhou, T. 2014. Promoter-mediated transcriptional dynamics. Biophysical Journal 106: 479–488. DOI: 10.1016/j.bpj.2013.12.011, PMID: 24461023

Zhou, T., Zhang, J. 2012. Analytical results for a multistate gene model. SIAM Journal On Applied Mathmatics 72: 789–818. DOI: 10.1137/110852887, xPMID:

Rodriguez, J., Ren, G., Day, C.R., Zhao, K., Chow, C.C., Larson, D.R. 2019. Intrinsic dynamics of a human gene reveal the basis of expression heterogeneity. Cell 176: 213-226.e218. DOI: 10.1016/j.cell.2018.11.026, PMID: 30554876

Zoller, B., Nicolas, D., Molina, N., Naef, F. 2015. Structure of silent transcription intervals and noise characteristics of mammalian genes. Molecular Systems Biology 11: 823. DOI: 10.15252/msb.20156257, PMID: 26215071

Daigle, B.J., Jr., Soltani, M., Petzold, L.R., Singh, A. 2015. Inferring single-cell gene expression mechanisms using stochastic simulation. Bioinformatics 31: 1428–1435. DOI: 10.1093/bioinformatics/btv007, PMID: 25573914

Kumar, N., Singh, A., Kulkarni, R.V. 2015. Transcriptional bursting in gene expression: analytical results for general stochastic models. PLoS Computational Biology 11: e1004292. DOI: 10.1371/journal.pcbi.1004292, PMID: 26474290

Schwabe, A., Rybakova, K.N., Bruggeman, F.J. 2012. Transcription stochasticity of complex gene regulation models. Biophysical Journal 103: 1152–1161. DOI: 10.1016/j.bpj.2012.07.011, PMID: 22995487

Stinchcombe, A.R., Peskin, C.S., Tranchina, D. 2012. Population density approach for discrete mRNA distributions in generalized switching models for stochastic gene expression. Physical Review E: Statistical Nonlinear and Soft Matter Physics 85: 061919. DOI: 10.1103/PhysRevE.85.061919, PMID: 23005139

Zhang, J., Zhou, T. 2019. Markovian approaches to modeling intracellular reaction processes with molecular memory. Proceedings of the National Academy of Sciences 116: 23542–23550. DOI: 10.1073/pnas.1913926116, PMID: 31685609

Zhang, J.J., Zhou, T.S. 2019. Stationary moments, distribution conjugation and phenotypic regions in stochastic gene transcription. Mathematical Biosciences and Engineering 16: 6134–6166. DOI: 10.3934/mbe.2019307, PMID: 31499756

Sisson, S.A., Fan, Y., Tanaka, M.M. 2007. Sequential Monte Carlo without likelihoods. Proceedings of the National Academy of Sciences 104: 1760–1765. DOI: 10.1073/pnas.0607208104, PMID: 17264216

Gómez-Schiavon, M., Chen, L.F., West, A.E., Buchler, N.E. 2017. BayFish: Bayesian inference of transcription dynamics from population snapshots of single-molecule RNA FISH in single cells. Genome Biology 18: 164. DOI: 10.1186/s13059-017-1297-9, PMID: 28870226

Jiang, Q., Fu, X., Yan, S., Li, R., Du, W., Cao, Z., Qian, F., Grima, R. 2021. Neural network aided approximation and parameter inference of non-Markovian models of gene expression. Nature Communications 12: 2618. DOI: 10.1038/s41467-021-22919-1, PMID: 33976195

Wang, S., Fan, K., Luo, N., Cao, Y., Wu, F., Zhang, C., Heller, K.A., You, L. 2019. Massive computational acceleration by using neural networks to emulate mechanism-based biological models. Nature Communications 10: 4354. DOI: 10.1038/s41467-019-12342-y, PMID: 31554788

Michoski, C., Milosavljević, M., Oliver, T., Hatch, D.R. 2020. Solving differential equations using deep neural networks. Neurocomputing 399: 193–212. DOI: 10.1016/j.neucom.2020.02.015, PMID:

Davis, C.N., Hollingsworth, T.D., Caudron, Q., Irvine, M.A. 2020. The use of mixture density networks in the emulation of complex epidemiological individual-based models. PLoS Computational Biology 16: e1006869. DOI: 10.1371/journal.pcbi.1006869, PMID:

Tang, Y., Weng, J., Zhang, P. 2023. Neural-network solutions to stochastic reaction networks. Nature Machine Intelligence 5: 376–385. DOI: 10.1038/s42256-023-00632-6, PMID:

Gaskin, T., Pavliotis, G.A., Girolami, M. 2023. Neural parameter calibration for large-scale multiagent models. Proceedings of the National Academy of Sciences 120: e2216415120. DOI: 10.1073/pnas.2216415120, PMID: 36763529

Sukys, A., Öcal, K., Grima, R. 2022. Approximating solutions of the chemical master equation using neural networks. iScience 25: 105010. DOI: 10.1016/j.isci.2022.105010, PMID: 36117994

Tang, W., Jørgensen, A.C.S., Marguerat, S., Thomas, P., Shahrezaei, V. 2023. Modelling capture efficiency of single cell RNA-sequencing data improves inference of transcriptome-wide burst kinetics. Preprint at bioRxiv https://doi.org/10.1101/2023.1103.1106.531327. DOI: 10.1101/2023.03.06.531327, PMID:

Gupta, A., Schwab, C., Khammash, M. 2021. DeepCME: a deep learning framework for computing solution statistics of the chemical master equation. PLoS Computational Biology 17: e1009623. DOI: 10.1371/journal.pcbi.1009623, PMID: 34879062

Perez-Carrasco, R., Beentjes, C., Grima, R. 2020. Effects of cell cycle variability on lineage and population measurements of messenger RNA abundance. Journal of the Royal Society Interface 17: 20200360. DOI: 10.1098/rsif.2020.0360, PMID: 32634365

Öcal, K., Gutmann, M.U., Sanguinetti, G., Grima, R. 2022. Inference and uncertainty quantification of stochastic gene expression via synthetic models. Journal of the Royal Society Interface 19: 20220153. DOI: 10.1098/rsif.2022.0153, PMID: 35858045

Sobol, I.M. 1967. On the distribution of points in a cube and the approximate evaluation of integrals. USSR Comput. Math. Math. Phys. 7: 784–802. DOI: 10.1016/0041-5553(67)90144-9, PMID:

Zhang, Z., Deng, Q., Wang, Z., Chen, Y., Zhou, T. 2021. Exact results for queuing models of stochastic transcription with memory and crosstalk. Physical Review E 103: 062414. DOI: 10.1103/PhysRevE.103.062414, PMID: 34271765

Zhang, J., Chen, A., Qiu, H., Zhang, J., Tian, T., Zhou, T. 2024. Exact results for gene-expression models with general waiting-time distributions. Physical Review E 109: 024119. DOI: 10.1103/PhysRevE.109.024119, PMID: 38491572

Gupta, P.B., Fillmore, C.M., Jiang, G., Shapira, S.D., Tao, K., Kuperwasser, C., Lander, E.S. 2011. Stochastic state transitions give rise to phenotypic equilibrium in populations of cancer cells. Cell 146: 633–644. DOI: 10.1016/j.cell.2011.07.026, PMID: 21854987

Cohen, A.A., Geva-Zatorsky, N., Eden, E., Frenkel-Morgenstern, M., Issaeva, I., Sigal, A., Milo, R., Cohen-Saidon, C., Liron, Y., Kam, Z., Cohen, L., Danon, T., Perzov, N., Alon, U. 2008. Dynamic proteomics of individual cancer cells in response to a drug. Science 322: 1511–1516. DOI: 10.1126/science.1160165, PMID: 19023046

Bessarabova, M., Kirillov, E., Shi, W., Bugrim, A., Nikolsky, Y., Nikolskaya, T. 2010. Bimodal gene expression patterns in breast cancer. BMC Genomics 11 (Suppl 1): S8. DOI: 10.1186/1471-2164-11-s1-s8, PMID: 20158879

Sarkar, A., Stephens, M. 2021. Separating measurement and expression models clarifies confusion in single-cell RNA sequencing analysis. Nature Genetics 53: 770–777. DOI: 10.1038/s41588-021-00873-4, PMID: 34031584

Das, S., Suganthan, P.N. 2010. Differential evolution: A survey of the state-of-the-art. IEEE transactions on evolutionary computation 15: 4–31. DOI: 10.1109/TEVC.2010.2059031, PMID:

Tang, W., Jørgensen, A.C.S., Marguerat, S., Thomas, P., Shahrezaei, V. 2023. Modelling capture efficiency of single-cell RNA-sequencing data improves inference of transcriptome-wide burst kinetics. Bioinformatics 39. DOI: 10.1093/bioinformatics/btad395, PMID: 37354494

Gu, J., Laszik, N., Miles, C.E., Allard, J., Downing, T.L., Read, E.L. 2025. Scalable inference and identifiability of kinetic parameters for transcriptional bursting from single cell data. Bioinformatics. DOI: 10.1093/bioinformatics/btaf581, PMID: 41131798

Li, C., Xu, X., Chen, S., Xu, A., Guan, T., Wu, H., Pei, D., Liu, J. 2024. Epigenetic reshaping through damage: promoting cell fate transition by BrdU and IdU incorporation. Cell and Bioscience 14: 9. DOI: 10.1186/s13578-024-01192-x, PMID: 38229206

Newman, J.R., Ghaemmaghami, S., Ihmels, J., Breslow, D.K., Noble, M., DeRisi, J.L., Weissman, J.S. 2006. Single-cell proteomic analysis of S. cerevisiae reveals the architecture of biological noise. Nature 441: 840–846. DOI: 10.1038/nature04785, PMID: 16699522

Consortium, G.O. 2015. Gene Ontology Consortium: going forward. Nucleic Acids Research 43: D1049–1056. DOI: 10.1093/nar/gku1179, PMID: 25428369

Smits, V.A., Medema, R.H. 2001. Checking out the G2/M transition. Biochimica et Biophysica Acta (BBA)-Gene Structure and Expression 1519: 1–12. DOI:

Subramanian, A., Tamayo, P., Mootha, V.K., Mukherjee, S., Ebert, B.L., Gillette, M.A., Paulovich, A., Pomeroy, S.L., Golub, T.R., Lander, E.S., Mesirov, J.P. 2005. Gene set enrichment analysis: a knowledge-based approach for interpreting genome-wide expression profiles. Proceedings of the National Academy of Sciences 102: 15545–15550. DOI: 10.1073/pnas.0506580102, PMID: 16199517

Rosina, M., Langone, F., Giuliani, G., Cerquone Perpetuini, A., Reggio, A., Calderone, A., Fuoco, C., Castagnoli, L., Gargioli, C., Cesareni, G. 2019. Osteogenic differentiation of skeletal muscle progenitor cells is activated by the DNA damage response. Scientific Reports 9: 5447. DOI: 10.1038/s41598-019-41926-3, PMID: 30931986

Riccio, F., Micarelli, E., Secci, R., Giuliani, G., Vumbaca, S., Massacci, G., Castagnoli, L., Fuoco, C., Cesareni, G. 2022. Transcription Factor Activation Profiles (TFAP) identify compounds promoting differentiation of Acute Myeloid Leukemia cell lines. Cell Death Discovery 8: 16. DOI: 10.1038/s41420-021-00811-7, PMID: 35013135

Liu, Z., Wang, L., Welch, J.D., Ma, H., Zhou, Y., Vaseghi, H.R., Yu, S., Wall, J.B., Alimohamadi, S., Zheng, M., Yin, C., Shen, W., Prins, J.F., Liu, J., Qian, L. 2017. Single-cell transcriptomics reconstructs fate conversion from fibroblast to cardiomyocyte. Nature 551: 100–104. DOI: 10.1038/nature24454, PMID: 29072293

Kuipers, E.J., Grady, W.M., Lieberman, D., Seufferlein, T., Sung, J.J., Boelens, P.G., van de Velde, C.J., Watanabe, T. 2015. Colorectal cancer. Nature Reviews Disease Primers 1: 15065. DOI: 10.1038/nrdp.2015.65, PMID: 27189416

Bunz, F., Hwang, P.M., Torrance, C., Waldman, T., Zhang, Y., Dillehay, L., Williams, J., Lengauer, C., Kinzler, K.W., Vogelstein, B. 1999. Disruption of p53 in human cancer cells alters the responses to therapeutic agents. Journal of Clinical Investigation 104: 263–269. DOI: 10.1172/jci6863, PMID: 10430607

Chang, G.S., Chen, X.A., Park, B., Rhee, H.S., Li, P., Han, K.H., Mishra, T., Chan-Salis, K.Y., Li, Y., Hardison, R.C., Wang, Y., Pugh, B.F. 2014. A comprehensive and high-resolution genome-wide response of p53 to stress. Cell Reports 8: 514–527. DOI: 10.1016/j.celrep.2014.06.030, PMID: 25043190

Park, S.R., Namkoong, S., Friesen, L., Cho, C.S., Zhang, Z.Z., Chen, Y.C., Yoon, E., Kim, C.H., Kwak, H., Kang, H.M., Lee, J.H. 2020. Single-cell transcriptome analysis of colon cancer cell response to 5-fluorouracil-induced DNA damage. Cell Reports 32: 108077. DOI: 10.1016/j.celrep.2020.108077, PMID: 32846134

Hongmei, Z. 2012, Apoptosis and medicine. InTechOpen.

Sun, Y., Rigas, B. 2008. The thioredoxin system mediates redox-induced cell death in human colon cancer cells: implications for the mechanism of action of anticancer agents. Cancer Research 68: 8269–8277. DOI: 10.1158/0008-5472.Can-08-2010, PMID: 18922898

Laha, D., Pramanik, A., Chattopadhyay, S., kumar Dash, S., Roy, S., Pramanik, P., Karmakar, P. 2015. Folic acid modified copper oxide nanoparticles for targeted delivery in in vitro and in vivo systems. RSC Advances 5: 68169–68178. DOI: 10.1039/C5RA08110F, PMID:

Bwatanglang, I.B., Mohammad, F., Yusof, N.A., Abdullah, J., Alitheen, N.B., Hussein, M.Z., Abu, N., Mohammed, N.E., Nordin, N., Zamberi, N.R., Yeap, S.K. 2016. In vivo tumor targeting and anti-tumor effects of 5-fluororacil loaded, folic acid targeted quantum dot system. Journal of Colloid and Interface Science 480: 146–158. DOI: 10.1016/j.jcis.2016.07.011, PMID: 27428851

Chenna, S., Koopman, W.J.H., Prehn, J.H.M., Connolly, N.M.C. 2022. Mechanisms and mathematical modeling of ROS production by the mitochondrial electron transport chain. American Journal of Physiology: Cell Physiology 323: C69–c83. DOI: 10.1152/ajpcell.00455.2021, PMID: 35613354

Adam-Vizi, V. 2005. Production of reactive oxygen species in brain mitochondria: contribution by electron transport chain and non-electron transport chain sources. Antioxidants & Redox Signaling 7: 1140–1149. DOI: 10.1089/ars.2005.7.1140, PMID: 16115017

Shokolenko, I.N., Wilson, G.L., Alexeyev, M.F. 2014. Aging: a mitochondrial DNA perspective, critical analysis and an update. World Journal of Experimental Medicine 4: 46–57. DOI: 10.5493/wjem.v4.i4.46, PMID: 25414817

Handali, S., Moghimipour, E., Rezaei, M., Ramezani, Z., Kouchak, M., Amini, M., Angali, K.A., Saremy, S., Dorkoosh, F.A. 2018. A novel 5-Fluorouracil targeted delivery to colon cancer using folic acid conjugated liposomes. Biomedicine & Pharmacotherapy 108: 1259–1273. DOI: 10.1016/j.biopha.2018.09.128, PMID: 30372827

Hwang, P.M., Bunz, F., Yu, J., Rago, C., Chan, T.A., Murphy, M.P., Kelso, G.F., Smith, R.A., Kinzler, K.W., Vogelstein, B. 2001. Ferredoxin reductase affects p53-dependent, 5-fluorouracil-induced apoptosis in colorectal cancer cells. Nature Medicine 7: 1111–1117. DOI: 10.1038/nm1001-1111, PMID: 11590433

Kuranaga, N., Shinomiya, N., Mochizuki, H. 2001. Long-term cultivation of colorectal carcinoma cells with anticancer drugs induces drug resistance and telomere elongation: an in vitro study. BMC Cancer 1: 10. DOI: 10.1186/1471-2407-1-10, PMID: 11518543

Nabetani, A., Ishikawa, F. 2011. Alternative lengthening of telomeres pathway: recombination-mediated telomere maintenance mechanism in human cells. J Biochem 149: 5–14. DOI: 10.1093/jb/mvq119, PMID: 20937668

Cristofari, G., Adolf, E., Reichenbach, P., Sikora, K., Terns, R.M., Terns, M.P., Lingner, J. 2007. Human telomerase RNA accumulation in Cajal bodies facilitates telomerase recruitment to telomeres and telomere elongation. Molecular Cell 27: 882–889. DOI: 10.1016/j.molcel.2007.07.020, PMID: 17889662

Lipinska, N., Romaniuk, A., Paszel-Jaworska, A., Toton, E., Kopczynski, P., Rubis, B. 2017. Telomerase and drug resistance in cancer. Cellular and Molecular Life Sciences 74: 4121–4132. DOI: 10.1007/s00018-017-2573-2, PMID: 28623509

Wiecek, A.J., Cutty, S.J., Kornai, D., Parreno-Centeno, M., Gourmet, L.E., Tagliazucchi, G.M., Jacobson, D.H., Zhang, P., Xiong, L., Bond, G.L., Barr, A.R., Secrier, M. 2023. Genomic hallmarks and therapeutic implications of G0 cell cycle arrest in cancer. Genome Biology 24: 128. DOI: 10.1186/s13059-023-02963-4, PMID: 37221612

Hänzelmann, S., Castelo, R., Guinney, J. 2013. GSVA: gene set variation analysis for microarray and RNA-seq data. BMC Bioinformatics 14: 7. DOI: 10.1186/1471-2105-14-7, PMID: 23323831

Dar, R.D., Razooky, B.S., Singh, A., Trimeloni, T.V., McCollum, J.M., Cox, C.D., Simpson, M.L., Weinberger, L.S. 2012. Transcriptional burst frequency and burst size are equally modulated across the human genome. Proceedings of the National Academy of Sciences 109: 17454–17459. DOI: 10.1073/pnas.1213530109, PMID: 23064634

Zenklusen, D., Larson, D.R., Singer, R.H. 2008. Single-RNA counting reveals alternative modes of gene expression in yeast. Nature Structural & Molecular Biology 15: 1263–1271. DOI: 10.1038/nsmb.1514, PMID: 19011635

Phillips, R., Kondev, J., Theriot, J., Garcia, H. 2012. Physical Biology of the Cell. Garland Science.

Lomvardas, S., Thanos, D. 2002. Opening chromatin. Molecular Cell 9: 209–211. DOI: 10.1016/s1097-2765(02)00463-x, PMID: 11864594

Fuda, N.J., Ardehali, M.B., Lis, J.T. 2009. Defining mechanisms that regulate RNA polymerase II transcription in vivo. Nature 461: 186–192. DOI: 10.1038/nature08449, PMID: 19741698

Nordick, B., Yu, P.Y., Liao, G., Hong, T. 2022. Nonmodular oscillator and switch based on RNA decay drive regeneration of multimodal gene expression. Nucleic Acids Research 50: 3693–3708. DOI: 10.1093/nar/gkac217, PMID: 35380686

Maizels, R.J. 2024. A dynamical perspective: moving towards mechanism in single-cell transcriptomics. Philos Trans R Soc Lond B Biol Sci 379: 20230049. DOI: 10.1098/rstb.2023.0049, PMID: 38432314

Sukys, A., Grima, R. 2025. Cell-cycle dependence of bursty gene expression: insights from fitting mechanistic models to single-cell RNA-seq data. Nucleic Acids Research 53. DOI: 10.1093/nar/gkaf295, PMID: 40240003

Garrido-Rodriguez, M., Lopez-Lopez, D., Ortuno, F.M., Peña-Chilet, M., Muñoz, E., Calzado, M.A., Dopazo, J. 2021. A versatile workflow to integrate RNA-seq genomic and transcriptomic data into mechanistic models of signaling pathways. PLoS Computational Biology 17: e1008748. DOI: 10.1371/journal.pcbi.1008748, PMID: 33571195

Wan, Y., Anastasakis, D.G., Rodriguez, J., Palangat, M., Gudla, P., Zaki, G., Tandon, M., Pegoraro, G., Chow, C.C., Hafner, M., Larson, D.R. 2021. Dynamic imaging of nascent RNA reveals general principles of transcription dynamics and stochastic splice site selection. Cell 184: 2878-2895.e2820. DOI: 10.1016/j.cell.2021.04.012, PMID: 33979654

Gorin, G., Pachter, L. 2022. Modeling bursty transcription and splicing with the chemical master equation. Biophysical Journal 121: 1056–1069. DOI: 10.1016/j.bpj.2022.02.004, PMID: 35143775

Golan-Lavi, R., Giacomelli, C., Fuks, G., Zeisel, A., Sonntag, J., Sinha, S., Köstler, W., Wiemann, S., Korf, U., Yarden, Y., Domany, E. 2017. Coordinated pulses of mRNA and of protein translation or degradation produce EGF-Induced protein bursts. Cell Reports 18: 3129–3142. DOI: 10.1016/j.celrep.2017.03.014, PMID: 28355565

Nicolas, D., Zoller, B., Suter, D.M., Naef, F. 2018. Modulation of transcriptional burst frequency by histone acetylation. Proceedings of the National Academy of Sciences 115: 7153–7158. DOI: 10.1073/pnas.1722330115, PMID: 29915087

Liu, T., Zhang, J., Zhou, T. 2016. Effect of Interaction between chromatin loops on cell-to-cell variability in gene expression. PLoS Computational Biology 12: e1004917. DOI: 10.1371/journal.pcbi.1004917, PMID: 27153118

Wang, Z., Zhang, Z., Luo, S., Zhou, T., Zhang, J. 2024. Power-law behavior of transcriptional bursting regulated by enhancer-promoter communication. Genome Research 34: 106–118. DOI: 10.1101/gr.278631.123, PMID: 38171575

Wang, Z., Luo, S., Zhang, Z., Zhou, T., Zhang, J. 2023. 4D nucleome equation predicts gene expression controlled by long-range enhancer-promoter interaction. PLoS Computational Biology 19: e1011722. DOI: 10.1371/journal.pcbi.1011722, PMID: 38109463

Kepler, T.B., Elston, T.C. 2001. Stochasticity in transcriptional regulation: origins, consequences, and mathematical representations. Biophysical Journal 81: 3116–3136. DOI: 10.1016/s0006-3495(01)75949-8, PMID: 11720979

Hope, B., Kosofsky, B., Hyman, S.E., Nestler, E.J. 1992. Regulation of immediate early gene expression and AP-1 binding in the rat nucleus accumbens by chronic cocaine. Proc Natl Acad Sci U S A 89: 5764–5768. DOI: 10.1073/pnas.89.13.5764, PMID: 1631058

Wen, L., Tang, F. 2022. Recent advances in single-cell sequencing technologies. Precision Clinical Medicine 5: pbac002. DOI: 10.1093/pcmedi/pbac002, PMID: 35821681

Miao, Z., Humphreys, B.D., McMahon, A.P., Kim, J. 2021. Multi-omics integration in the age of million singlecell data. Nature Reviews Neurology 17: 710–724. DOI: 10.1038/s41581-021-00463-x, PMID: 34417589

Ramsköld, D., Hendriks, G.J., Larsson, A.J.M., Mayr, J.V., Ziegenhain, C., Hagemann-Jensen, M., Hartmanis, L., Sandberg, R. 2024. Single-cell new RNA sequencing reveals principles of transcription at the resolution of individual bursts. Nature Cell Biology 26: 1725–1733. DOI: 10.1038/s41556-024-01486-9, PMID: 39198695

Gupta, A., Martin-Rufino, J.D., Jones, T.R., Subramanian, V., Qiu, X., Grody, E.I., Bloemendal, A., Weng, C., Niu, S.Y., Min, K.H., Mehta, A., Zhang, K., Siraj, L., Al’ Khafaji, A., Sankaran, V.G., Raychaudhuri, S., Cleary, B., Grossman, S., Lander, E.S. 2022. Inferring gene regulation from stochastic transcriptional variation across single cells at steady state. Proceedings of the National Academy of Sciences 119: e2207392119. DOI: 10.1073/pnas.2207392119, PMID: 35969771

Gorin, G., Carilli, M., Chari, T., Pachter, L. 2024. Spectral neural approximations for models of transcriptional dynamics. Biophysical Journal 123: 2892–2901. DOI: 10.1016/j.bpj.2024.04.034, PMID: 38715358

Kuntz, J., Thomas, P., Stan, G.-B., Barahona, M. 2021. Stationary distributions of continuous-time Markov chains: a review of theory and truncation-based approximations. SIAM Review 63: 3–64. DOI:

Peccoud, J., Ycart, B. 1995. Markovian modeling of gene-product synthesis. Theoretical Population Biology 48: 222–234. DOI: 10.1006/tpbi.1995.1027, PMID:

Kim, J.K., Marioni, J.C. 2013. Inferring the kinetics of stochastic gene expression from single-cell RNA-sequencing data. Genome Biology 14: R7. DOI: 10.1186/gb-2013-14-1-r7, PMID: 23360624

Zoller, B., Little, S.C., Gregor, T. 2018. Diverse spatial expression patterns emerge from unified kinetics of transcriptional bursting. Cell 175: 835-847.e825. DOI: 10.1016/j.cell.2018.09.056, PMID: 30340044

Hoppe, C., Bowles, J.R., Minchington, T.G., Sutcliffe, C., Upadhyai, P., Rattray, M., Ashe, H.L. 2020. Modulation of the promoter activation rate dictates the transcriptional response to graded BMP signaling levels in the drosophila embryo. Dev Cell 54: 727-741.e727. DOI: 10.1016/j.devcel.2020.07.007, PMID: 32758422

Boguñá, M., Lafuerza, L.F., Toral, R., Serrano, M. 2014. Simulating non-Markovian stochastic processes. Physical Review E: Statistical Nonlinear and Soft Matter Physics 90: 042108. DOI: 10.1103/PhysRevE.90.042108, PMID: 25375439

Masuda, N., Rocha, L.E. 2018. A Gillespie algorithm for non-Markovian stochastic processes. SIAM Review 60: 95–115. DOI: 10.1137/16M1055876, PMID:

Bar-Sinai, Y., Hoyer, S., Hickey, J., Brenner, M.P. 2019. Learning data-driven discretizations for partial differential equations. Proceedings of the National Academy of Sciences 116: 15344–15349. DOI: 10.1073/pnas.1814058116, PMID: 31311866

Risso, D., Perraudeau, F., Gribkova, S., Dudoit, S., Vert, J.P. 2018. A general and flexible method for signal extraction from single-cell RNA-seq data. Nature Communications 9: 284. DOI: 10.1038/s41467-017-02554-5, PMID: 29348443

Love, M.I., Huber, W., Anders, S. 2014. Moderated estimation of fold change and dispersion for RNA-seq data with DESeq2. Genome Biology 15: 550. DOI: 10.1186/s13059-014-0550-8, PMID: 25516281

Kingma, D.P., Ba, J. 2014. Adam: a method for stochastic optimization. Preprint at arXiv https://doi.org/10.48550/arXiv.1412.6980. DOI: 10.48550/arXiv.1412.6980, PMID:

Goodfellow, I., Bengio, Y., Courville, A. 2016. Deep Learning. MIT press.

Glorot, X., Bengio, Y. 2010. Understanding the difficulty of training deep feedforward neural networks. Proceedings of the thirteenth international conference on artificial intelligence and statistics: 249–256. DOI:

Keskar, N.S., Mudigere, D., Nocedal, J., Smelyanskiy, M., Tang, P.T.P. 2016. On large-batch training for deep learning: generalization gap and sharp minima. Preprint at arXiv https://arxiv.org/abs/1609.04836. DOI:

Wang, J., Huang, M., Torre, E., Dueck, H., Shaffer, S., Murray, J., Raj, A., Li, M., Zhang, N.R. 2018. Gene expression distribution deconvolution in single-cell RNA sequencing. Proceedings of the National Academy of Sciences 115: E6437–E6446. DOI: 10.1073/pnas.1721085115, PMID: 29946020

Macosko, E.Z., Basu, A., Satija, R., Nemesh, J., Shekhar, K., Goldman, M., Tirosh, I., Bialas, A.R., Kamitaki, N., Martersteck, E.M., Trombetta, J.J., Weitz, D.A., Sanes, J.R., Shalek, A.K., Regev, A., McCarroll, S.A. 2015. Highly parallel genome-wide expression profiling of individual cells using nanoliter dsroplets. Cell 161: 1202–1214. DOI: 10.1016/j.cell.2015.05.002, PMID: 26000488

