## Supplemental File for "Deep learning linking mechanistic models to single-cell transcriptomics data reveals transcriptional bursting in response to DNA damage"

### 1. DeepTX Framework

#### 1.1 Modelling gene expression processes in DNA damage

Generally, gene expression switches between on and off states, and the two-state model can model this gene expression process. However, DNA damage will cause multiple different transcription rates during DNA transcription, that is, multiple steps. Therefore, we use a more general model of gene transcription (TX) to model the transcription process, taking DNA damage into account. We first describe the classic telegraph model (CTM), which is a specific case of the TX model.

The CTM model is widely used model for capturing Markovian gene expression burst kinetics. It describes stochastic gene expression as a sequence of four biochemical reactions involving two gene states (ON and OFF), mRNA transcription and degradation:

$$\begin{aligned}&R_1 : \mathrm{OFF} \xrightarrow{k_{\mathrm{off}}} \mathrm{ON},\ R_2 : \mathrm{ON} \xrightarrow{k_{\mathrm{on}}} \mathrm{OFF}\ ,\\ &R_3 : \mathrm{ON} \xrightarrow{k_{\mathrm{syn}}} \mathrm{ON} + \mathrm{mRNA},\ R_4 : \mathrm{mRNA} \xrightarrow{k_{\mathrm{deg}}} \varnothing,\end{aligned} \tag{1}$$

$k_{\mathrm{off}}$ as the rate at which the gene switches from OFF to ON, $k_{\mathrm{on}}$ as the rate at which the gene switches from ON to OFF, $k_{\mathrm{syn}}$ as the rate of mRNA synthesis and $k_{\mathrm{deg}}$ as the rate of mRNA degradation. In this model, gene switching between active and inactive states is governed by a memoryless Markovian process, where the waiting times for transitions follow exponential distributions.

However, promoter-state switching involves multiple biochemical reaction processes, resulting in the number of effective states of most promoters being greater than two and diverse switching between states. Hence, we consider the more general TX model, in which the dwell time of the state switch follows the general distribution. The process of gene transcription is described by following the reaction diagram

$$\begin{aligned}&R_1 : \mathrm{OFF} \xrightarrow{f_{\mathrm{off}}(t,\omega_{\mathrm{off}})} \mathrm{ON},\\ &R_2 : \mathrm{ON} \xrightarrow{f_{\mathrm{on}}(t,\omega_{\mathrm{on}})} \mathrm{OFF},\\ &R_3 : \mathrm{ON} \xrightarrow{k_{\mathrm{syn}}} \mathrm{ON} + \mathrm{mRNA},\\ &R_4 : \mathrm{mRNA} \xrightarrow{k_{\mathrm{deg}}} \varnothing,\end{aligned} \tag{2}$$

where $R_1$ and $R_2$ describe the gene state switches randomly between the ON state and OFF state. The dwell time distributions of OFF and ON state are $f_{\mathrm{off}}(t,\omega_{\mathrm{off}})$ and $f_{\mathrm{on}}(t,\omega_{\mathrm{on}})$, respectively. When the gene is in the active state, $R_3$, the synthesis of mRNA molecules is governed by a Poisson process, with a probability per unit time of a synthesis event equal to $k_{\mathrm{syn}}$. Each mRNA molecule is stochastically degraded, $R_4$, with probabilities per unit time, per molecule, of $k_{\mathrm{deg}}$. Each TX model can be determined by a group of parameters, denoted as $\boldsymbol{\theta}_i = \left(\boldsymbol{\omega}_{\mathrm{off}}, \boldsymbol{\omega}_{\mathrm{on}}, k_{\mathrm{syn}}, k_{\mathrm{deg}}\right)$.

#### 1.2 Model analysis

In this part, we theoretically solve the statistics of the formula. Let $M(t)$ represent the number of mRNA at time $t$, and $G(t)$ be the state at time $t$. $E_{\text{off}}(t)$ ($E_{\text{on}}(t)$) represents the elapsed time for the gene state to transition to the OFF (ON) state at time $t$. Then, $\{M(t), G(t), E_{\text{off}}(t), E_{\text{on}}(t); t \geq 0\}$ is a continuous Markov process. Let $p_{\text{off}}(n,\tau,t)$ and $p_{\text{on}}(n,\tau,t)$ represent $n$ mRNA molecules produced at time $t$ and the elapsed time is the probability density function of $\tau$. Therefore, we can get

$$\begin{aligned} p_{\text{off}}(n,\tau,t)\Delta\tau &= \Pr\{N(t)=n, G(t)=\text{OFF}, \tau < E_{\text{off}}(t) \leq \tau+\Delta\tau\}, \\ p_{\text{on}}(n,\tau,t)\Delta\tau &= \Pr\{N(t)=n, G(t)=\text{ON}, \tau < E_{\text{on}}(t) \leq \tau+\Delta\tau\}. \end{aligned} \tag{3}$$

From the above equation, we can get the following Chapman–Kolmogorov backward equation

$$\begin{aligned} p_{\text{off}}(n,\tau+\Delta t,t+\Delta t) &= p_{\text{off}}(n,\tau,t)(1-nr_{\text{deg}}\Delta t)(1-H_{\text{off}}(\tau)\Delta t) \\ &\quad + p_{\text{off}}(n+1,\tau,t)(n+1)r_{\text{deg}}\Delta t(1-H_{\text{off}}(\tau)\Delta t) + o(\Delta t), \\ p_{\text{on}}(n,\tau+\Delta t,t+\Delta t) &= p_{\text{on}}(n,\tau,t)(1-nr_{\text{deg}}\Delta t)(1-r_{\text{syn}}\Delta t)(1-H_{\text{on}}(\tau)\Delta t) \\ &\quad + p_{\text{on}}(n+1,\tau,t)(n+1)r_{\text{deg}}\Delta t(1-r_{\text{syn}}\Delta t)(1-H_{\text{on}}(\tau)\Delta t) \\ &\quad + p_{\text{on}}(n-1,\tau,t)(1-nr_{\text{deg}}\Delta t)r_{\text{syn}}\Delta t(1-H_{\text{on}}(\tau)\Delta t) + o(\Delta t), \end{aligned} \tag{4}$$

where $H_{\text{off}}(\tau) = f_{\text{off}}(\tau)/S_{\text{off}}(\tau)$ is a hazard rate function with the survival function $S_{\text{off}}(\tau) = \int_{\tau}^{\infty} f_{\text{off}}(t)dt$. The definitions of $H_{\text{on}}(\tau)$ and $S_{\text{on}}(\tau)$ are similar.

Next, we focus on the steady-state distributions. Simulations in the main text have verified that the stationary distributions of $p_{\text{off}}(n,\tau,t)$ and $p_{\text{on}}(n,\tau,t)$ exist, and they are denoted by $p_{\text{off}}(n,\tau)$ and $p_{\text{on}}(n,\tau)$, respectively. Eq. (4) converts to the following stationary chemical master equation in the limit of small $\Delta t$ and large $t$,

$$\begin{aligned} \frac{\partial}{\partial\tau} p_{\text{off}}(n,\tau) &= (n+1)k_{\text{deg}} p_{\text{off}}(n+1,\tau) - (nk_{\text{deg}} + H_{\text{off}}(\tau))p_{\text{off}}(n,\tau), \\ \frac{\partial}{\partial\tau} p_{\text{on}}(n,\tau) &= (n+1)k_{\text{deg}} p_{\text{on}}(n+1,\tau) + k_{\text{syn}} p_{\text{on}}(n-1,\tau) - (nk_{\text{deg}} + k_{\text{syn}} + H_{\text{on}}(\tau))p_{\text{on}}(n,\tau), \end{aligned} \tag{5}$$

with the integral boundary condition

$$p_{\text{off}}(n,0) = \int_0^{\infty} p_{\text{on}}(n,\tau)H_{\text{on}}(\tau)d\tau, \quad p_{\text{on}}(n,0) = \int_0^{\infty} p_{\text{off}}(n,\tau)H_{\text{off}}(\tau)d\tau. \tag{6}$$

Based on Eq. (5) with its boundary conditions, we use the binomial moment method to calculate the mRNA stationary distribution $P(n) = \int_0^{\infty}[p_{\text{off}}(n,\tau) + p_{\text{on}}(n,\tau)]d\tau$ and its statistical characteristics. Binomial moments of the mRNA stationary distribution are defined as $b_n = \sum_{m \geq n}^{\infty} \binom{m}{n} P(m)$, where the symbol $\binom{m}{n}$ represents the combinatorial number. Note that binomial moments converge to zero as their orders go to infinity, and it can be used to reconstruct $P(n)$ by $P(n) = \sum_{m=n}^{\infty} (-1)^{m-n} \binom{m}{n} b_m$. After some algebra calculations of Eq. (4), we can obtain the $n$-th binomial moment of mRNA in a recursive form

$$b_n = \frac{1}{n!}\left(\frac{k_{\text{syn}}}{k_{\deg}}\right)^n \sum_{i=0}^{n-1}\binom{n-1}{i} C_i \sum_{j=0}^{n-1-i}\binom{n-1-i}{j}(-1)^{n-1-i-j}\tilde{S}_{\text{on}}\left((n-1-j)k_{\deg}\right), \tag{7}$$

where

$$C_n = \frac{\tilde{f}_{\text{off}}\left(nk_{\deg}\right)}{1-\tilde{f}_{\text{off}}\left(nk_{\deg}\right)\tilde{f}_{\text{on}}\left(nk_{\deg}\right)} \sum_{i=0}^{n-1}\binom{n}{i} C_i \sum_{j=0}^{n-i}\binom{n-i}{j}(-1)^{n-i-j}\tilde{f}_{\text{on}}\left((n-j)k_{\deg}\right), \tag{8}$$

for $n = 1,2,\cdots$. Here $C_0 = \left(\langle\tau_{\text{off}}\rangle + \langle\tau_{\text{on}}\rangle\right)^{-1}$ is equal to the burst frequency. $\tilde{f}(s)$ represents the Laplace transform of function $f(t)$. Especially, $\tilde{S}_{\text{on}}(s) = \left(1-\tilde{f}_{\text{on}}(s)\right)/s$ and $\tilde{S}_{\text{on}}(0) = \langle\tau_{\text{on}}\rangle$. According to the relationship between binomial moments and central moments and Eq. (7) and Eq. (8), we obtain the mean and noise of mRNA expression

$$\begin{aligned} &\text{Mean}=\frac{k_{\text{syn}}\langle\tau_{\text{on}}\rangle}{k_{\deg}\left(\langle\tau_{\text{off}}\rangle+\langle\tau_{\text{on}}\rangle\right)}, \\ &\text{CV}^2 = \frac{1}{\text{Mean}} + \frac{\langle\tau_{\text{off}}\rangle}{\langle\tau_{\text{on}}\rangle} - \frac{\langle\tau_{\text{off}}\rangle+\langle\tau_{\text{on}}\rangle}{k_{\deg}\langle\tau_{\text{on}}\rangle^2}\frac{\left(1-\tilde{f}_{\text{off}}\left(k_{\deg}\right)\right)\left(1-\tilde{f}_{\text{on}}\left(k_{\deg}\right)\right)}{1-\tilde{f}_{\text{off}}\left(k_{\deg}\right)\tilde{f}_{\text{on}}\left(k_{\deg}\right)}. \end{aligned} \tag{9}$$

Note that if let $\tau_{\text{off}}k_{\deg}$ and $\tau_{\text{on}}k_{\deg}$ be the rescaled random variables for OFF-state and ON-state dwell times, and $k_{\text{syn}}/k_{\deg}$ be the rescaled mean synthesis rate, the mean expression not only equals the product of the mean synthesis rate and the stationary probability of ON state, but also the product of burst size and burst frequency.

The binomial moment can obtain any order statistics of the model solution. Next, we introduce the five statistics we use in training neural networks: (1) The mean value $\mu_1$ is the most commonly used indicator in statistics, and it represents the average level of mRNA expression in scRNA-seq data. (2) The noise strength is a measurement of the dispersion of the probability distribution, defined as $\mu_2/\mu_1^2$, where $\mu_2$ is the variance. (3) The Fano factor is another statistic that measures the dispersion of a probability distribution relative to a Poisson distribution, defined as $\mu_2/\mu_1$. (4) The skewness is a description of the symmetry of the distribution, and it is defined as $\mu_3/\mu_2^{3/2}$, where $\mu_3$ is the third central moment. (5) The kurtosis describes whether the peak of the distribution is abrupt or flat, which is defined as $\mu_4/\mu_2^2$, where $\mu_4$ is the fourth central moment. (6) The bimodality coefficient can describe the bimodal distribution, which is usually a critical feature in a dynamical system. Precisely, we can calculate the central moments with the binomial moment:

$$\mu_k(t) = \left(-b_1(t)\right)^k + \sum_{i=0}^{k-1}\sum_{j=0}^{k-1} R(k,i,j)(j!)\left(b_1(t)\right)^i b_j(t), \tag{10}$$

in which $R(k,i,j) = (-1)^i\binom{k}{i}s(k-i,j)$ with $S(n,k) = \sum_{i=0}^{k}(-1)^{k-i}\binom{k}{i}i^n$ being the Stirling number of the second kind. Therefore, the above summary statistics can be expressed by binomial moments as follows

$$
\begin{aligned}
&\text{Mean} = \mathrm{b}_1, \\
&\text{NoiseStrength} = \frac{2b_2 + b_1 - b_1^2}{b_1^2}, \\
&\text{FanoFactor} = \frac{2b_2 + b_1 - b_1^2}{b_1}, \\
&\text{Skewness} = \frac{6b_3 + 6b_2 + b_1 - 3b_1\left(2b_2 + b_1\right) + 2b_1^3}{\left(2b_2 + b_1 - b_1^2\right)^{3/2}}, \\
&\text{Kurtosis} = \frac{24b_4 + 36b_3 + 14b_2 + b_1 - 4b_1\left(6b_3 + 6b_2 + b_1\right) + 6b_1^2\left(2b_2 + b_1\right) - 3b_1^4}{\left(2b_2 + b_1 - b_1^2\right)^2} - 3, \\
&\text{BimodalityCoefficient} = \frac{\text{Skewness}^2 + 1}{\text{Kurtosis}}.
\end{aligned}
\tag{11}
$$

It should be noted that we can extend the summary statistics to higher-order moments because our binomial moments can compute arbitrary high-order moment statistics.

#### 1.3 Algorithm pipeline of deepTX framework

**Dataset:** We use the Sobol algorithm to sample from a preset range of TXmodel parameters to obtain the parameters $\{\boldsymbol{\theta}_i\}$. For each set of $\boldsymbol{\theta}_i$, the distribution and moments are obtained using the SSA algorithm and binomial moment theory to form the dataset $\{(\boldsymbol{\theta}_i, P_{\text{simulation},i}, \boldsymbol{s}_i)\}$.

**Input**：A batch parameter of TXmodel $\{\boldsymbol{\theta}_i\}_{\text{batch}}$.

**Output:** The solution of the TXmodel obtained from the neural network and the corresponding statistics $\{(P_{\text{neuralnet},i}, \hat{\boldsymbol{s}}_i)\}_{\text{batch}}$.

Initialize the neural network architecture as well as the weights and select hyperparameters.

**For every epoch do**

(1) Sampling from dataset $\{(\boldsymbol{\theta}_i, P_{\text{simulation},i}, \boldsymbol{s}_i)\}$ to get a batch with parameter set $\{(\boldsymbol{\theta}_i, P_{\text{simulation},i}, \boldsymbol{s}_i)\}_{\text{batch}}$.

(2) Input parameter set $\{\boldsymbol{\theta}_i\}_{\text{batch}}$ to the neural network to get the distribution and statistics predicted by the neural network $\{(P_{\text{neuralnet},i}, \hat{\boldsymbol{s}}_i)\}_{\text{batch}}$.

(3) Train the neural network by the loss function $L = \sum_{i=1}^{n_{\text{batch}}} \mathrm{KL}\left(P_{\text{simulation},i}, P_{\text{neuralnet},i}\right) + \lambda \sum_{j=1}^{n_{\text{STATS}}} \log\left(s_{ij} / \hat{s}_{ij}\right)$.

**End for**

#### 1.4 Derivation of hierarchical negative binomial mixture distribution

Assume the following hierarchical distribution

$$
\begin{aligned}
&Y|X \sim \mathrm{Binomial}\left(x|y, p\right), \\
&X \sim \mathrm{NB}\left(n, q\right).
\end{aligned}
\tag{12}
$$

Calculate the marginal distribution as follows

$$
\begin{aligned}
\Pr[Y=y] &= \sum_{x=0}^{\infty} P(X=x, Y=y) = \sum_{x=0}^{\infty} P(Y=y \mid X=x) P(X=x) \\
&= \sum_{x=y}^{\infty} \binom{x}{y} \binom{x+n-1}{x} p^y (1-p)^{x-y} (1-q)^n q^x \ (\text{conditional probability is 0 if } y > x) \\
&= \sum_{x=y}^{\infty} \frac{x!}{y!(x-y)!} \frac{(x+n-1)!}{x!(n-1)!} \frac{(n+y-1)!}{(n+y-1)!} p^y (1-p)^{x-y} (1-q)^n q^x \\
&= \sum_{x=y}^{\infty} \binom{x+n-1}{x-y} \binom{n+y-1}{y} p^y (1-p)^{x-y} (1-q)^n q^x q^y q^{-y} \\
&= \binom{n+y-1}{y} (pq)^y (1-q)^n \sum_{x=y}^{\infty} \binom{x+n-1}{x-y} \left((1-p)q\right)^{x-y} \\
&= \binom{n+y-1}{y} \left(\frac{pq}{1-q+pq}\right)^y \left(\frac{1-q}{1-q+pq}\right)^n .
\end{aligned} \tag{13}
$$

Further,

$$
Y \sim \mathrm{NB}\left(n, \frac{pq}{1-q+pq}\right). \tag{14}
$$

If we assume that

$$
P(X=x) = w_1 P_1(X=x) + w_2 P_2(X=x), \tag{15}
$$

where $P_1$ and $P_2$ are both negative binomial distributions, then we can calculate them separately, and the result is the addition of two negative binomial distributions.

#### 2. Relationship between noise and burst dynamics

For a TX model,

$$
\begin{aligned}
&\mathrm{BF} = \frac{1}{\langle \tau_{\mathrm{on}} \rangle + \langle \tau_{\mathrm{off}} \rangle}, \quad \mathrm{BS} = r_{\mathrm{syn}} \langle \tau_{\mathrm{on}} \rangle, \\
&\langle X \rangle = \frac{r_{\mathrm{syn}} \langle \tau_{\mathrm{on}} \rangle}{r_{\deg} \left( \langle \tau_{\mathrm{on}} \rangle + \langle \tau_{\mathrm{off}} \rangle \right)} = \frac{\mathrm{BF} \cdot \mathrm{BS}}{r_{\deg}}, \\
&\eta_X^2 = \frac{1}{\langle X \rangle} + \frac{r_{\deg} \langle \tau_{\mathrm{off}} \rangle^2}{r_{\deg} \langle \tau_{\mathrm{on}} \rangle \langle \tau_{\mathrm{off}} \rangle + \langle \tau_{\mathrm{on}} \rangle + \langle \tau_{\mathrm{off}} \rangle}.
\end{aligned} \tag{16}
$$

When the mean protein $\langle X \rangle$ is fixed to a constant $M$, $\mathrm{BS}$ and $\mathrm{BF}$ are inversely proportional. We consider following cases to indicate that the noise $\eta_X^2$ is a monotonically decreasing (increasing) function about $\mathrm{BF}$ ($\mathrm{BS}$).

(1) $\langle \tau_{\mathrm{on}} \rangle$ is fixed.

$\langle \tau_{\mathrm{off}} \rangle$ and $r_{\mathrm{syn}}$ satisfy the following relation,

$$
r_{\mathrm{syn}} = M r_{\deg} \left( 1 + \frac{\langle \tau_{\mathrm{off}} \rangle}{\langle \tau_{\mathrm{on}} \rangle} \right). \tag{17}
$$

On the other hand, $\mathrm{BF}$ decreases with an increasing $\langle\tau_{\mathrm{off}}\rangle$, and $\mathrm{BS}$ increases with the corresponding increasing $r_{\mathrm{syn}}$. In this case,

$$\eta_X^2=\frac{1}{M}+\frac{r_{\mathrm{deg}}}{\frac{r_{\mathrm{deg}}\langle\tau_{\mathrm{on}}\rangle+1}{\langle\tau_{\mathrm{off}}\rangle}+\frac{\langle\tau_{\mathrm{on}}\rangle}{\langle\tau_{\mathrm{off}}\rangle^2}} \tag{18}$$

increases with an increasing $\langle\tau_{\mathrm{off}}\rangle$. Therefore, the noise $\eta_X^2$ is a monotonically decreasing (increasing) function about $\mathrm{BF}$ ($\mathrm{BS}$).

(2) $r_{\mathrm{syn}}$ is fixed.

$\langle\tau_{\mathrm{off}}\rangle$ and $\langle\tau_{\mathrm{on}}\rangle$ satisfy the following relation,

$$\langle\tau_{\mathrm{off}}\rangle=K\langle\tau_{\mathrm{on}}\rangle, \tag{19}$$

where the constant $K=(1-C)/C$, and $C=Mr_{\mathrm{deg}}/r_{\mathrm{syn}}$. Note that $C\in(0,1)$, then $K>0$. Thus, $\langle\tau_{\mathrm{off}}\rangle$ is directly proportional to $\langle\tau_{\mathrm{on}}\rangle$.

On the other hand, $\mathrm{BF}$ decreases with an increasing $\langle\tau_{\mathrm{on}}\rangle$ and the corresponding increasing $\langle\tau_{\mathrm{off}}\rangle$, and $\mathrm{BS}$ increases with an increasing $\langle\tau_{\mathrm{on}}\rangle$. In this case,

$$\eta_X^2=\frac{1}{M}+\frac{r_{\mathrm{deg}}}{\frac{r_{\mathrm{deg}}}{K}+\frac{1}{K\langle\tau_{\mathrm{on}}\rangle}+\frac{1}{K^2\langle\tau_{\mathrm{on}}\rangle}} \tag{20}$$

increases with an increasing $\langle\tau_{\mathrm{off}}\rangle$. Therefore, the noise $\eta_X^2$ is a monotonically decreasing (increasing) function about $\mathrm{BF}$ ($\mathrm{BS}$).

(3) The growth of $r_{\mathrm{syn}}$ is consistent with the growth of $\langle\tau_{\mathrm{on}}\rangle$.

$\langle\tau_{\mathrm{off}}\rangle$ can be obtained by

$$\langle\tau_{\mathrm{off}}\rangle=K(r_{\mathrm{syn}})\langle\tau_{\mathrm{on}}\rangle, \tag{21}$$

where $K(r_{\mathrm{syn}})=(r_{\mathrm{syn}}-Mr_{\mathrm{deg}})/Mr_{\mathrm{deg}}$ is a monotonically increasing function about $r_{\mathrm{syn}}$. Similar to case 2, it is easy to derive that the noise $\eta_X^2$ is a monotonically decreasing (increasing) function about $\mathrm{BF}$ ($\mathrm{BS}$).

#### 3. The definition of Quality Score

The Quality Score (QS) is calculated based on the difference in z-scores derived from GSVA (Gene set variation analysis) of gene sets upregulated and downregulated during the quiescent phase, and is defined as $\mathrm{QS}=z(\text{up genes})-z(\text{down genes})$ (Wiecek AJ et al., 2023). $z(\text{up genes})$ represents the standardized enrichment score of the gene set upregulated during quiescence in each sample. A higher value indicates that the quiescence-associated upregulated genes are actively expressed, suggesting that the sample is more likely to be in a quiescent (G0) state. $z(\text{down genes})$ corresponds to the standardized enrichment score of genes downregulated during quiescence. A lower value implies effective suppression of these genes, which is also consistent with quiescence. The difference score $\mathrm{QS}$ serves as an integrated indicator of the quiescent state:

A higher value reflects simultaneous activation of quiescence-associated upregulated genes and repression of downregulated genes, indicating a gene expression profile that strongly aligns with the G0/quiescent state. A lower or negative value suggests a deviation from the quiescent signature, potentially reflecting a proliferative state or failure to enter quiescence.

#### 4. The rationale for the parameter space

In our study, we considered five parameters: $\boldsymbol{\theta} = (k_{\text{off}}, r_{\text{off}}, k_{\text{on}}, r_{\text{on}}, k_{\text{syn}})$. The parameters $k_{\text{off}}$ and $k_{\text{on}}$ represent the number of intermediate reaction steps involved in transcriptional state transitions. These values were sampled uniformly from the range 1 to 15, which aligns with biological evidence indicating that most genes undergo either direct (single-step) transitions or a small number of intermediate steps, typically fewer than ten (Tunnacliffe E & Chubb JR, 2020). This range is sufficient to capture both widely used single-step models and more detailed multi-step mechanisms without introducing biologically implausible complexity.

Among these parameters, $r_{\text{off}}$ and $r_{\text{on}}$ denote the rate constants governing stochastic transitions between the OFF and ON transcriptional states, respectively. The mean duration of the OFF state, which corresponds to the time between transcriptional bursts, is given by $\langle \tau_{\text{off}} \rangle = k_{\text{off}} / r_{\text{off}}$, and falls within the range $\langle \tau_{\text{off}} \rangle \in (0.1, 150)$. Experimental measurements report a median value of $\langle \tau_{\text{off}} \rangle$ approximately 3.7 (Gupta A et al., 2022) which is well contained within this range. Similarly, the mean duration of the ON state, referred to as the burst duration, is defined by $\langle \tau_{\text{on}} \rangle = k_{\text{on}} / r_{\text{on}}$, and spans the interval $\langle \tau_{\text{on}} \rangle \in (0.1, 1500)$. The experimentally observed median value of 0.12 (Gupta A et al., 2022) confirms that the parameter range adequately captures biologically realistic dynamics.

The parameter $k_{\text{syn}}$ represents the normalized synthesis rate after accounting for molecular degradation. Its range was chosen based on empirical observations of transcriptional burst sizes, which typically vary from single molecules to several dozen (Gupta A et al., 2022). Considering the relationship $\text{BS} = k_{\text{syn}} * \langle \tau_{\text{on}} \rangle$, the selected range of $k_{\text{syn}}$ ensures that the experimentally observed burst sizes are well represented within the defined parameter space.

#### Supplementary Figures

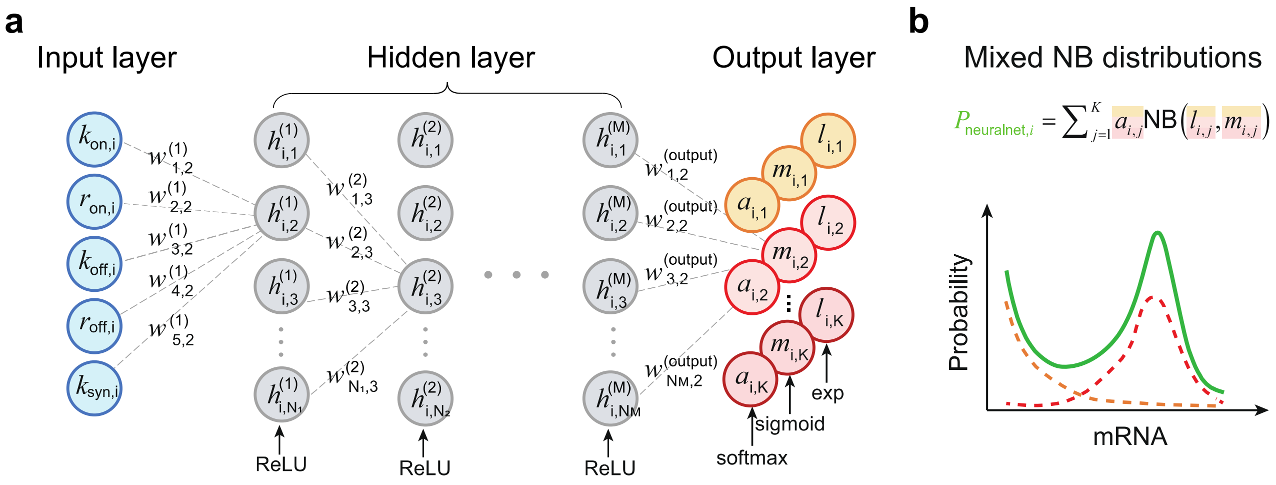

**Supplementary Figure S1. Network architecture of DeepTX. (a)** The input of the neural network is the parameter of the mechanism model, and the output is the parameter of the mixed negative binomial distribution. The neural network has one or more hidden layers. The layers are connected through the weight parameter $\boldsymbol{w}$. Specifically, the activation functions corresponding to the hidden layer and the output layer are marked at the bottom of the layer. **(b)** Schematic diagram of mixed negative binomial distribution.

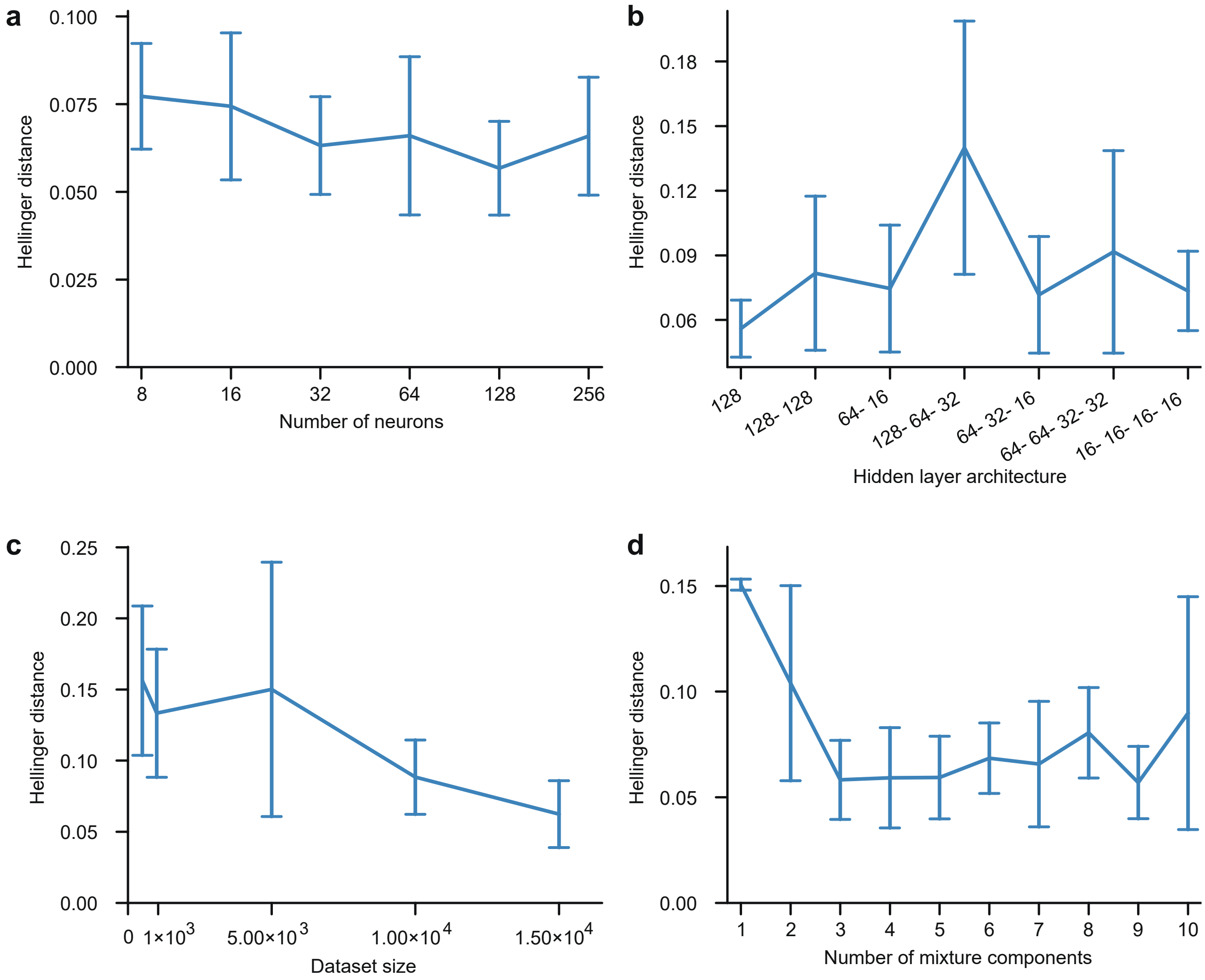

**Supplementary Figure S2. Hyperparameter tuning for DeepTX. (a)** Hyperparameter tuning of neuron number. **(b)** Hyperparameter tuning of neuron hidden layer architecture. **(c)** Hyperparameter tuning of the number of dataset size. **(d)** Hyperparameter tuning of the mixture components.

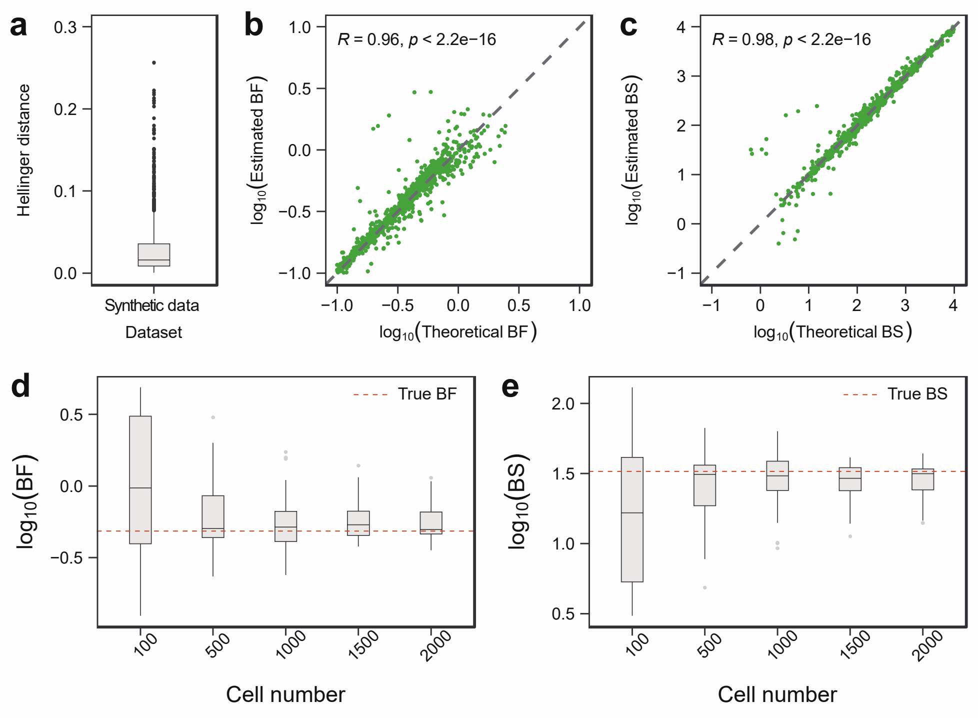

**Supplementary Figure S3. Precision and robustness of inferred results. (a)** Verification of the Hellinger distance between the distribution obtained by DeepTXsolver and the distribution obtained by SSA simulation of the test set. **(b-c)** Correlation scatterplot of inferred and theoretical BS (BF). (**d-e)** Boxplots of BF and BS obtained by inferring from different cell numbers, where the red dashed line corresponds to the true parameter.

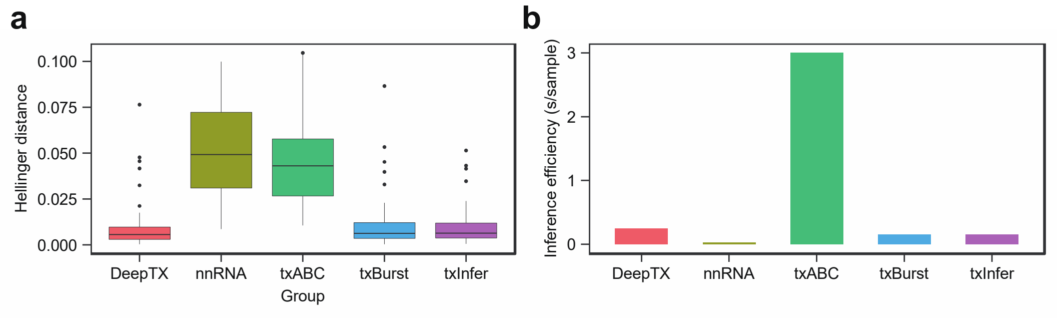

**Supplementary Figure S4.** Comparison of inference accuracy and efficiency between DeepTX and other algorithms. **(a)** Hellinger distance for accuracy. **(b)** Average inference time per sample (seconds) for efficiency.

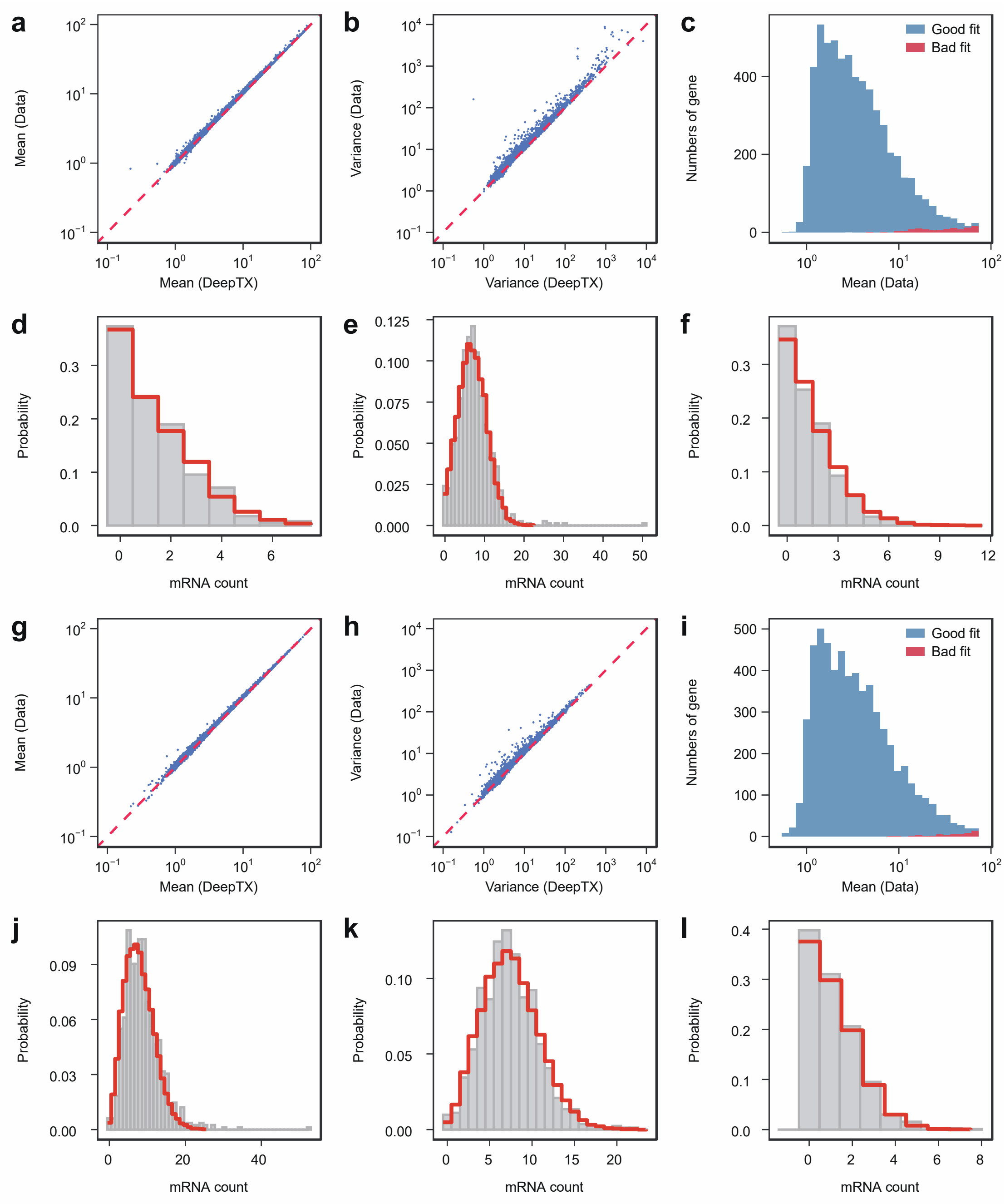

**Supplementary Figure S5. Inference results of scRNA-seq data treated with DMSO and IdU. (a-b)** Comparative analysis of the mean and variance between the inferred distribution and the distribution of control scRNA-seq data. **(c)** The goodness-of-fit test is demonstrated for all inferred genes, with the histogram displaying gene counts exhibiting good fit (blue) and poor fit (red) relative to mean expression levels. **(d-f)** Comparison between mRNA distributions and DeepTX-inferred distributions for scRNA-seq data of three genes. The gray bars and the red stepped lines correspond to the distributions derived from scRNA-seq data and inferred by DeepTX, respectively. **(g-l)** The result of IdU treatment scRNA-seq data, similar to **(a-f)**.

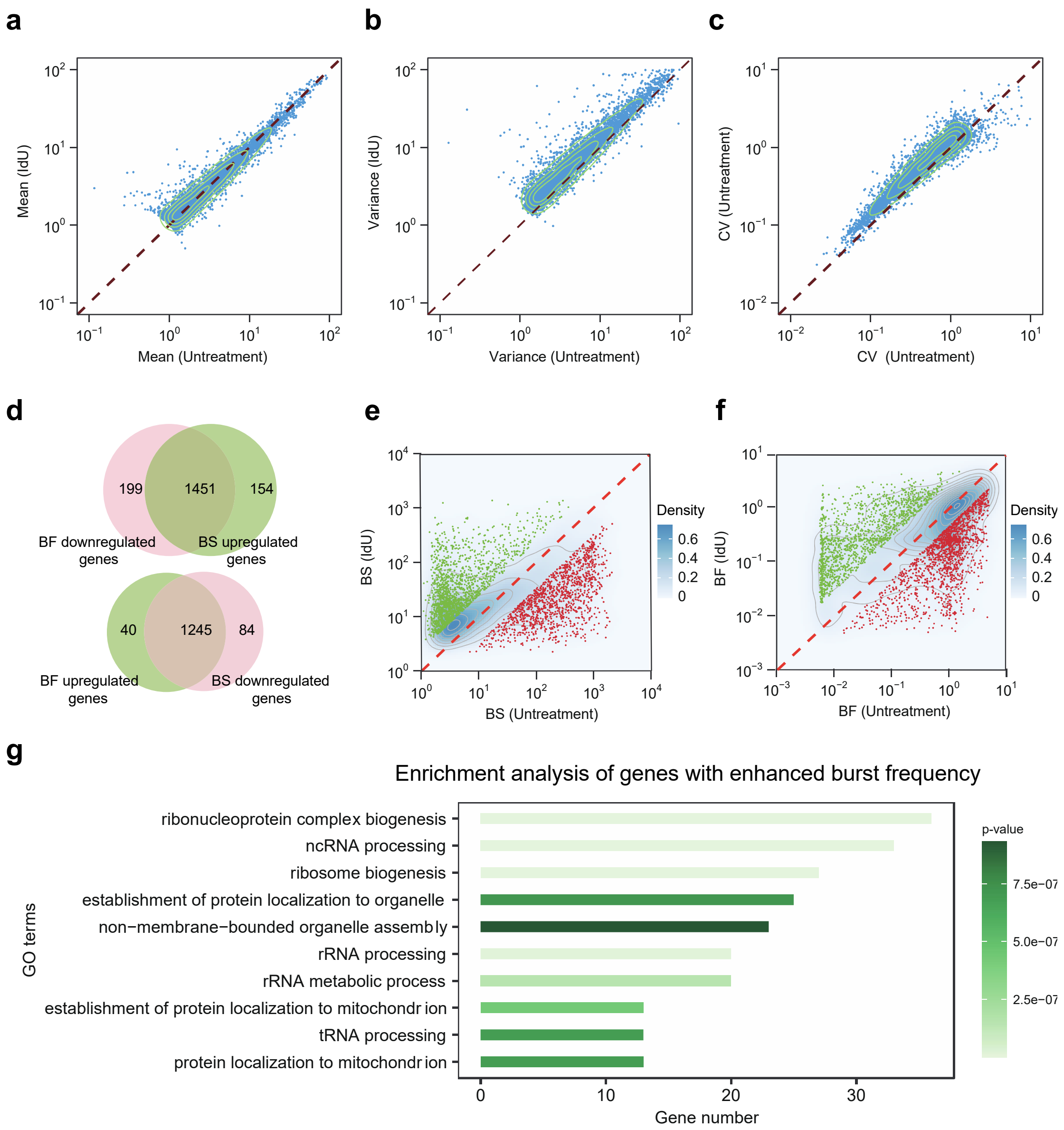

**Supplementary Figure S6. Analysis results of scRNA-seq data treated with IdU and without IdU.(a-c)** Mean, variance and CV analysis results of IdU treatment and control scRNA-seq data, the blue scatter represents genes. **(d)** The number of intersections of genes up-regulated by BF(BS) and genes down-regulated by BF (BS). **(e)** Red dots represent genes significantly down-regulated by BS, and green dots represent genes significantly up-regulated by BS. **(f)** Red dots represent genes significantly down-regulated by BF, and green dots represent genes significantly up-regulated by BF. **(g)** Enrichment analysis results of BS down-regulated differential genes.

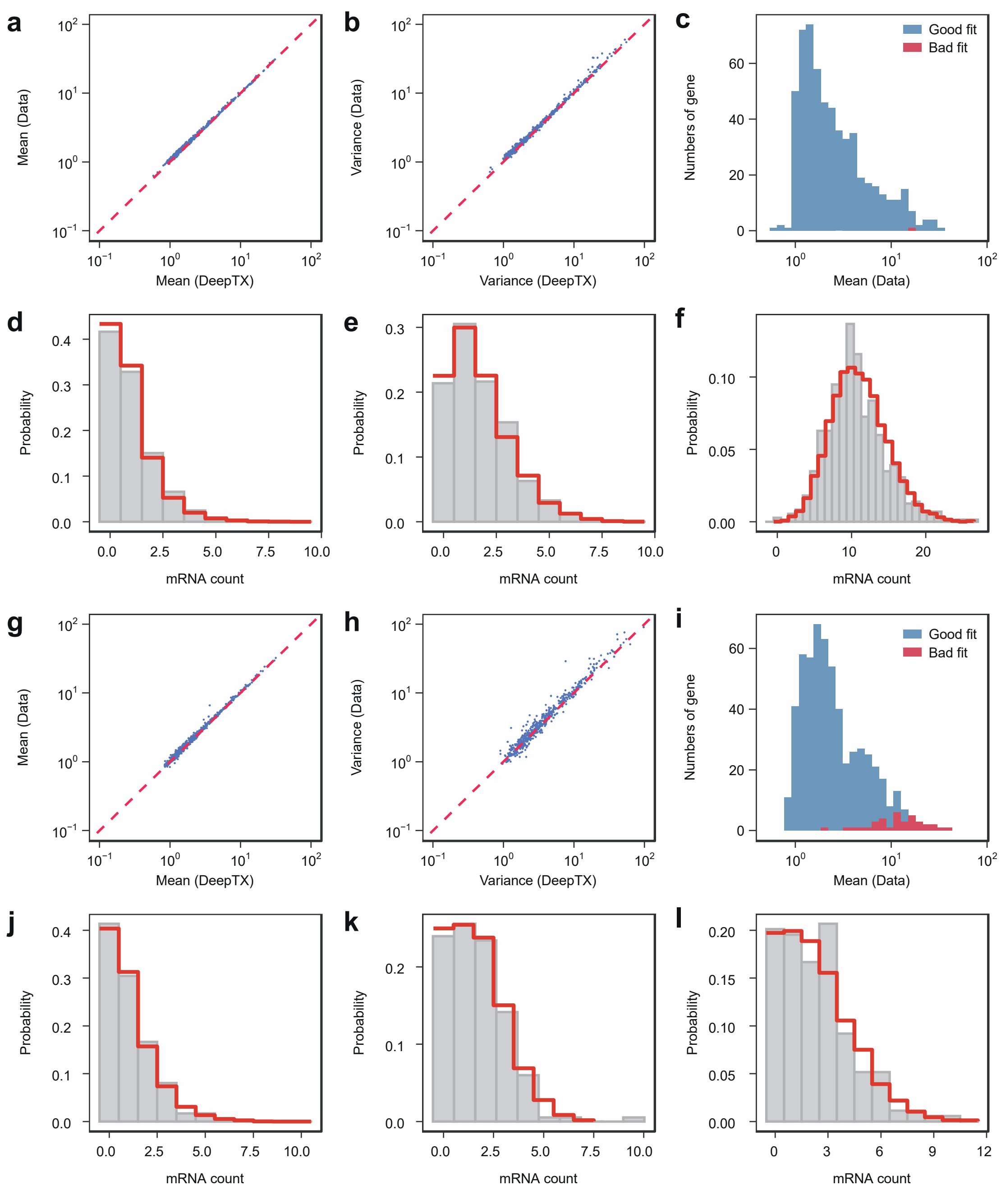

**Supplementary Figure S7. Inference results of scRNA-seq data treated with low dose 5FU and without 5FU treatment. (a-b)** Comparative analysis of the mean and variance between the inferred distribution and the distribution of control scRNA-seq data. **(c)** The goodness-of-fit test is demonstrated for all inferred genes, with the histogram displaying gene counts exhibiting good fit (blue) and poor fit (red) relative to mean expression levels. **(d-f)** Comparison between mRNA distribution and inferred distribution by DeepTX for scRNA-seq data of three genes. The gray bars and the red stepped line correspond to the distributions derived from scRNA-seq data and inferred by DeepTX, respectively. **(g-l)** The result of low dose 5FU treatment scRNA-seq data, similar to **(a-f)**.

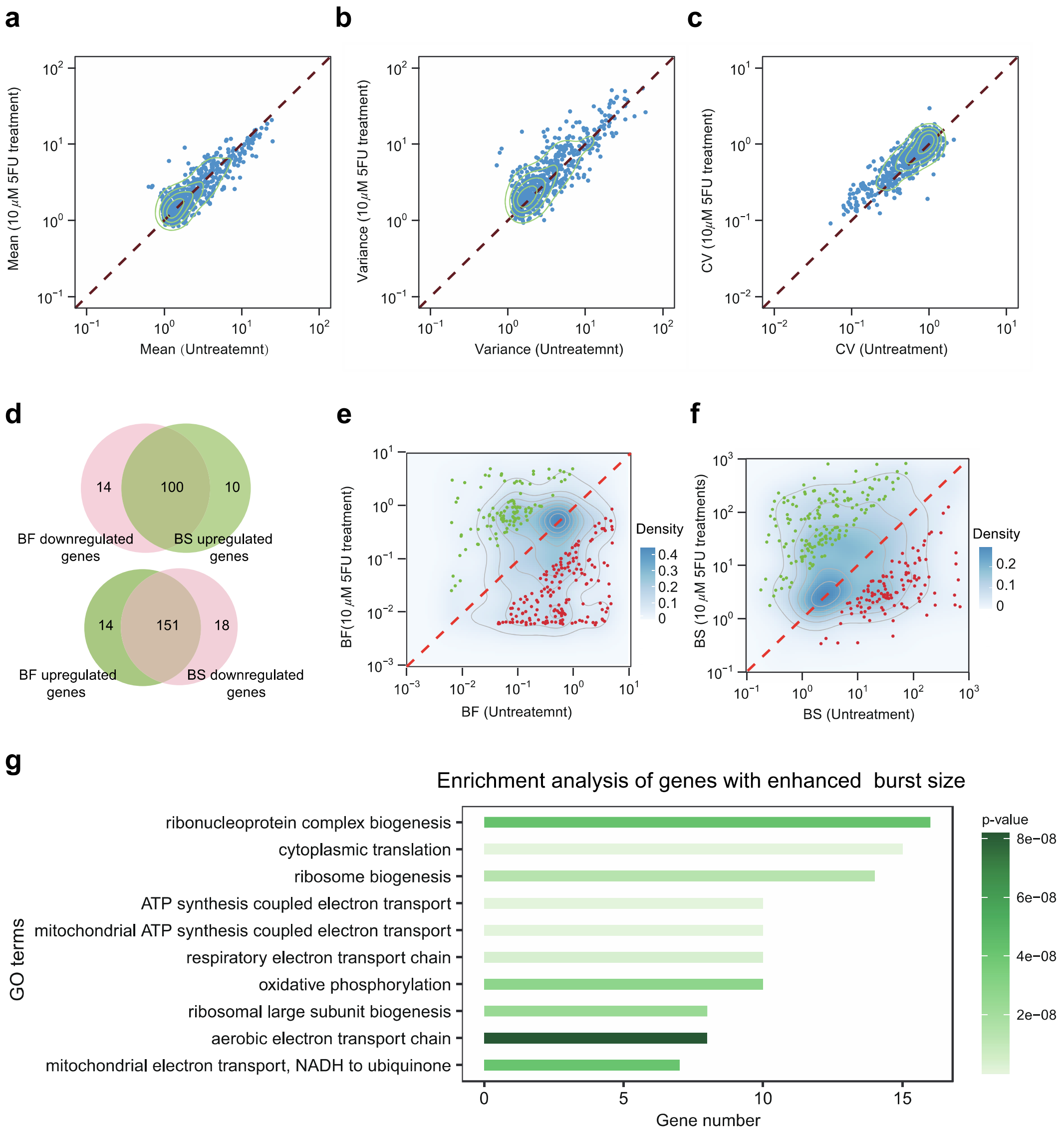

**Supplementary Figure S8. Analysis results of scRNA-seq data treated with low dose 5FU and without 5FU treatment. (a-c)** Mean, variance and CV analysis results of low dose 5FU treatment and control scRNA-seq data, each blue scatter represents a gene. **(d)** The number of intersections of genes up-regulated by BF (BS) and genes down-regulated by BF (BS). **(e)** Red dots represent genes significantly down-regulated by BF, and green dots represent genes significantly up-regulated by BF. **(f)** Red dots represent genes significantly down-regulated by BS, and green dots represent genes significantly up-regulated by BS. **(g)** Enrichment analysis results of BS up-regulated differential genes.

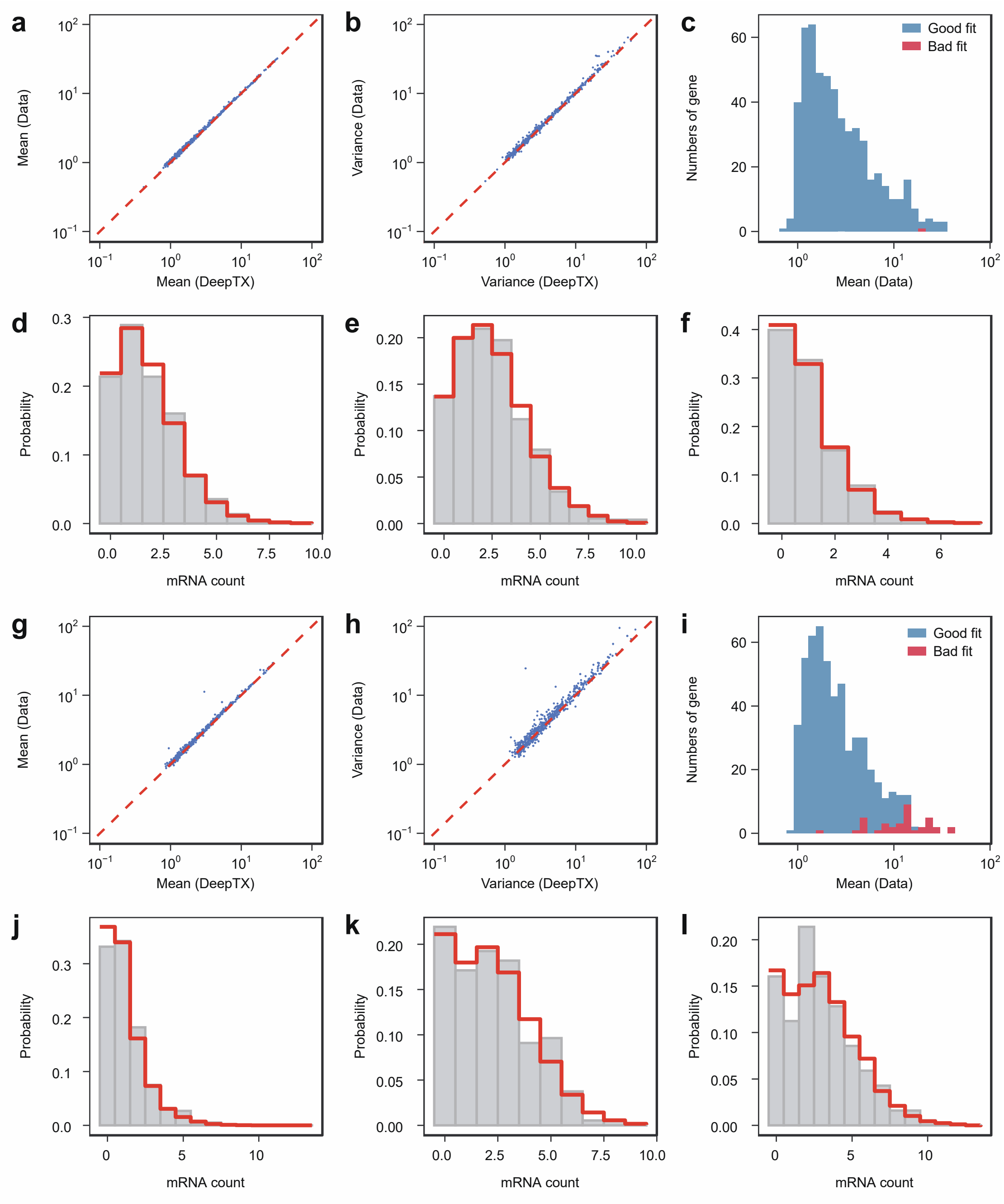

**Supplementary Figure S9. Inference results of scRNA-seq data treated with high dose 5FU and without 5FU. (a-b)** Comparative analysis of the mean and variance between the inferred distribution and the distribution of control scRNA-seq data. **(c)** The goodness-of-fit test is demonstrated for all inferred genes, with the histogram displaying gene counts exhibiting good fit (blue) and poor fit (red) relative to mean expression levels. **(d-f)** Comparison between mRNA distributions and DeepTX-inferred distributions for scRNA-seq data of three genes. The gray bars and the red stepped line correspond to the distributions derived from scRNA-seq data and inferred by DeepTX, respectively. **(g-l)** The result of high dose 5FU treatment scRNA-seq data, similar to a-f.

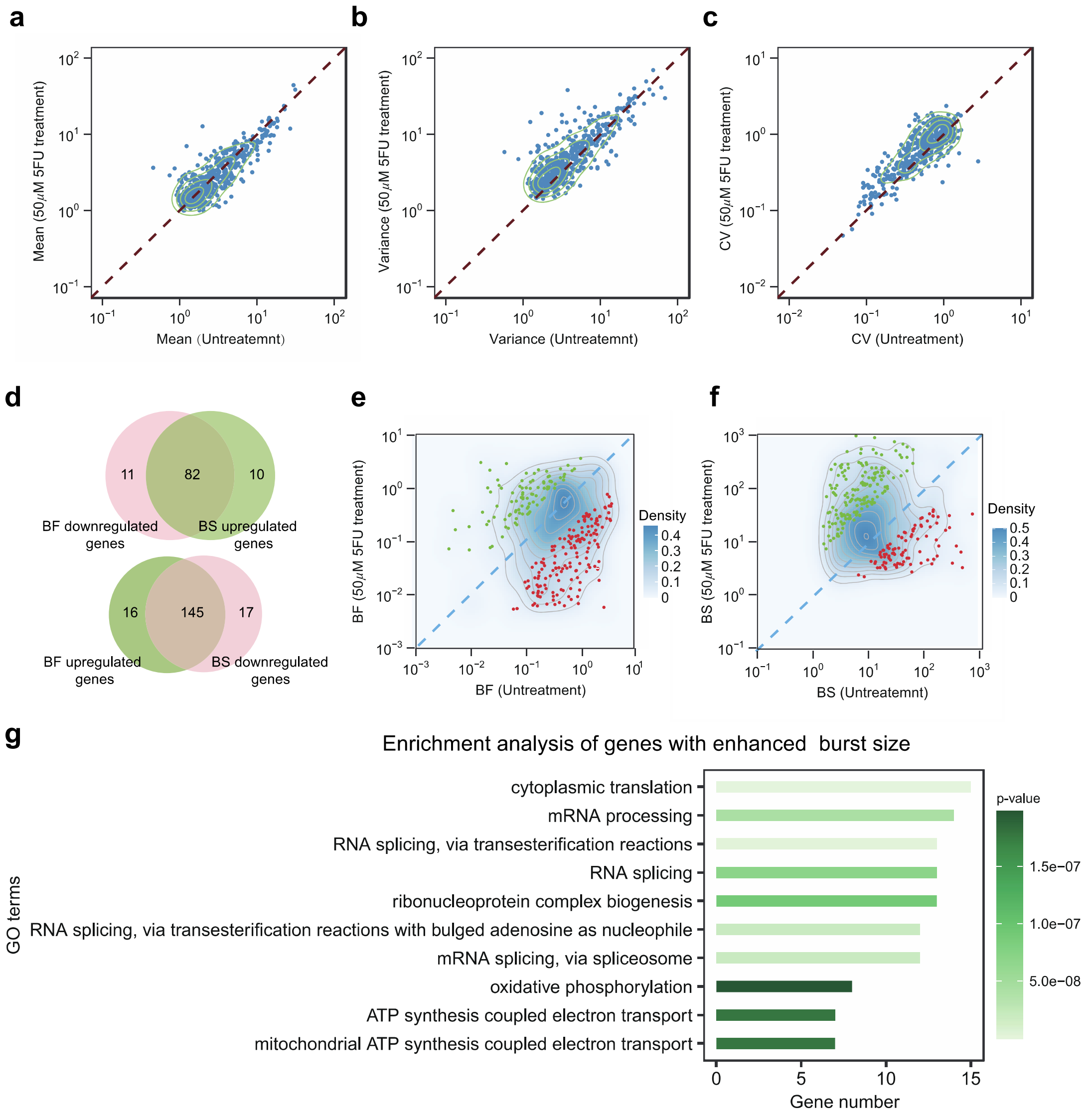

**Supplementary Figure S10. Analysis results of scRNA-seq data treated with high dose 5FU and without 5FU treatment. (a-c)** Mean, variance and CV analysis results of high dose 5FU treatment and control scRNA-seq data, each blue scatter represents a gene. **(d)** The number of intersections of genes up-regulated by BF (BS) and genes down-regulated by BF (BS). **(e)** Red dots represent genes significantly down-regulated by BF, and green dots represent genes significantly up-regulated by BF. **(f)** Red dots represent genes significantly down-regulated by BS, and green dots represent genes significantly up-regulated by BS. **(g)** Enrichment analysis results of BS up-regulated differential genes.

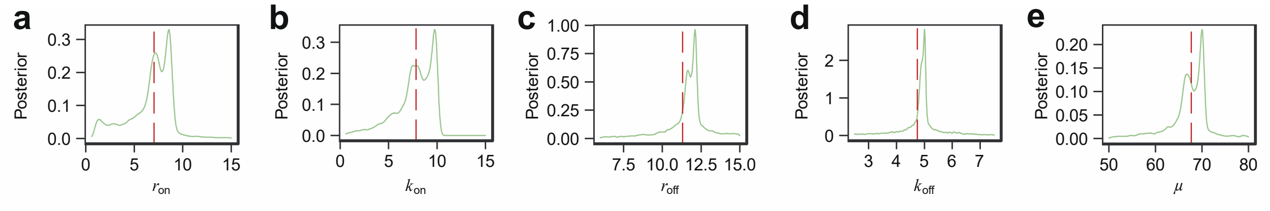

**Supplementary Figure S11.** The marginal density for the parameters of the TX model. **(a-e)** The marginal density for the five parameters of the model, where the red dashed line represents the true value.

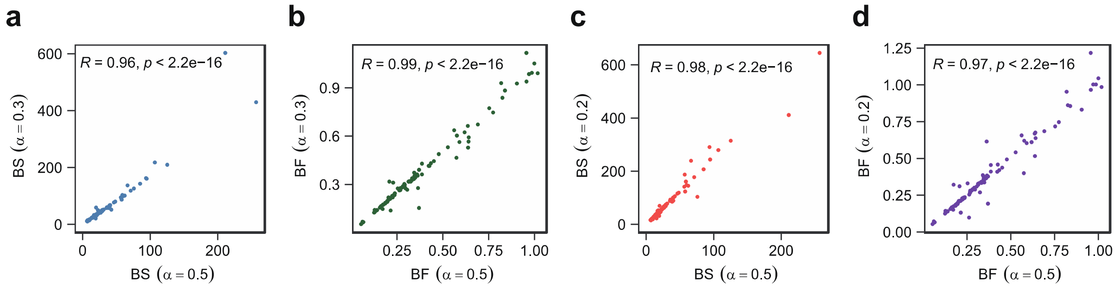

**Supplementary Figure S12.** Comparison of inferred burst dynamics under different capture efficiencies. **(a–b)** Scatter plots comparing the inferred burst size (BS) and burst frequency (BF) between capture efficiencies of 0.3 and 0.5. **(c–d)** Scatter plots comparing the inferred BS and BF between capture efficiencies of 0.2 and 0.5.

#### Reference

Wiecek, A.J., Cutty, S.J., Kornai, D., Parreno-Centeno, M., Gourmet, L.E., Tagliazucchi, G.M., Jacobson, D.H., Zhang, P., Xiong, L., Bond, G.L., Barr, A.R., Secrier, M. 2023. Genomic hallmarks and therapeutic implications of G0 cell cycle arrest in cancer. *Genome Biology* **24**: 128. DOI: https://dx.doi.org/10.1186/s13059-023-02963-4, PMID: 37221612

Tunnacliffe, E., Chubb, J.R. 2020. What is a transcriptional burst? *Trends in Genetics* **36**: 288-297. DOI: https://dx.doi.org/10.1016/j.tig.2020.01.003, PMID: 32035656

Gupta, A., Martin-Rufino, J.D., Jones, T.R., Subramanian, V., Qiu, X., Grody, E.I., Bloemendal, A., Weng, C., Niu, S.Y., Min, K.H., Mehta, A., Zhang, K., Siraj, L., Al' Khafaji, A., Sankaran, V.G., Raychaudhuri, S., Cleary, B., Grossman, S., Lander, E.S. 2022. Inferring gene regulation from stochastic transcriptional variation across single cells at steady state. *Proceedings of the National Academy of Sciences* **119**: e2207392119. DOI: https://dx.doi.org/10.1073/pnas.2207392119, PMID: 35969771
